# The dirty north: Evidence for multiple colonisations and Wolbachia infections shaping the genetic structure of the widespread butterfly *Polyommatus icarus* in the British Isles

**DOI:** 10.1101/2020.09.03.267203

**Authors:** Saad Arif, Michael Gerth, William G. Hone-Millard, Maria D. S. Nunes, Leonardo Dapporto, Timothy G. Shreeve

**Author notes:** Author for Correspondence Correspondence: **Saad Arif**, Centre for Functional Genomics, Oxford Brookes University, OX3 0BP Oxford, United Kingdom Department of Biological and Medical Sciences, Oxford Brookes University, OX3 0BP, United Kingdom.

## Abstract

The paradigm of differentiation in southern refugia during glacial periods followed by expansions during interglacials, producing limited genetic diversity and population sub-division in northern areas, dominates European phylogeography. However, the existence of complex structured populations in formerly glaciated areas, and on islands connected to mainland areas during glacial maxima, call for alternative explanations. Here, we reconstruct the mtDNA phylogeography of the widespread *Polyommatus icarus* butterfly over its native range, with an emphasis on the formerly glaciated and connected British Isles. We found distinct geographical structuring of *CO*1 mitotypes, with an ancient lineage restricted to the marginal European areas, including Northern Scotland and Outer Hebrides. We detected perfect mtDNA-*Wolbachia* associations in Northern Britain that support the possibility of at least two post-glacial *Wolbachia*-mediated sweeps, suggesting a series of sequential replacement of mtDNA in the British Isles and potentially in Europe. Population genomic analysis, using ddRADSeq genomic markers, also reveal unexpected genetic structuring within Britain. However, weak mito-nuclear concordance suggests the potential for independent histories of nuclear *versus* mitochondrial genomes. We found clustering of genomic SNPs of French samples, with respect to those in the British Isles, is not consistent with a scenario of a single recolonisation. Taken together our mtDNA and ddRADseq observations are consistent with a history of at least two distinct colonisations, a phylogeographic scenario previously put forth to explain diversity and structuring in other British flora and fauna. Additionally, we also present preliminary evidence that *Wolbachia-*induced feminization may be occurring in the isolated population in the Outer Hebrides.

## Introduction

Genetic differentiation among populations is the basis of evolution and speciation. Genetic differentiation typically emerges among allopatric populations after long-term geographic isolation. Due to changes in vicariance over geological timescales, current physical and ecological islands are not necessarily the same areas where diversification occurred. The Pleistocene period (2.6 mya (million years ago) – 11.7 kya (thousand years ago)) is characterised by a series of glacial/interglacial events. From the Last Glacial Maximum (22 kya) to *c*. 11.7 kya most of northern and central Europe was covered by ice caps, as well as the Alps and Pyrenees (Ehlers, Ehlers, Gibbard, & Hughes, 2011) and lowered sea levels connected many islands to the mainland and to each other (Hewitt, 1999). The end of the Pleistocene was characterised by a short period of rapid warming (17.5-12.8 kya) followed by cooling and glacial re-advance in the Younger Dryas (12.8–11.5 kya) before the current (but variable) warm period. Most European phylogeography is rooted in events during and following the last glacial period and there are many studies showing how diversification has emerged among the three southern European peninsulas and islands (Petit *et al*., 2003; Dapporto *et al*., 2019; Schmitt, 2007; Seddon, Santucci, Reeve, & Hewitt, 2001; Michaux, Libois, & Filippucci, 2005; Fiera, Habel, Kunz, & Ulrich, 2016) most likely from restriction and differentiation within southern isolated refugia in long cold periods followed by northward expansion during warm periods, resulting in lower genetic diversity in colonised than refugial areas. In particular many species in northern European areas, such the British Isles, are hypothesized to have been colonized via a single post-glacial colonization event and are expected to exhibit lower genetic diversity and lack of complex genetic structuring other than that resulting from serial founder events (Dincă *et al*., 2021; Hewitt, 1999; Mutanen *et al*., 2012). However, the increasing availability of DNA sequences, mostly based on mitochondrial markers, has, in some cases, revealed significant genetic structuring in northern European areas. Where this has been observed, it has been explained as the product of post-glacial colonization from different populations having persisted in reduced cryptic refugia in central Europe (e.g. Provan & Bennett, 2008; Schmitt & Varga, 2012) or by bottleneck events followed by recent local adaptation accentuated by reduced dispersal in the presence of short sea straits (Tison *et al*., 2014). Many islands which were connected to the mainland and to each other or covered by ice during the last glacial maxima also show genetically divergent populations (Cesaroni, Lucarelli, Allori, Russo, & Sbordoni, 1994; Dapporto *et al*., 2017; Scalercio *et al*., 2020 Tison *et al*., 2014). In these cases, successive post-glacial waves of colonization, likely driven by selective sweeps, and hampered by narrow sea straits have been hypothesized and reconstructed (Dapporto & Bruschini, 2012; Dapporto, Bruschini, Dincă, Vila, & Dennis, 2012, Tison *et al*., 2014).

A major drawback of many phylogeography studies is that they rely solely on mitochondrial DNA markers - usually a 650 base pair (bp) fragment of the mitochondrial *CO1* gene (e.g. Dapporto *et al*., 2017, 2019; Dincă *et al*., 2015; Hebert, Penton, Burns, Janzen, & Hallwachs, 2004; Lohman *et al*., 2010; Mendoza *et al*., 2016; Smith, Woodley, Janzen, Hallwachs, & Hebert, 2006; Scalercio *et al*. 2020). Mitochondrial DNA (mtDNA) markers can follow an evolutionary trajectory independent of the nuclear DNA. Due to the haploid nature and largely uniparental inheritance of the mtDNA, it has a fourfold lower effective population size compared to the nuclear genome. This lower effective population size means mtDNA loses genetic diversity *via* genetic drift at a faster rate than the nuclear genome (Charlesworth, 2009). Hence, mtDNA usually differentiates faster than the nuclear genome (Allio, Donega, Galtier, & Nabholz, 2017) during periods of isolation and will complete the process of lineage sorting more rapidly than the nuclear counterpart (Funk & Omland, 2003).

During range expansions (e.g. during interglacial periods) genetically differentiated lineages can meet, and in the absence of reproductive isolation, nuclear genomes can recombine and homogenize (e.g. Hinojosa *et al*., 2019). Discordance between mtDNA and nuclear genomic variation can result from introgression of mtDNA or sex-biased asymmetries, such as sex-biased dispersal (Toews & Brelsford, 2012; Dinca et al 2021). Insects in particular are also prone to infections by reproduction manipulating endosymbionts (e.g. *Wolbachia*) that can lead to cytoplasmic incompatibilities allowing mtDNA haplotypes (or mitotypes) to hitchhike to fixation without concomitant nuclear differentiation (Hurst & Jiggins, 2005). Hence, mtDNA variation by itself may provide an incomplete demographic history and should be complemented with neutral nuclear genetic markers in order to more accurately describe key demographic events, facilitate phylogeographic interpretations and delineate independent evolutionary lineages (Edwards, Potter, Schmitt, Bragg, & Moritz, 2016; Galtier, Nabholz, Glémin, & Hurst, 2009).

Endosymbionts like *Wolbachia are well known for their ability to induce cytoplasmic incompatability 9CI) but* may *also* alter host reproduction in other ways including male-killing (MK) and physiological feminization of genetic males (Makepeace & Gill, 2016). Although MK has been recorded in several instances in Lepidoptera and other insect groups, feminization has been observed much less frequently. Such reproduction manipulation strategies can have profound influence on host ecology and evolution (Drew, Frost, & Hurst, 2019). For example, bidirectional CI which leads to break down in reproduction between hosts harbouring different strains is expected to promote genetic divergence and potentially even speciation (Brucker & Bordenstein, 2012).

To understand how genetic structuring can emerge in formerly connected and glaciated areas, we focused on the British Isles, the largest European island system, which were connected to the European mainland until *c*. 8.0 kya but were covered by an ice cap up to 18.0 kya to the latitude of 51-53N degrees (Gibbard & Clark, 2011) with tundra and permafrost during the Younger Dryas period. The colonization of British islands by insects has been mostly dated from *c*. 13-10 kya (Atkinson, Briffa, & Coope, 1987; Coard & Chamberlain, 2016). We selected the Palearctic butterfly *Polyommatus Icarus* as a model species because of its abundance and widespread distribution. This species occupies a range of open biotopes including grasslands, sand dune systems and waste sites over a range of elevational gradients but is a host-plant specialist, with larvae feeding on low growing Fabaceae (chiefly *Lotus corniculatus*).

Phylogeographic analysis of this species across continental Europe and Asia recovered five divergent *CO1* lineages (Palaearctic, Iberia-Italy, Sierra-Nevada, Alicante-Provence and Crete), resulting presumably from multiple expansion/contraction cycles during the Pleistocene (Dincă, Dapporto, & Vila, 2011). More specifically, Bayesian divergence dating and ancestral range reconstruction suggests the existence of Palaearctic and southern European lineages *ca*. 1.8 mya. More recently (*ca*. 0.5 mya) there was an expansion of the Palearctic lineage into southern European refugia followed by divergence into a northern (Palaearctic) and southern European (Iberia-Italy) lineage. The expansion of the latter is concomitant with the range contraction and continued divergence of the ancient southern lineage into highly endemic and isolated lineages in the Sierra-Nevada, Alicante-Provence and Crete.

The phylogeography of this species in the UK remains unknown since no specimens from British islands were analysed by Dincă *et al*.’s (2011). However, Dincă *et al*.’s (2011) dating and range reconstruction on continental Europe suggests that the colonization of the British Isles likely consisted of a single or potentially two lineages (Palaeractic and/or Iberia-Italy). Additionally, an allozyme analysis suggested populations in the British Isles may have undergone a bottleneck (de Keyser, Shreeve, Breuker, Hails, & Schmitt, 2012), likely during the Younger Dryas period following colonisation in the early Holocene. However, It has also been suggested that the colonization of the British Isles by *P. icarus* could have involved more than one period of establishment following the Last Glacial Maximum (Dennis, 1977). Physiological differences (Howe, Bryant, & Shreeve, 2007) have been identified between populations in different parts of the British Isles, with Outer Hebrides populations flying with lower thoracic temperatures than southern populations. Additionally, modelling flight activity responses under climate change scenarios predicts differences in response to climate change between Outer Hebridean and mainland populations (Howe *et al*., 2007). Differences in life-history strategies also exist, with northern populations being (potentially obligate) univoltine (one brood of offspring annually) whilst southern ones are facultative polyvotine (> two to three broods of offspring annually)(de Keyser, 2012). There is thus potential for British islands to host genetically structured populations of *P. icarus* despite the relatively recent colonization.

Here, we describe comprehensively *P. icarus* mtDNA diversity and distribution in the British Isles and across the species’ entire native range. Focussing on the British Isles we then use genome-wide ddRADseq genetic markers to determine concordance with mtDNA to infer the potential colonization history of the British Isles. We also leveraged the ddRADseq data to conduct a survey of *Wolbachia* infection in *P. icarus*, in the British Isles, to integrate any influence of *Wolbachia* sweeps on our phylogeographic interpretations. We compare British Isles with European mtDNA data to infer possible invasion sequences into the British Isles. We compare our findings with existing interpretations of the phylogeography of *P. icarus* throughout Europe and demonstrate that combining mtDNA sequence data with nuclear genetic markers derived from genome-wide ddRADseq data and *Wolbachia* sequence data can provide comprehensive data for phylogeographic inferences. In addition, we also provide some preliminary evidence for the phenotypic effects of *Wolbachia* in the northern populations of the British Isles, including the possibility of a rare case of feminization.

## Materials and Methods

### Sample collection and CO1 sequencing

We sampled 190 butterflies from 14 sites spread across the British Isles together with a single site in central-southern France (Table S1, Figure S1) to serve as a reference out-group. Numbers of individuals collected per site varied between 6-15, with an average of 13. We aimed to collect similar numbers of males and females from each site, but our samples are male biased, due to cryptic female behaviour. Butterflies were sexed based on wing colouring and pattern dimorphism and abdominal tip morphology. We removed heads and legs of individuals anesthetized on ice and stored these in 95% ethanol for DNA extraction. Wings and bodies were dried and stored separately as specimen vouchers.

We sequenced a 655 bp fragment of *cytochrome c oxidase subunit 1* (*CO1*) for a subset of 140 individuals (Table S2). The fragment was amplified by PCR using primers piLepF1 (5“-TCTACAAATCATAAAGATATTGGAAC-3”) and LepR1 (5“-TAAACTTCTGGATGTCCAAAAAAATCA-3”) (Hebert *et al*., 2004) using OneTaq Mastermix with standard buffer (New England Biolabs) under standard cycling conditions. The resulting sequences were trimmed for primers and quality and then aligned using AliView (Larsson, 2014).

### Reconstruction of CO1 haplogroups in Europe and the British Isles

To determine the phylogenetic relationships of British *P. icarus* with those elsewhere in Europe (Dincă *et al*., 2011) we used our newly generated *CO1* sequences and publicly available *P. icarus CO1* DNA sequences from Europe and Eurasia archived in the Barcode of Life Data Systems (Ratnasingham & Herbert, 2007) and NCBI’s GenBank database. Sequences were trimmed for primers and quality and then aligned using AliView (Larsson, 2014) and truncated or potential contaminant sequences were removed, resulting in a final alignment of 585 specimens with length between 610-658 bp (Table S2). We constructed mitochondrial haplotype networks using the *CO1* alignments and TCS networks as implemented in *TCS 1.21* (Clement, Posada, & Crandall, 2000) by imposing a 95% connection limit (11 steps). Different haplogroups have been identified by creating a UPGMA dendrogram based on p-distances and calculated with the “dist.dna” function of the “ape” package (Paradis & Schliep, 2019). Hierarchical clustering was performed using the hclust function in R v.3.6.2 (R Core Team, 2019). Following the Dinca *et al*. (2011) assessment, we cut the tree at the depth of the fourth node using the “cutree” function to obtain 5 groups (based on Dincă *et al*., (2011)). The geographic distributions of these groups are visualised on a map using pie charts, with each group assigned a specific colour.

### ddRADseq library construction, sequencing, and SNP filtering

DNA was extracted from head and legs of all 190 individual butterflies using a salt extraction protocol (Miller, Dykes, & Polesky, 1988) and eluted in 60 μl of dH_2_0. DNA was quantified using a Qubit 2.0 flourometer (Life Technologies) using a Qubit dsDNA high sensitivity assay kit (Life Technologies) and individual DNA samples for ddRADseq libraries were normalized to10 ng/μl. Library preparation was performed at Floragenex (Portland, Oregon) following a protocol similar to Han *et al*. (2018) using a digestion with PstI/MseI along with a SBG 100-Kit v2.0 (Keygene N.V., Wageningen, the Netherlands). Barcoded samples were sequenced over two lanes at the University of Oregon Genomics and Cell Characterization Facility (Eugene, Oregon) on a HiSeq4000 with single-end 100bp chemistry.

We used Stacks 2.4 (Catchen, Hohenlohe, Bassham, Amores, & Cresko, 2013) to assemble RAD loci and call SNP genotypes from the raw ddRADseq data. Raw reads were demultiplexed using the process_radtags.pl script while discarding low quality reads (< 10 average Phred score) and removing any restriction site tags. After demultiplexing and discarding low quality reads, three individuals were removed from the final assembly due to low number of reads (<500,000; Table S3). We initially used a subset of 24 individuals to determine optimal combinations of the major parameters (*m* :minimum number of raw reads required to call a stack, *M*: number of mismatches allowed between stacks, *n*: number of mismatches allowed between loci of different individuals) involved in assembling RAD loci with Stacks following guidelines in Paris, Stevens, and Catchen (2017) and Rochette and Catchen (2017). We varied values of *m* from 2-12, while holding *M* and *n* constant at 2, and evaluated how the number of RAD loci and number of polymorphic loci present in 80% of the samples (r80 rule; Paris *et al*., 2017) stabilized as a function of m. After, obtaining a suitable value for *m*, we varied *M* and *n* from 1-8, with the constraint that *M*=*n* (Rochette & Catchen, 2017), and used the r80 rule and checked for stability of the proportion of loci with 1-5 SNPs to determine suitable values for *M* and *n*. The number of total and polymorphic RAD loci begin to stabilize around a value of 4 for all three major parameters (*m, M*, and *n;* Figure S2). The value of *M*=*n*=4 at *m*=4 was sufficient to stabilize the distribution of loci with 1-5 SNPS (Figure S3) and was used to assemble the final set of RAD loci. The average number of reads per individual was 2.95 million (standard deviation (sd): 1.09 million; Table S3). The average coverage per locus after assembling stacks was 45.8x (sd: 14.57x). SNP markers with minimum allele frequencies of < 0.05 and a maximum observed heterozygosity > 0.65 (to exclude potential paralogues) were further excluded. To obtain a set of widely available loci for downstream population genomic analysis, we only retained SNPs that were present in at least 50% (*r*-50) of the individuals within the 15 sampled localities (*p*-15) which yielded 4852 loci, 1915 of which were monomorphic.

To remove any contamination of our ddRADseq markers with mitochondrial DNA we used Centrifuge v.1.0.4 (Kim, Song, Breitwieser, & Salzberg, 2016) to search all RAD loci against NCBI’s database of mitochondrial RefSeq Genomes (https://www.ncbi.nlm.nih.gov/genome/organelle/), which includes complete mitogenomes of several Lepidopteran species. We also excluded contamination from *Wolbachia* using Centrifuge to search RAD loci against the Archaeal and Bacterial RefSeq Genomes in the NCBI database (ftp://ftp.ccb.jhu.edu/pub/infphilo/centrifuge/data/p_compressed_2018_4_15.tar.gz).

Finally, we used vcftools v0.1.17 (Danecek *et al*., 2011) to further exclude any loci with greater than 5 SNPs (potentially erroneous loci) and then generated two SNP data sets filtered on levels of missing data per individual: one that excluded individuals with > 50% missing data (*p*15*r*50miss50) and another excluding individuals with > 25% missing data (*p*15*r*50miss25). This resulted in two biallelic SNP datasets with 2,824 loci and 5,592 SNPs. The dataset *p*15*r*50miss50 consisted of 176 individuals and the dataset *p*15*r*50miss25 had 148 individuals. We generated two additional marker datasets, using the exact same procedure as above, but with a more stringent requirement to only retain SNPS that were present in at least 60% (*p*15*r*60miss25) or 70 % (*p*15*r*70miss25) of the individuals within each locality.

### Population structure based on putatively neutral and outlier ddRADseq markers

To examine genomic level population structure we conducted Principal Component Analysis (PCA) on the Stacks derived and filtered SNP datasets using the package ade4 v.1.7-13 (Dray & Dufour, 2007) in R v3.6.2 (R Core Team, 2019). To assess the impact of missing data in reconstructing population structure we performed PCAs on the two datasets filtered for individuals with different thresholds of missing data (*p*15*r*50miss50 or *p*15*r*50miss25). As linked SNPs can influence population clustering techniques like PCA we performed a further PCA on the *p*15*r*50miss25 dataset but retaining only a single SNP per RAD locus. Additionally, to assess the influence of varying number of RAD markers on population structure, we also conducted PCA on SNP datasets with more stringent inclusion criteria on the availability of loci (datasets *p*15*r*60miss25 or *p*15*r*70miss25).

To differentiate population structure arising from demographic and historical processes *versus* those potentially due to local adaptation or natural selection, we partitioned the *p*15*r*50miss25 SNPs into outlier and putatively neutral loci (Allendorf, Hohenlohe, & Luikart, 2010). We detected outlier loci using Bayescan v2.1 (Foll and Gaggiotti, 2008) and a maximum likelihood based approach as implemented in OutFLANK v. 0.2 (Whitlock & Lotterhos, 2015). Bayescan was run with default settings except that we used 1:100 prior odds and 100,000 iterations and a burn in of 50,000. We used a false discovery rate (FDR) of 1% as a cut-off for classifying a SNP as an outlier. OutFLANK was run with default settings and a false discovery rate of 5%, with an expectation of generating more conservative results (Whitlock & Lotterhos, 2015). The FRN (out-group) and RVS (southern Scotland, sample size < 5) samples were filtered while detecting outliers using either method. The union of the set of all SNPs detected by both in the *p*15*r*50miss25 were treated as outlier SNPs. The union of all loci associated with these outlier SNPs were removed from the *p*15*r*50miss25 dataset to generate a dataset of putatively neutral SNPs, thus generating sets of outlier and putatively neutral loci. PCAs were then performed individually for the outlier and putatively neutral SNP datasets.

We also calculated pairwise Weir and Cockerham *F*_st_ between all 15 populations, for both outlier and putatively neutral SNP datasets, using the R package dartR 1.1.11 (Gruber, Unmack, Berry, & Georges, 2018) on the *p*15*r*50miss25 dataset. Statistical significance between each pairwise *F*_st_ was determined using 10000 bootstrap replicates. We used fineRadstructure and RADpainter (Malinsky, Trucchi, Lawson, & Falush, 2018) to assess fine-scale population structure based on shared genetic co-ancestry using only the putatively neutral SNPs.

### Assessing concordance between mtDNA and genomic markers

We used an analysis of molecular variance (AMOVA) to assess the concordance between mtDNA variation and the putatively nuclear loci as derived from ddRADseq data. Individuals were assigned to groupings based on clustering of *CO1* sequences (as above) to assess how genomic variation partitioned based on mtDNA haplogroup. Strong concordance between mtDNA and genomic variation would be supportive of a hypothesis of multiple discrete colonisation events. AMOVA was performed on the *p*15*r*50miss25SNPs data using the R package poppr v.2.8.3 (Kamvar, Tabima, & Grünwald, 2014). Samples from France were excluded for this analysis and statistical significance was assessed with 10,000 permutations.

### Predicting Wolbachia infection in individuals

To predict *Wolbachia* infection in each individual, we searched each read from demultiplexed individual fastq files (generated in the initial stages of the Stacks 2.4 pipeline using process_radtags) against the index of NCBI’s database of Archaeal and Bacterial RefSeq Genomes (ftp://ftp.ccb.jhu.edu/pub/infphilo/centrifuge/data/p_compressed_2018_4_15.tar.gz) using Centrifuge v.1.0.4 (Kim *et al*., 2016). We used Pavian (Breitwieser & Salzberg, 2019) to summarize results from Centrifuge. To quantify *Wolbachia* infection level in each individual we calculated the number of reads mapping to a single (most common) *Wolbachia* strain as a fraction of the total reads mapped to any bacterial or archaeal genome. Previous predictions of *Wolbachia* infection status based on short read data from whole genome shotgun libraries have been highly successful (Richardson *et al*., 2012; 98.8% concordant with PCR-based results). Additionally, Illumina read depth have been shown to be a reliable proxy for *Wolbachia* titre, showing strong correlation with copy number estimated from quantitative PCR (Early & Clark, 2013). Differences in proportions of infected individuals between localities were determined using a Fisher’s Exact test with Bonferroni adjustment for multiple comparisons using base R v3.6.2.

### Generating ddRADseq SNPs for Wolbachia

In order to generate *Wolbachia* genotypes for the subset of infected individuals we used seqtk 1.3-r106 (https://github.com/lh3/seqtk) to retain only reads tagged by Centrifuge as mapping to the most common *Wolbachia* taxon (endosymbiont of *Drosophila simulans, w*No; NCBI txid: 77038). This set of filtered reads for the infected individuals was processed through the same final Stacks 2.4 pipeline as above (except the initial run of process_tags.pl) with parameters *m*:4-*M*:4-*n*:4 and excluding genotypes with allele frequencies of < 0.05, a maximum observed heterozygosity > 0.65. Additionally, the SNPs had to present in at least 50% of individuals predicted as infected (*r*-50) in at least half of the locations harbouring infected individuals (*p*-3). This resulted in 124 loci, 86 of which were monomorphic. BLAST searches revealed that 117 of the 124 loci were 100% identical to at least one *Wolbachia* genome assembly from NCBI, and a further 5 were >=98.5% identical to *Wolbachia* sequences in NCBI Genbank. BLAST searches against NCBI GenBank further showed that the remaining 2 loci best matched *Wolbachia* sequences but identities were low (94.7% and 92.6%); however, these loci harboured no variants and hence were not used in any downstream analyses. The closest matching *Wolbachia* genomes for all loci were from supergroup B strains and 115 of the 124 loci could be assigned to a *Wolbachia* protein with sequence identities of >=95%. Matching loci were mostly housekeeping genes and showed an enrichment for hypothetical proteins. Loci carrying SNPs (38) were further filtered to remove any loci with >5 SNPs and we then generated two SNP marker datasets for downstream analysis, one in which individuals with >50% missing data were removed and another that included all individuals.

### Testing association between Wolbachia strains and mitotypes

To assess the congruence between *Wolbachia* genotypes and *P. icarus* mitochondrial haplotypes, we independently clustered CO1 sequences of infected individuals and concatenated SNPs from *Wolbachia* genotypes derived from the Stacks pipeline (two datasets: one with >50% missing data individual exclusion criteria and the other without). Both sets of data were clustered independently using bitwise distance (or Hamming’s distance) with UPGMA and 1000 bootstrap replicates for support using the R package poppr v.2.8.3 (Kamvar *et al*., 2014) and clustering dendograms were visualized and annotated using the R package ggtree v.1.17.4

### Identifying sex-Linked loci to Investigate Wolbachia Induced feminization

To investigate potential feminization in *Wolbachia* infected females we sought to identify sex-linked SNPs to establish the genetic sex for each individual. Butterflies generally possess a chromosomal ZW/ZZ sex determination mechanism where females are the heterogametic (ZW or ZO) sex (Traut, Sahara, & Marec, 2007). Hence, female-specific sex markers (loci polymorphic in females but homozygous in males), assuming partial homology between Z and W chromosomes, should help to determine the genetic sex of an individual butterfly: discordance between genetic and morphological sex of infected females would be consistent with physiological feminization. To identify female-specific sex markers we first fitted a baseline generalized linear model (using a logit link function) with morphological sex as the dependent variable and PC1 and PC2 from the PCA of SNPs as the independent variables. We then fitted an additional model by adding a single SNP marker as an additional independent variable and iterated this for all SNPs in the dataset. A significant association between a SNP marker and morphological sex was determined by performing a likelihood ratio test of the baseline model and the baseline model with the added SNP term and using a strict Bonferroni corrected 5% type I error rate. Significant markers were further excluded if they had more than 2 genotypic classes, were not homozygous for males or happened to be a single SNP from a RAD locus harbouring multiple SNPs. Generalized linear models and likelihood ratio tests were performed using the *glm() and anova()* functions in base R v3.6.2.

## Results

### Geographic distribution of CO1 haplogroups in Europe and the British Isles

An UPGMA clustering of 585 CO1 sequences (Figure S4) identified the same five main haplogroups as identified by Dincă *et al*. (2011). Specimens belonging to these five groups are highlighted in a TCS haplotype network (Figure 1A) with the Crete lineage (red) being the most divergent haplogroup. The Sierra Nevada lineage (yellow) was limited to this geographic region but a second specimen, differentiated by two mutations, has been found in Austria (Figure 1B). The second lineage (Alicante-Provence, purple) limited to Iberia and France according to Dincă *et al*. (2011), was also found in Norway, Germany and the British Isles. Most specimens from Southern-Western Europe belong to a different haplogroup (Iberian-Italian lineage, orange in Figure 1) compared to most specimens form Central-Eastern Europe and the Middle East (Palaearctic group, green in Figure 1). When data for islands and their closest mainland (or larger island) is available, islands always showed higher incidence of haplogroups identified as having expanded from refugia early in the Holocene by Dincă *et al*. (2011). Moreover, in central Europe, which is almost completely inhabited by the Palaearctic haplogroup (green) some, possibly relict, haplogroups occur (arrows in Figure 1B).

**Figure 1.**
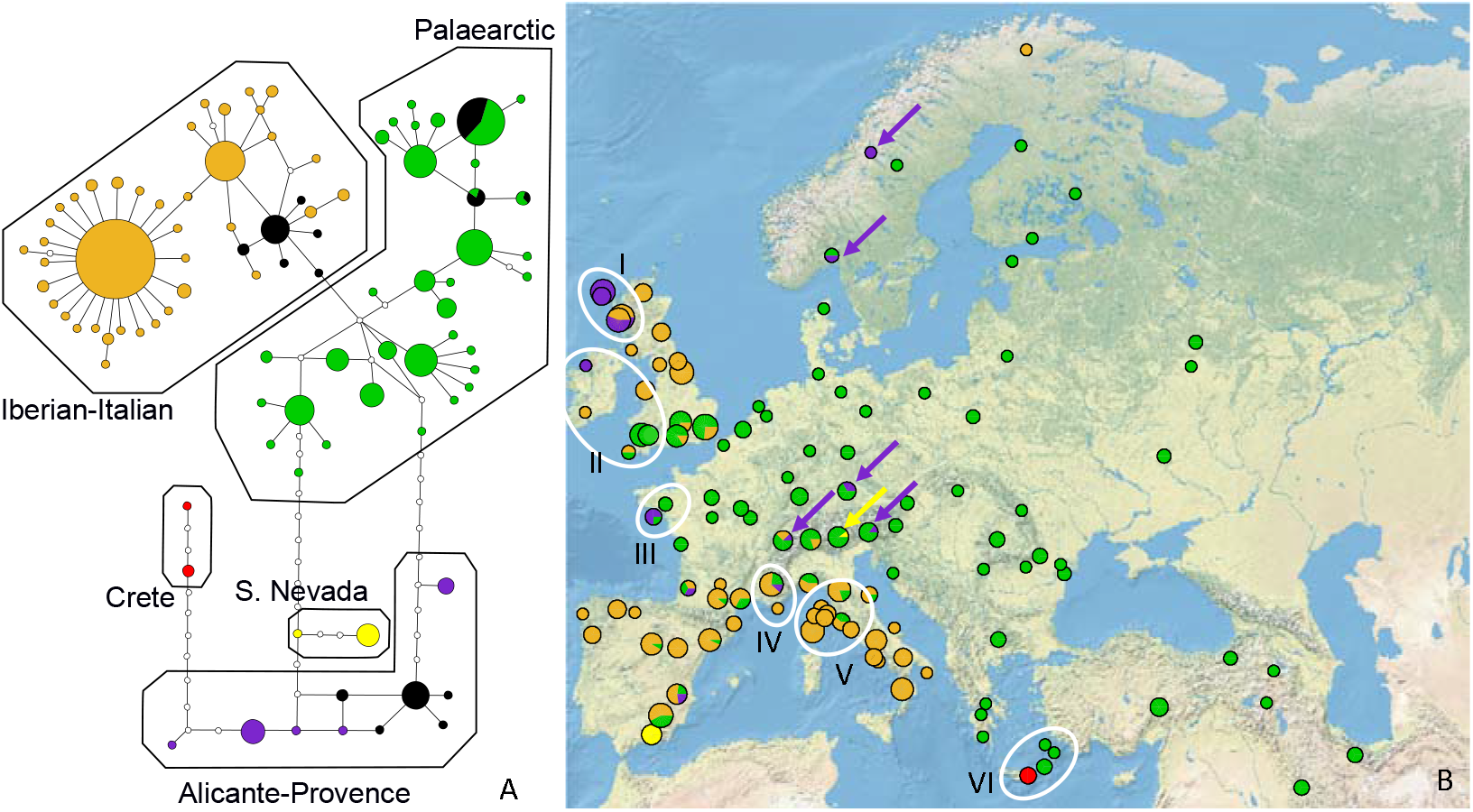
The haplotype network based on COI and divided in five main haplogroups according to a division in a UPGMA clustering which aligns to previous assessments **(A)**. Specimens in black belong to British islands. The collection sites of the specimens included in the haplotype network (same colours as in A) are also mapped **(B)**. Specimens are grouped in pie charts for squares of 2×2 degrees of latitude-longitude and circle area is proportional to the number of specimens. The systems of island-mainland (or larger island) are indicated with roman numbers (I, Hebrides-Britain; II, Ireland-Britain; III, Belle-Île-en-Mer-France mainland; IV, Levant island-France mainland; V, Tyrrhenian islands-Italian mainland; VI Crete-neighbouring islands-Greek and Turkish mainland. Many specimens with haplotypes regarded as relict from past colonization waves are also found in Central Europe and Middle East and highlighted with arrows.

Samples from the British Isles did not belong to a single haplogroup (black sectors in Figure 1A), but exhibited strong geographic clustering (Figure 1B, magnified in Figure S5). Those from the Outer Hebrides together with some from the adjacent Scottish mainland were part of the Alicante-Provence lineage. Those from southwestern and southern parts of the British mainland and Wales were part of the main Palaearctic group, whilst those from central and northern parts of mainland Britain were part of the Iberian-Italian lineage.

### Population structure of British Isles P. icarus using genome-wide ddRADseq SNPs

The first component, accounting for 6.8% of the variation and which potentially corresponds to latitude, separates the northern Scottish samples (BER, TUL, MLG, DGC, and OBN) from all the southern, southwestern and Welsh samples (Figure 2A). Individuals from RHD (northern England) and RVS (southern Scotland) samples fell between these extremes along the first component. Surprisingly, the FRN (French) samples also fell in between the extremes of PC1. The second component, accounting for 1.7% of the variation, distinguishes the Outer Hebrides (BER, TUL) from the northern Scottish mainland locations (MLG, DGC, OBN) (Figure 2A).

**Figure 2.**
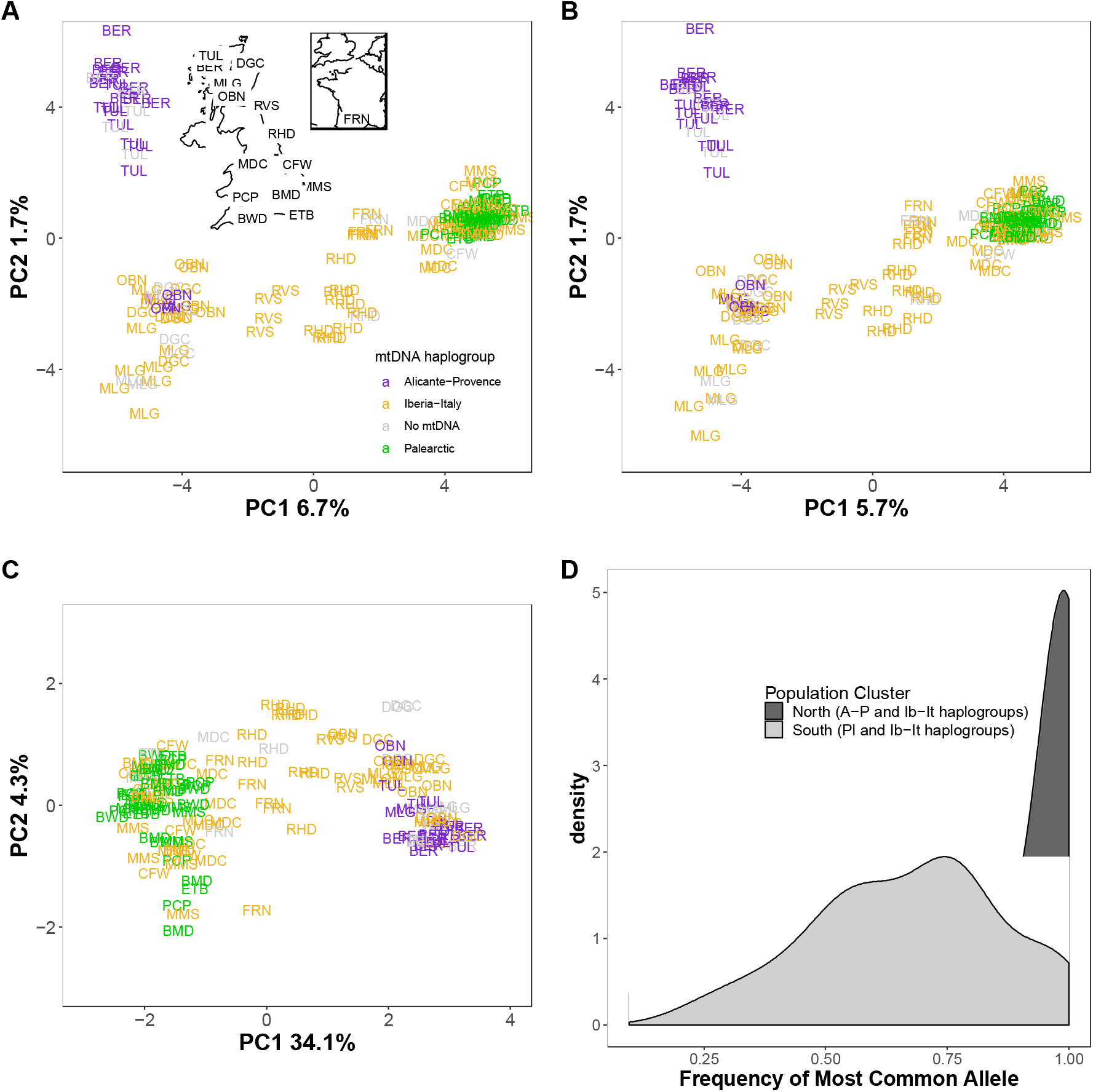
Population structure based on principal component analysis (PCA) of 5592 SNPs (2824 RAD loci) of 148 *Polyommatus icarus* individuals across the British Isles. PCAs are shown for the entire dataset **(A)**, a subset of 5387 putatively neutral SNPS **(B**), and a subset of 104 outlier SNPs (across 84 RAD loci) **(C). (A)** inset is a labelled map of the sampled localities for quick reference. Populations are coloured by their mtDNA haplogroup, where data is available, to help visualize concordance between mtDNA and genomic markers. (D) Density distributions of the frequency of the most common allele (MCA) for the 104 outlier SNPs across northern and southern British Isles. Individual sampling locations are aggregated by geographic clustering of localities along PCA 1 in **A-C (A-P: Alicante-Provence, Ib-It: Iberia-Italy, Pl: Palaearctic)**. Localities in intermediate positions (RHD, RVS) and reference outgroups (FRN) were excluded. North: TUL, BER, 13 DGC, MLG, OBN; South: MDC, CFW, MMS, PCP, BMD, ETB.

Results of the PCA were largely invariant to missingness or marker number. PCA analyses of the ddRADseq SNP data filtered for individuals with either >25% (*p*15*r*50miss25) or >50% (*p*15*r*50miss50) missing data and with the dataset filtered for 25% missing data (*p*15*r*50miss25) but using unlinked SNPs all produced virtually identical results (Figure 2A and Figure S6A-B). Datasets with smaller number of markers based on more stringent inclusion criteria (datasets (*p*15*r*60miss25 and (*p*15*r*70miss25) also yielded similar results but the signal decayed with decreasing number of markers (Figure S6C-D).

Next, we partitioned *p*15*r*50miss25 into outlier and putatively neutral loci to disentangle population structure arsing potentially from natural selection *versus* that from historical demographic processes.

Together the two outlier detection methods recovered 104 SNPs (Bayescan: 103, OutFLANK: 10) across 84 ddRADseq loci. These 84 loci including all SNPs (even those not deemed as outliers) were further filtered to produce set of putatively neutral SNPs from the *p*15*r*50miss25 data set. This putatively neutral SNP dataset had 5387 SNPS across 2740 loci for 148 individuals and was, unless otherwise stated, the primary SNPs data set for the genomic analyses that follow. Population structure based on PCA of the neutral SNPs (Figure 2B) recovers the same topology as that using the full set of SNPs (Figure 2A), except the variation explained by PC1 is reduced to 5.7%. However, the PCA for outlier SNPs (Figure 2C) only exhibits clustering between English/Welsh and Scottish samples, with RVS, RHD, and FRN again falling in between the two extremes. PC1 explains 34.1 % of the variation in the outlier SNP dataset, while PC2 explains 4.3%, although no obvious stratification is apparent in the second component. To better understand the variation in the outlier SNPs between the northern and southern clusters in the PCA (Figures 2A-C), we plotted the frequency of the most common allele (MCA) for all 104 outlier SNPs (Figure 2D). In general, northern populations showed higher frequencies and mostly fixation of the MCA compared to southern populations.

There was significant structuring based on neutral Pairwise *F*_st_ values among locations, across the British Isles, other than those in south (Figure S7A). Pairwise *F*_st_ values for neutral markers between northern Scottish and all southern populations suggest moderate levels of differentiation (0.075-0.109; Figure S7A), while those between Outer Hebrides and the northern Scottish mainland locations suggest small, yet significant, levels of differentiation (0.047-0.064, Figure S7A). Differences in Pairwise *F*_st_ values based on outlier SNPs *versus* those on neutral markers (Figures S7A-B mirrored the differences in the PCAs for neutral and outlier markers (Figures 2B-C). Pairwise *F*_st_ values for outlier SNPs were extremely high between northern Scottish and southern populations (0.385-0.563 Figure S7B).

Analysis of shared genetic co-ancestry using fineRADstructure also presents clustering of individuals (Figure S8) qualitatively similar to those from the PCA of entire and neutral markers only data sets. The resulting clustered co-ancestry matrix (Figure S8) revealed three distinct clusters consisting of the (i) northern Scottish samples, (ii) southern and central Great British samples, and (iii) the 6 individuals from southern France. There was evidence of further substructure in the northern Scottish population with the Outer Hebrides (BER, TUL) samples forming their own distinct sub-cluster within the northern Scottish cluster (Figure S8). Individuals from RVS and RHD clustered together with the French population sharing genetic ancestry with both the northern mainland Scottish and southern British populations.

### Concordance between mtDNA and Genomic Markers

If the British Isles had been populated by three discrete colonization and establishment events, as implied by the geographic structuring of mtDNA haplogroups in the British Isles (Figure 1, Figure S5), we would expect strong concordance between mtDNA and genomic markers. Some association is evident between mtDNA and genomic markers (Figure 2A-C) and we evaluated this relationship in an AMOVA framework. The association of mtDNA haplogroups with genomic variation was weak (4.8%) but significant (*p*-value < 0.0001) while genomic divergence within the haplogroups was stronger (15.6%, *p*-value < 0.0001).

### Prediction of Wolbachia infection

Using Centrifuge to match all demultiplexed reads (for all 190 individuals) to NCBI’s RefSeq genomes of archaeal and bacterial genomes, we identified an average 5.07 % (standard deviation: ±1.96%) of the total reads per individual were classified as matching an archaeal or bacterial genome (Table S4) with most matches being to bacterial genomes (5.05 ± 1.96 %; Table S4). The most frequently encountered *Wolbachia* genome was identified as an endosymbiont of *Drosophila simulans, w*No (NCBI txid: 77038). The total percentage of classified reads with matches to this genome varied by several orders of magnitude across individuals (min= 0.00036%, max=53.56%, Table S4). There was no relationship between the total number of raw reads and the percentage of classified reads mapping to this taxon (*Spearman’s rank correlation* = 0.131, *P*= 0.07544; Figure S9A). There was a natural discontinuity in the percentage of classified reads mapping to *Wolbachia* across all individuals (Figure S9B) and this metric clearly had a bimodal distribution. Using a threshold of log_2_ (percentage of classified reads mapping to *Wolbachia*) > 0 to classify an individual as infected or not, only individuals from BER, TUL, DGC, MLG, OBN and one individual from RHD were classified as infected with infection percentages ranging from 87.5% (14 of 16 individuals), in TUL in the Outer Hebrides, to 8% (1 of 13) in RHD near Durham northern mainland Britain) (Figure 3A). The proportion of infected populations differed significantly between RHD and all other localities; however, all other pairwise comparisons were insignificant (Table S5). All females from the Outer Hebrides (BER, TUL) were infected with a high percentage of classified reads mapping to *Wolbachia* (min=15.6%, max=53.56%; Figure 3B). However, no such dimorphism in number infected or percentage mapped reads was apparent in the mainland populations (DGC, MLG, OBN; Figure 3B).

**Figure 3.**
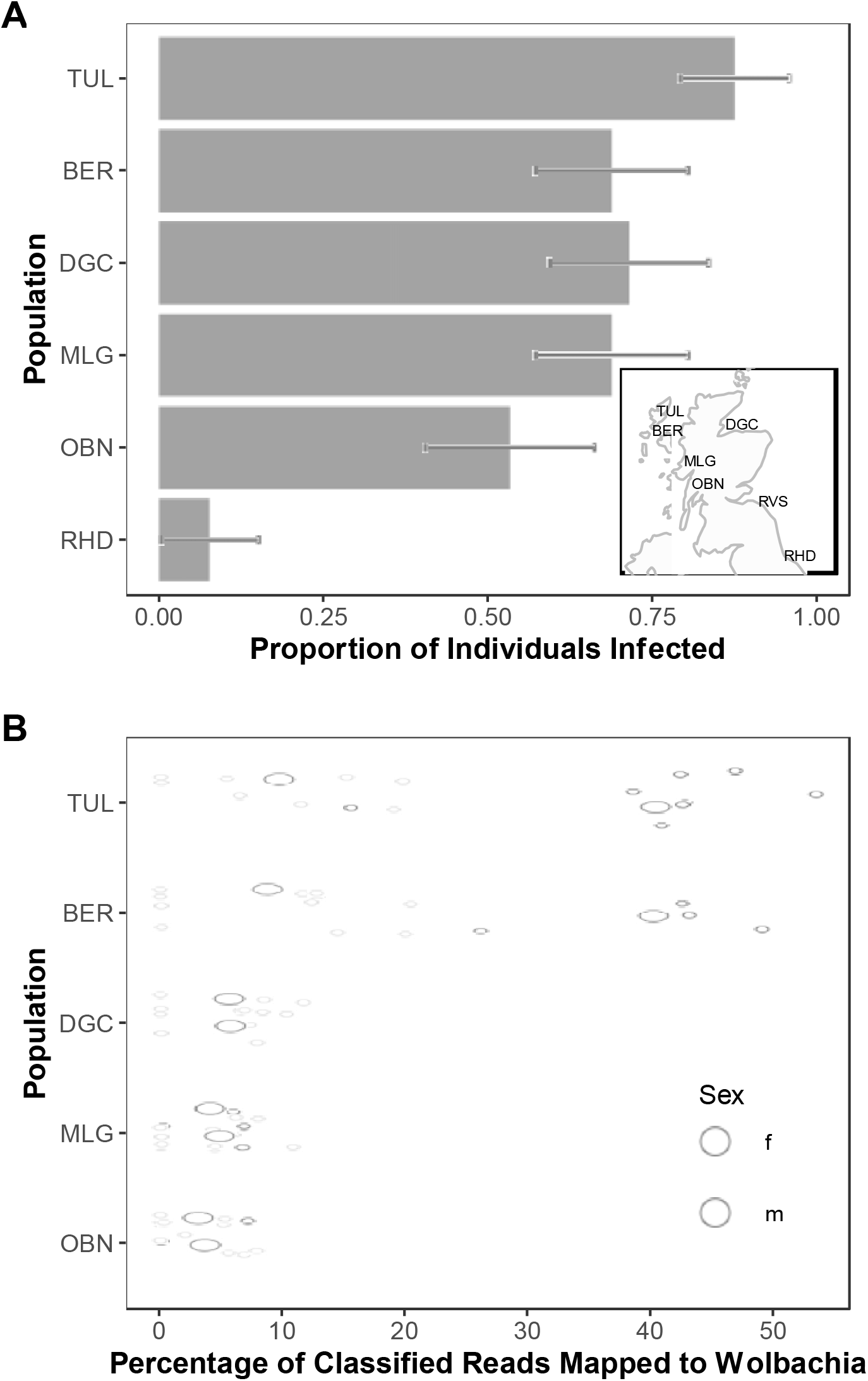
**(A)**. Proportion of individuals classified as infected based on a threshold of log_2_(percentage of classified reads mapping to Wolbachia) > 0. Only populations with at least one infected individual are shown. Error bars are the standard errors of the estimated proportions. Inset map spans the geographical range of locations harbouring infected individuals. **(B)**. Distributions of the percentage of classified reads mapping to *Wolbachia* for locations with individuals classified as infected. RHD with a single infected male is not shown. Larger circles represent the average for each sex within locations.

### Congruence between CO1 mitotypes and Wolbachia strains

To determine association between *Wolbachia* genotypes and *CO1* mitotypes, we performed UPGMA clustering of Hamming’s distance for each for *CO1* sequences (derived from of a subset of 38 infected individuals where sequence information was available) and *Wolbachia* ddRADseq SNPS (derived from all 55 infected individuals). For *CO1* sequences we recovered two clusters with strong support, corresponding to the Alicante-Sierra Nevada and Iberia-Italy CO1 haplogroups (Figure 4). The Alicante-Sierra Nevada cluster was composed entirely of individuals from the Outer Hebrides except for three individuals from the nearby western coast of mainland Scotland (MLGm002, MLGf010, & OBNm110; Figure 4). UPGMA clustering of Hamming’s distance between concatenated SNPs derived from the *Wolbachia* genotypes dataset with and without the >50% missing data exclusion criteria were near identical, only the latter is shown (Figure 4). Clustering of *Wolbachia* SNPs also recovered two strongly supported clusters, corresponding to strain *w*Ica1 (Outer Hebrides) and *w*Ica2 (mainland). The three individuals from the mainland (MLGm002, MLGf010, & OBNm110) possessing the Alicante-Sierra Nevada haplotype also carried the *w*Ica1 strains. There is perfect association between individuals bearing *Wolbachia* strain *w*Ica1 and the Alicante-Sierra Nevada CO1 haplotype and *Wolbachia* strain *w*Ica2 and the Italy-Iberia CO1 haplotype (although not all individuals with this haplotype are infected by *Wolbachia*) regardless of geographical locality.

**Figure 4.**
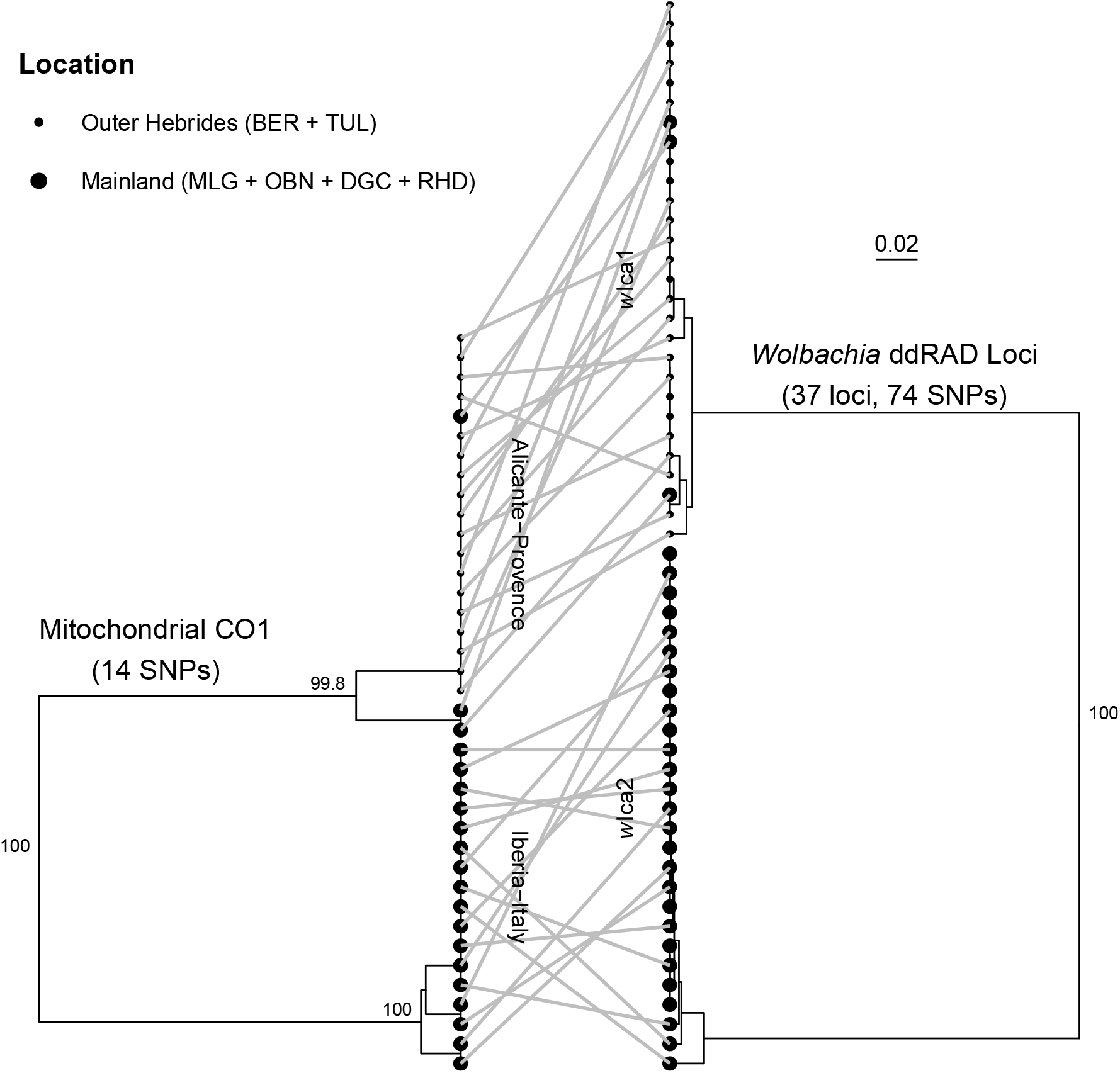
UPGMA clustering of mitochondrial CO1 fragments from 38 *Polyommatus icarus* individuals based on bitwise distance (left). UPGMA clustering of 74 concatenated SNPs from ddRADseq *Wolbachia* genotypes derived from reads mapping to *Wolbachia* from 55 infected individuals (right). Numbers on nodes represent bootstrap branch support values based on 1000 bootstrap replicates, values < 70% are not shown. Large circles represent individuals from mainland populations (DGC, MLG, OBN, RHD) and smaller circles represent individuals from the Outer Hebrides (BER, TUL). Lines between dendograms connect the *CO1* haplotype and *Wolbachia* strain for the same individual. Scale bar reflects the proportion of loci that are different.

### Sex-specific markers and feminization in Outer Hebrides

*Wolbachia* dosage can influence phenotypic outcomes in the host (Arai, Lin, Nakai, Kunimi, & Inoue, 2020; Breeuwer & Werren, 1993), thus the high levels of *w*Ica1 observed in females of the Outer Hebrides population could be indicative of *Wolbachia* induced feminization. In this scenario, morphological males should all be homozygous and morphological females should be heterozygous for female-specific markers (homozygous for Z but a novel allele for W chromosome). Discordance between the latter would suggest potential feminization of males. We used a combination of association analysis and filtering to identify such a set of 10 putatively female-specific SNPs (Table S6) across 7 RAD loci. Strikingly 5 out of 10 SNPs (across 3 RAD loci) showed perfect discordance of morphological sex with genetic sex except for females carrying the *w*Ica1 strain (Figure 5). The additional 5 SNPs also show complete discordance for morphological and genetic sex for *w*Ica1 infected females but also includes a small number of homozygote of females from uninfected or *w*Ica2 carrying individuals (Table S6).

**Figure 5.**
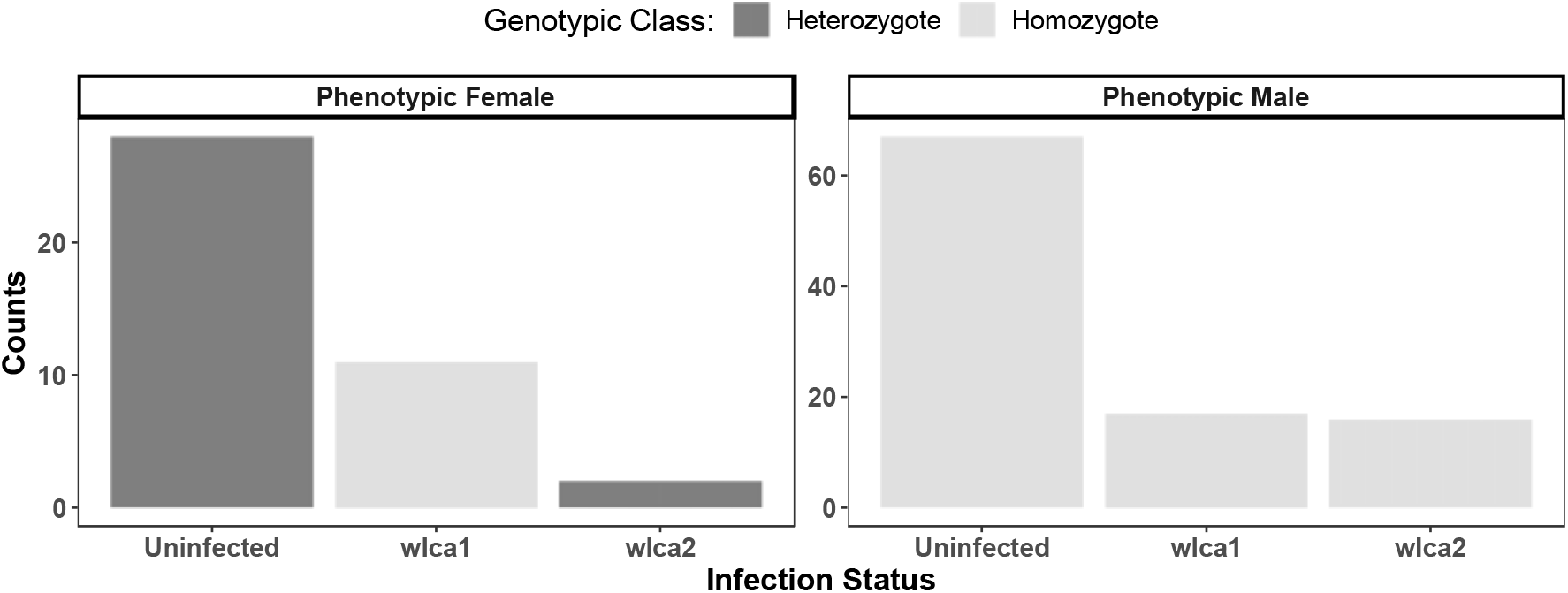
Feminization of *w*Ica1 infected females as suggested by discordance between morphological sex, based on abdominal tip morphology and dimorphic wing patterning, and genetic sex based on female-specific markers. Discordance is exemplified by data on a single marker here (11011_27), see Table S7 for all sex-linked loci. All *w*Ica1 infected morphological females (all females in the Outer Hebrides and one from the mainland (MLGf010) are homozygous for female-specific markers (heterozygous ZW), consistent with feminization of males.

## Discussion

In this study we used mtDNA and ddRADseq data to describe the genetic variability and structure of *P. icarus* within the British Isles and additionally to determine the historical processes underlying this contemporary genetic structure. We found strong geographical structuring in both mtDNA and the nuclear genome. However, there was only weak concordance between mtDNA and genomic variation at the nuclear level, suggesting the potential for partially independent evolutionary trajectories for the mitochondrial and nuclear genomes. Our results suggest that recurrent *Wolbachia*-mediated mtDNA sweeps can strongly contribute to the sorting of mtDNA haplogroups in the British Isles (and potentially in Europe). Moreover, co-ancestry of genomic clusters within the British Isles and the putative out-group samples from France raises the possibility of a distinct two-phase colonization that merits further investigation. Finally, we also present some preliminary evidence for potential *Wolbachia*-mediated feminization in an isolated population in the Outer Hebrides.

### Evidence for multiple mtDNA sweeps across Europe and the British Isles

Using mitochondrial *CO1* and the nuclear gene *ITS-1*, Dincă *et al*. (2011) identified five lineages with geographic structuring in southern Europe. Using the observed distribution of lineages and inferred divergence time, ranging from 1.8 mya to 0.5 mya, Dincă *et al*. (2011) reconstructed and dated to the Pleistocene a series of divergence and dispersal events followed by genetic sweeps which could have potentially produced the observed genetic structuring. The reconstruction predicted that Iberian and Crete lineages diverged in these areas around 1.8 mya ago, followed by the separation of an early Iberian lineage (1.2 mya) into the Sierra Nevada and the Alicante-Provence lineages. They would have then been replaced in most of the Iberian Peninsula by the Iberia-Italy haplogroup (500 kya), in turn replaced over central Europe by the Palaearctic one in the upper Pleistocene. By extending the geographical sampling to Northern Europe, we provide further evidence for the occurrence of similar waves of colonization also over the formerly glaciated areas of Europe. In fact, the pattern showing the more ancient colonizer appearing as a relict in the southern-most Mediterranean areas is completely reversed in Northern Europe, where the supposed ancient colonizer is limited to northern-most and marginal areas. The wider distribution of haplogroups on European islands is consistent with multiple waves of colonisation with recent evidence from islands and neighbouring mainland areas indicating that islands have a higher proportion of mitotypes from supposed earlier colonization events (based on dating from Dincă *et al*. (2011)). Finally, relict mitotypes occur throughout areas of central Europe representing potential invasion sequences, including the occurrence in Austria of a specimen of the Sierra Nevada haplogroup previously supposed to have evolved in southern Spain and possibly representing the first colonization wave.

Over the British Isles the three haplogroups also show a clear geographic stratification with the Alicante-Provence haplogroup restricted to the Outer Hebrides (with a single sample from northwestern Ireland); the northern mainland occupied by the Iberia-Italy haplogroup; and the southern English and Welsh populations being largely composed of the Palaearctic haplogroup. Additionally, the Alicante-Provence and Iberia-Italy haplogroups present in the British Isles are perfectly endemic (Figure 1A), whereas the Palaearctic haplogroup present in the British Isles are a sub-sample of the continental European mitotypes. This incomplete lineage sorting of Palaearctic mitotypes in the British Isles could be indicative of a more recent invasion. The persistence and structuring of Alicante-Provence and Iberia-Italy on the other hand could be explained by our finding of mitotype-specific *Wolbachia* association within the British Isles (further discussed below).

Our mtDNA data demonstrates all the main lineages except the Cretan one occur in the areas covered by ice sheets at the time of the Last Glacial Maximum (22 kya) which imposes a relatively short time limit to the waves of dispersal to northern Europe and the British Isles. The earliest time period that *P. icarus* could have occurred within the British Isles is *c*. 13 kya following rapid warming at the end of the Last Glacial Maximum. Most dating of insects arriving to British islands and persisting now are between the end of the Younger Dryas (11.8 kya) and the severance of the British Isles from mainland Europe by a permanent sea-straight c. 8 kya. Again, the islands currently showing genetic contrasts from adjacent mainland areas (Levant, Belle-Ile-en-Mer, some Tuscan islands, Hebrides, Ireland) were connected to the mainland in the Last Glacial Maximum, but for the northerly islands connections would mostly have been available during the period when both climate and vegetation was tundra. Thus, the sequential invasions would have occurred in the interglacial when the sea barriers were re-established and hampered genetic sweeps (Dapporto & Bruschini, 2012).

### Distinct genomic clusters within the British Isles

Adults of *P. icarus* are described as relatively mobile (Asher *et al*., 2001; Cowley *et al*., 2001) and early work at the European scale using allozymes revealed little genetic differentiation over mainland Europe, and no geographic regionalisation, although samples from the British Isles exhibited lower allelic diversity than in mainland Europe (de Keyser *et al*., 2012). Our data contrasts with this broad finding, demonstrating a pattern of geographic variation even over the smaller spatial scale of British Isles. Population structure and demographic inference based on putatively neutral genomic markers reveal substantial differentiation between northern (mostly Scottish) and southern populations (Figure 2B). There is further subdivision between the Outer Hebrides Islands and the mainland in Northern Scotland. The latter substructure in northern Scotland is relatively weak (1.7% compared to 5.6% in PCA between northern and southern populations) and could be a direct result of the geographical isolation of Outer Hebridean populations but could, theoretically, also result from *Wolbachia*-mediated bidirectional cytoplasmic incompatibility (see below).

There is stronger divergence between northern (Outer Hebridian plus Scottish Highlands) and southern (Central/Southern England and Wales) populations for both neutral and outlier markers (Figure 2B and C). Outlier loci detected based on tests for directional selection are often hypothesized to be loci involved in local adaptation and it is possibly that these markers may reflect or be linked to markers adapted to regional environmental conditions. However, the significance of this divergence, presumably resulting from both neutral demographic processes and directional selection, is not entirely obvious. There is evidence of switch from bivoltine life history to a univoltine life history along the Scottish-English borders (de Keyser, 2012) that could act as a barrier to gene flow but there is still overlap of flight period among reproductive adults (Matechou, Dennis, Freeman, & Brereton, 2014). However, an *in situ* barrier to gene flow does not explain the complete absence of genetic structure (Figures 2A-C, Figure S7-8) in southern locations compared to those in the north, and the high level of heterozygosity in outlier loci (Figure 2D) observed in this cluster.

An additional surprising observation was the relationship of the south-central French (FRN) samples to the northern and southern British clusters. The FRN samples were selected as a reference out-group that would be expected to be ancestral to all populations in the British Isles, under a model of a single colonization event. However, both the PCA analyses (Figure 2) and the fineRadStructure co-ancestry analyses (Figure S8) place the FRN samples as intermediate to northern and southern populations. Such an observation could result from an admixture in southern France. However, another possibility is that the northern and southern clusters in the British Isles results from two independent invasion and expansions events from an ancestor of the contemporary FRN population. This scenario would also imply that the southern British cluster results from the admixture of two different colonization sources, which would be consistent with the lack of genetic structure and higher heterozygosity of outlier loci (isolate breaking) in this population. This hypothesis requires further corroboration by a wider sampling of European specimens to infer potential ancestral populations.

### Impact of Wolbachia infection on demographic inference in the British Isles

Past and contemporary *Wolbachia* sweeps can confound reconstruction of phylogeographic dynamics for arthropods based upon mtDNA alone (Galtier *et al*., 2009; Hurst & Jiggins, 2005). Within the British Isles, populations in the north harboured a large proportion (>50% in all cases) of infected individuals, whereas no predictions of infection were made for individuals in the south suggesting that individuals in the south are not infected or infection levels are much lower. We found evidence for two distinct *Wolbachia* strains, *w*Ica1 and *w*Ica2, that show perfect association with the Alicante-Provence and Italy-Iberia haplogroups, respectively. Such genetic structuring of *Wolbachia* could potentially explain the persistence of these early colonising haplogroups of the host butterfly under male-biased dispersal and CI (Hurst & Jiggins, 2005) between infected males and uninfected females (and between males and females infected with different strains). Firstly, this could account for the observation that intermediate individuals in southern Scotland/northern England (RVS, RHD) and several southern populations (CFW, MDS, MMS, BMW) harbour the Iberian-Italian haplogroup. We detected *Wolbachia* infection (albeit as a single case) as far south as northern England (RHD).

Second, bidirectional CI between *w*Ica1 and *w*Ica2 individuals coupled with imperfect vertical transmission could promote stable coexistence of both strains (Telschow, Yamamura, & Werren, 2005) which could account for the persistence of the relict haplogroup in the Outer Hebrides. We did not detect any double-infected individuals (i.e., those carrying both strains), which would be consistent with bidirectional CI. However, it remains possible that are our sequencing data are not sensitive enough to detect double infections.

Existence of *Wolbachia*-mediated mtDNA sweeps in the British Isles also raise the possibility of their influence on mtDNA phylogeography in continental Europe. Recent C*O1* analyses reveal that many butterfly species exhibit diverse and complex mtDNA genealogies across Eurasia (Dincă *et al*., 2015). However, more comprehensive genomic analysis, that have only recently become available, show that differentiation in the nuclear genome is not always concordant with mtDNA variation (Dincă, Lee, Vila, & Mutanen, 2019; Hinojosa *et al*., 2019; Tóth *et al*., 2017) and the observed mito-nuclear discordance could be explained may be past or contemporary *Wolbachia* infections (Després, 2019; Gaunet *et al*., 2019). A systematic survey of infections on continental Europe (e.g. Sucháčková Bartoňová *et al*., 2021) and divergence dating of *Wolbachia* strains, could offer more conclusive insight into the impact of *Wolbachia* on genetic structuring in *P. icarus*. However, *Wolbachia* infections show high turnover (Bailly-Bechet *et al*., 2017) and determining the influence of past infections on host biogeography remains challenging.

### Evidence for a sequential colonization of the British Isles

Our mtDNA analysis was suggestive of three independent colonisations of the British Isles. We also detected three clusters based on our genomic data (Figure 2A-B) but there is only weak association between mtDNA haplogroups and genomic variation (AMOVA, 4.8%). However, we argue, that some of our observations are consistent with a scenario of two discrete colonization events. Most suggestive of these is the relationship of the French samples to northern and southern British genomic clusters as discussed above. The placement of these samples along the PCA (Figure 2A-B) and the co-ancestry matrix (Figure S8) suggest that the northern and southern British clusters are independent expansions most likely from an ancestor of the French population. The high heterozygosity and lack of genetic structure in the southern population could result from an admixture of a resident and a second recolonizing population. The second source population could be from a central European refugium, potentially representing the recent Palaearctic lineage expansion as reconstructed by Dincă *et al*. (2011). Despite the weak concordance between mtDNA and genomic data, most of the southern British populations consist largely of the Palearctic mtDNA haplogroup and the spread of mtDNA (but not nuclear DNA) further north could be retarded by cytoplasmic incompatibility with *Wolbachia* infected Italy-Iberia individuals.

On a time-line the possibility for this phased invasion of the British Isles since the Last Glacial Maximum is from some time after 18 kya to c. 12.9 kya and then again from *c*. 11.7 kya to *c*. 8 kya (after the Younger Dryas and before complete isolation from the continental mainland). During the glacial re-advance of the Younger Dryas (∼12.9-11.7 kya) *P. icarus* may have possibly persisted in a few refugial areas on south facing slopes in southern England. The Alicante-Provence and Iberia-Italy haplotypes likely entered the British Isles during, or before, this late cold-period. These potentially cold-adapted populations may have expanded northwards as the ice sheet retreated *c*. 11.7 while allowing for another recolonization (Palearctic haplotype) from continental Europe up until *c*. 8 kya when all land bridges were inundated. Contemporary populations on the Outer Hebrides are much better suited to flight at lower temperatures than southern populations on the British mainland (Howe *et al*., 2007). Additionally, the two-stage colonization hypothesis would imply that the colonization of Ireland likely occurred during the first colonization period before the separation of Ireland and mainland Britain c. 15 kya. Indeed mtDNA of individuals from Ireland harbour the same Alicante-Provence and Iberia-Italy mitotypes found in northern England and Scotland. This scenario has a direct and testable implication: genomes of Irish individuals should also cluster more closely with the northern British genomic cluster and could potentially be obtained in the future.

Although past climate events do support the idea of multiple invasion sequences for some butterfly species (Dennis, 1977), there is little direct genetic evidence for extant butterfly species of the British Isles to be the result of multiple distinct colonizations. Previous work on the mitochondrial genetic structuring of *Coenonympha tullia* has been suggested to indicate the possibility of two separate colonization events (Joyce *et al*., 2009), although this study did not include any continental samples and could also not make any strong inferences on routes of post-glacial colonization. The occurrence of distinct mitotypes of *Euphydryas aurinia* in the north and south of the British Isles have also been suggested to support a double colonization (Joyce & Pullin, 2001). However, in both cases (Joyce et al., 2009; Joyce & Pullin, 2001), low levels of nuclear variation, based on allozymes or a few nuclear markers, do not unequivocally support a hypothesis of two discrete colonization events. Notably, karyotype, mtDNA and nuclear DNA markers from several small mammals have also been suggested to support a two-stage colonization as proposed here (Searle *et al*., 2009). More recent phylogeographic analysis of the herbaceous perennial *Campanula rotundifolia* strongly suggests the possibility of two independent colonization events for this species from two distinct European refugium (Sutherland, Quarles, & Galloway, 2018; Wilson *et al*., 2020). Interestingly, *C. rotundifolila* exhibits two clusters, based on cytotype and ploidy level, within the British Isles that segregate between Ireland and western Britain mainland on one hand and from eastern and southern Britain mainland on the other.

Further corroboration for the possibility of a two-colonisation scenario for *P. icarus* would require detailed reconstruction of the recent demographic histories of the genomic clusters highlighted in this study. Dense genomic data, such as that from whole genome sequencing, may be able to provide demographic inference given the relatively recent ancestry and gene flow between these populations. Genomic data from Irish individuals would also help either support or refute the hypothesis. Finally, wider sampling of European populations would also be helpful in determining the number and potential sources of refugium.

### Potential Wolbachia-mediated feminization on the Outer Hebrides

Although, we currently have no evidence for the capability of either *w*Ica1 or *w*Ica2 to induce CI, two observations do indicate a potential phenotypic effect of the *w*Ica1 strain. Firstly, all females from the Outer Hebrides were predictedto be infected with *w*Ica but not all males, whereas infection status on the mainland exhibited no such dimorphism. Second, Outer Hebridean morphological females carried a *Wolbachia* load that was, based on our proxy metric for copy number, an order of magnitude higher than in morphological males. *Wolbachia* density can be indicative of its phenotypic effect on the host (Arai *et al*., 2020; Breeuwer & Werren, 1993). Our observations here are consistent with *Wolbachia*-induced feminization of genetic males (Stouthamer, Breeuwer, & Hurst, 1999). Such feminization would predict that female-specific markers in a species with a ZW sex determination system; with the caveat of partial homology between Z and W chromosomes would yield male (homozygous) genotypes in “feminized” males. As expected, all morphological females carrying the *w*Ica1 had male genotypes.

Feminization is a well-known reproductive manipulation strategy deployed by *Wolbachia* (Werren, Baldo, & Clark, 2008), however, it has not been encountered frequently within Lepidoptera (Duplouy & Hornett, 2018). The best documented instance of feminization in butterflies refers to the discovery of sex-biased female lines in two species of pierid *Eurema* butterflies in Japan (Kato, 2000). *w*Fem occurs at low frequencies in natural populations of *Eurema* and has not been detected in males. A causative role of *w*Fem in feminization has been suggested by antibiotic treatment of infected larva which results in intersex individuals (Narita, Kageyama, Nomura, & Fukatsu, 2007) and antibiotic treatment of adult females leads to all male progeny (Kern, Cook, Kageyama, & Riegler, 2015). The *w*Fem pattern of infection contrasts with that of *w*Ica1, where both females and males can be infected, but females carry a higher bacterial copy number. If *Wolbachia*-induced feminization indeed occurs in Outer Hebridean *P. icarus*, it will add to the small number of potential model systems to study the molecular basis underlying this poorly understood process in Lepidoptera. However, more direct evidence for *w*Ica1-induced feminization is required (Hiroki, Kato, Kamito, & Miura, 2002; Kageyama *et al*., 2017).

The potential existence of feminizing *Wolbachia* and the relative genetic isolation indicated the distinctiveness of the populations on the Outer Hebrides. Because of the current geographic isolation, we suggest that the populations on the Outer Hebrides represent a distinct genetic and ecological evolutionary unit that warrants consideration as being of conservation interest and monitoring. The eco-evolutionary dynamics of this *Wolbachia*-host system and their impact on past and future biology of these butterflies warrants further investigation.

## Conclusions

The contemporary population structure and phylogeography of the flora and fauna of the British Isles has directly resulted from the events following the last glacial period (Hewitt, 1999; Provan & Bennett, 2008). This paradigm of “southern richness and northern purity” implies limited intraspecific diversity and genetic structure in northern areas. Using mtDNA and ddRADseq data for *P. icarus* butterflies in the British Isles, we provide evidence for substantial and unexpected levels of genetic structuring and variability across a fine-scale spatial resolution. Geographic structuring of nuclear genomic variation and mtDNA variation was only weakly concordant and we argue that this pattern could be explained by multiple *Wolbachia*-meditated mtDNA sweeps and potentially two or three discrete colonization events of the British Isles before and after the glacial re-advance of the Younger Dryas *c*. 13 kya. We consider this a strong hypothesis that requires further corroboration and there may yet be alternative explanations for the genomic structuring observed within mainland Britain. It should be noted that several butterfly species in the British Isles exist as distinct geographical and/or genetic populations along a north-south divide and a case for multiple colonisations has been advanced on at least two occasions (Joyce et al., 2009; Joyce & Pullin, 2001) and there is also some evidence for two distinct colonization events in other flora and fauna (Searle *et al*., 2009; Wilson e*t al*., 2020). The current dearth of high resolution genomic data for the flora and fauna restricts any evaluation of the generality or plausibility of the sequential colonization hypothesis. However, with initiatives to provide high-quality genomes of all British flora and fauna (darwintreeoflife.org) and continued dropping costs of sequencing we may see a renewed interest and rigour in the phylogeographic reconstruction of the British Isles and beyond.

## Acknowledgements

This work was funded by Oxford Brookes University, Oxford, UK including a start-up fund to Saad Arif. LD is supported by the project funded by the Tuscan Archipelago National Park “Ricerca e conservazione sugli Impollinatori dell’Arcipelago Toscano e divulgazione sui Lepidotteri del parco”. We thank the late Bruce J. Riddoch for help collecting samples. We would like to thank the following for permissions to sample on their land: Mark Dinning and the Durham Wildlife Trust, Paul Nunns at the UK Forestry Commission, Alastair Gardner and Jenny Loring at Natural England, Simone Bullion and the Suffolk Wildlife Trust, and Simeon LD Jones and the Carmarthenshire County Council. This manuscript is dedicated to the memory of our dearly departed colleague: Bruce J Riddoch.

## Author Contributions

S.A., T.G.S and L.D. designed the study and conducted preliminary analysis. S.A., T.G.S, M.D.S.N and W.H.H-M performed all fieldwork. S.A. performed all laboratory work. S.A., L.D., W.G.H-M., M.G. and M.D.S.N. performed data analysis. S.A., T.G.S. and L.D. wrote the manuscript with input from all authors.

## Data Availability Statement

*CO1* sequence data are publicly available from NCBI or BOLD with individuals accession numbers available in Table S1. Additional *CO1* sequences generated for this study are deposited in NCBI Genbank with accession numbers also listed in Table S1. Raw ddRADSeq reads have been deposited on the NCBI SRA under the project accession PRJNAXXXX. Barcodes for de-multiplexing illumina data, processed data files and Bash scripts for the genotype calls and analyses can be found on github/xxxx.

