## Supplementary Information for "The dirty north: Evidence for multiple colonisations and Wolbachia infections shaping the genetic structure of the widespread butterfly *Polyommatus icarus* in the British Isles"

### Multiple colonisations and Wolbachia infections influence the fine-scale population structure of the widespread *Polyommatus icarus* in the British Isles

Saad Arif, Michael Gerth, William G. Hone-Millard, Maria D. S. Nunes, Leonardo Dapporto, Timothy G. Shreeve

Table S1: Sampling Localities in the British Isles

| Population abbreviation | Location | Sampling Date | Longitude | Latitude | Females | Males |
|---|---|---|---|---|---|---|
| MLG | Mallaig, Inverness-shire, Scotland | 11-July-2017 | -5.834874 | 56.99196 | 5 | 11 |
| MDC | Conwy, Conwy, Wales | 5-August-2017 | -3.843251 | 53.29568 | 8 | 4 |
| BER | Berneray, Northern Uist, Outer Hebrides, Scotland | 12-July-2017 | -7.213534 | 57.71385 | 4 | 12 |
| TUL | Traigh Uuige, Isle of Lewis, Outer Hebrides, Scotland | 15-July-2017 | -7.025334 | 58.18553 | 8 | 8 |
| DGC | Dornoch, Ross and Cromarty, Scotland | 17-July-2017 | -4.016982 | 57.87810 | 2 | 12 |
| RVS | Ravensheugh Sand Dunes, East Lothian, Scotland | 17-August-2018 | -2.596053 | 56.02397 | 2 | 4 |
| OBN | Oban, Argyll, Scotland | 19-July-2017 | -5.482266 | 56.43357 | 2 | 13 |
| BWD | Bellever Wood, Devon, England | 13-June-2018 | -3.898247 | 50.57975 | 2 | 5 |
| FRN | Aveyron, Occitanie, France | 28-July-2018 | 1.982669 | 44.15734 | 2 | 4 |
| PCP | Pembrey Country Park, Carmarthenshire, Wales | 2-June-2018 | -4.308992 | 51.67422 | 1 | 12 |
| ETB | Eastbourne, Sussex, England | 3-June-2018 | 0.243747 | 50.76919 | 1 | 13 |
| MMS | Martin's Meadows, Suffolk, England | 8-June-2018 | 1.255096 | 52.16802 | 6 | 8 |
| CFW | Chamber's Farm Wood, Lincolnshire, England | 16-June-2018 | -0.281520 | 53.25443 | 0 | 12 |
| RHD | Raisby Hill, Durham, England | 18-June-2018 | -1.477433 | 54.71391 | 3 | 10 |
| BMD | Bernwood Meadow's, Oxfordshire, England | 29-July-2017, 18-August-2018 | -1.125439 | 51.78256 | 8 | 8 |

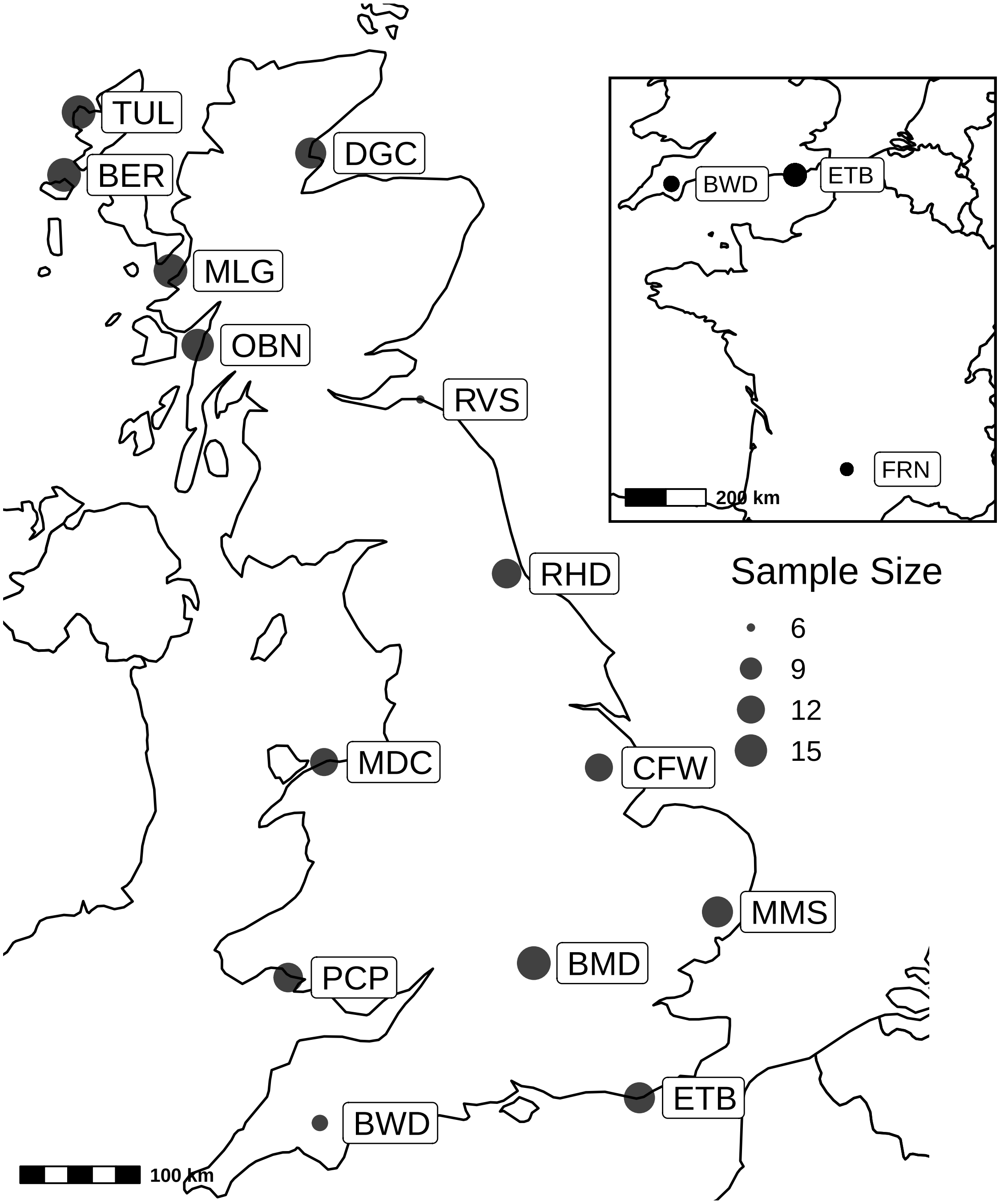


**Figure S1** A map of the British Isles with geographical locations of *Polyommatus icarus* sampled for this study. An additional 6 individuals were collected from southern France as an out-group (Inset). Size of circles is proportional to the number of samples acquired at each locality, except for the inset where circle are not to scale. For comparison RVS and FRN were both represented by 6 individuals here.

Table S2: Archived and newly generated *Polyommatus icarus* sequences used for CO1 mtDNA analysis

| Sample ID | Island | Geographic region | Latitude (dec. deg.) | Longitude (dec. deg.) | BOLD Process ID | GenBank Accession | Country |
|---|---|---|---|---|---|---|---|
| 150308PP10 | Crete | Eurasia | 35.15000 | 25.915600 | GBMIN32548-13 | JN084691 | Greece |
| 150308PP62 | Crete | Eurasia | 35.15000 | 25.915600 | GBMIN32571-13 | JN084692 | Greece |
| 150308PP67 | Crete | Eurasia | 35.15000 | 25.916000 | EULEP004-14 | KP871006 | Greece |
| 150308PP22 | Crete | Eurasia | 35.17665 | 25.709327 | GBMIN32539-13 | JN084709 | Greece |
| RVcoll11J518 | Karpathos | Karpathos | 35.56900 | 27.144000 | WMB6536-18 | NA | Greece |
| RVcoll11J519 | Karpathos | Karpathos | 35.56900 | 27.144000 | WMB6537-18 | NA | Greece |
| RVcoll11J508 | Karpathos | Karpathos | 35.59800 | 27.096000 | WMB6535-18 | NA | Greece |
| RVcoll11J530 | Rhodes | Rhodes | 36.23700 | 27.885000 | WMB6538-18 | NA | Greece |
| RVcoll11J497 | Nisyros | Bisyros | 36.61000 | 27.177000 | WMB6534-18 | NA | Greece |
| RVcoll14G023 | main | Eurasia | 36.98500 | 22.364000 | EULEP1397-15 | NA | Greece |
| RVcoll08J968 | main | Eurasia | 37.08100 | -3.378000 | EZSPN665-09 | GU676484 | Spain |
| RVcoll11I453 | main | Eurasia | 37.08100 | -3.378000 | GBGL20125-15 | KM459409 | Spain |
| RVcoll11I454 | main | Eurasia | 37.08100 | -3.378000 | EULEP162-14 | KM459410 | Spain |
| RVcoll11I456 | main | Eurasia | 37.08100 | -3.378000 | GBGL20126-15 | KM459411 | Spain |
| RVcoll11I457 | main | Eurasia | 37.08100 | -3.378000 | GBGL20127-15 | KM459412 | Spain |
| RVcoll11I458 | main | Eurasia | 37.08100 | -3.378000 | EULEP164-14 | KM459413 | Spain |
| RVcoll09V446 | main | Eurasia | 37.09400 | -3.115000 | EULEP104-14 | KM459361 | Spain |
| RVcoll08J960 | main | Eurasia | 37.10700 | -3.392000 | EZSPN661-09 | GU676489 | Spain |
| RVcoll08J962 | main | Eurasia | 37.10700 | -3.392000 | EULEP069-14 | KM459344 | Spain |
| RVcoll07D768 | main | Eurasia | 37.96000 | -2.550000 | EULEP027-14 | KP870765 | Spain |
| RVcoll07D769 | main | Eurasia | 37.96000 | -2.550000 | EULEP028-14 | KP870739 | Spain |
| RVcoll07D778 | main | Eurasia | 37.96000 | -2.550000 | EULEP029-14 | KP870755 | Spain |
| RVcoll09V486 | main | Eurasia | 37.96080 | -2.558100 | GBMIN32540-13 | JN084707 | Spain |
| RVcoll09V902 | main | Eurasia | 37.96080 | -2.558100 | GBMIN32544-13 | JN084699 | Spain |
| RVcoll09V488 | main | Eurasia | 37.96100 | -2.558000 | EULEP106-14 | KM459362 | Spain |
| RVcoll09V903 | main | Eurasia | 37.96100 | -2.558000 | WMB6530-18 | NA | Spain |
| RVcoll09V905 | main | Eurasia | 37.96100 | -2.558000 | EULEP109-14 | KM459363 | Spain |
| RVcoll09V907 | main | Eurasia | 37.96100 | -2.558000 | EZSPM467-09 | GU676068 | Spain |
| RVcoll09V941 | main | Eurasia | 37.96100 | -2.558000 | EZSPM465-09 | GU676071 | Spain |
| RVcoll09X366 | main | Eurasia | 37.96100 | -2.558000 | EULEP116-14 | KM459368 | Spain |
| RVcoll09X370 | main | Eurasia | 37.96100 | -2.558000 | GBGL20088-15 | KM459369 | Spain |
| RVcoll09X371 | main | Eurasia | 37.96100 | -2.558000 | GBGL20089-15 | KM459370 | Spain |
| RVcoll09X372 | main | Eurasia | 37.96100 | -2.558000 | EULEP118-14 | KM459371 | Spain |
| RVcoll09X373 | main | Eurasia | 37.96100 | -2.558000 | GBGL20090-15 | KM459372 | Spain |
| RVcoll09X374 | main | Eurasia | 37.96100 | -2.558000 | GBGL20091-15 | KM459373 | Spain |
| RVcoll06K697 | main | Eurasia | 38.00400 | -2.586000 | EZSPN171-09 | GU676767 | Spain |
| RVcoll06K699 | main | Eurasia | 38.00400 | -2.586000 | EZSPN173-09 | HM901841 | Spain |
| RVcoll14F830 | main | Eurasia | 38.03100 | 22.218000 | EULEP1340-15 | NA | Greece |
| RVcoll07C177 | main | Eurasia | 38.06800 | 36.146000 | GBMIN32569-13 | JN084696 | Turkey |
| RVcoll07C179 | main | Eurasia | 38.06800 | 36.146000 | GBGL20657-18 | NA | Turkey |
| RVcoll07F274 | main | Eurasia | 38.06800 | 36.146000 | WMB6528-18 | NA | Turkey |
| RVcollLD2371 | main | Eurasia | 38.12000 | 15.670000 | GBGL20150-15 | KM459436 | Italy |
| RVcollLD0292 | main | Eurasia | 38.19200 | 15.993000 | EULEP268-14 | KM459437 | Italy |
| RVcoll11I149 | main | Eurasia | 38.20100 | 15.980000 | GBGL20114-15 | KM459398 | Italy |
| RVcoll11I180 | main | Eurasia | 38.24000 | 15.710000 | GBGL20115-15 | KM459399 | Italy |
| RVcoll12M687 | main | Eurasia | 38.40200 | -3.954000 | EZSPM948-12 | KM517852 | Spain |

Table S2: Archived and newly generated *Polyommatus icarus* sequences used for CO1 mtDNA analysis *(continued)*

| Sample ID | Island | Geographic region | Latitude (dec. deg.) | Longitude (dec. deg.) | BOLD Process ID | GenBank Accession | Country |
|---|---|---|---:|---:|---|---|---|
| RVcoll08R537 | main | Eurasia | 38.46600 | 15.929000 | GBGL20084-15 | KM459356 | Italy |
| RVcoll07F225 | main | Eurasia | 38.51700 | 35.521000 | WMB6527-18 | NA | Turkey |
| RVcoll14O551 | main | Eurasia | 38.54300 | 22.585000 | EULEP3484-16 | NA | Greece |
| RVcoll14M901 | main | Eurasia | 38.54600 | -2.376000 | WMB5453-14 | NA | Spain |
| RVcoll07F178 | main | Eurasia | 38.55800 | 36.451000 | WMB6526-18 | NA | Turkey |
| RVcoll14D309 | main | Eurasia | 38.56500 | -2.265000 | WMB4447-14 | NA | Spain |
| RVcoll11E016 | main | Eurasia | 38.65800 | -0.310000 | EULEP160-14 | KM459387 | Spain |
| HBOK172-08 | main | Eurasia | 38.66700 | -2.491000 | HBOK172-08 | NA | Spain |
| HBOK173-08 | main | Eurasia | 38.66700 | -2.491000 | HBOK173-08 | NA | Spain |
| RVcoll08L281 | main | Eurasia | 38.77200 | -0.146000 | EZSPN772-09 | GU676372 | Spain |
| RVcoll07C260 | main | Eurasia | 39.00300 | 35.782000 | WMB6525-18 | NA | Turkey |
| RVcoll07E051 | main | Eurasia | 39.33600 | 16.358000 | EULEP034-14 | KM459332 | Italy |
| LEPSS00041 | main | Eurasia | 39.44330 | 16.603900 | BIBSA042-14 | NA | Italy |
| LEPSS00042 | main | Eurasia | 39.55970 | 16.750600 | BIBSA043-14 | NA | Italy |
| RVcoll08H394 | main | Eurasia | 39.58800 | -0.614000 | GBGL20068-15 | KM459336 | Spain |
| RVcoll08R025 | main | Eurasia | 39.65000 | -0.577000 | EZSPM345-09 | GU675891 | Spain |
| RVcoll08J743 | main | Eurasia | 39.82200 | -1.130000 | WMB3223-14 | NA | Spain |
| GWORR418-10 | main | Eurasia | 39.84250 | 15.992600 | GWORR418-10 | HM904304-SUPPRESSED | Italy |
| GWORU349-10 | main | Eurasia | 39.90170 | 16.115600 | GWORU349-10 | HM910575-SUPPRESSED | Italy |
| RVcoll11I239 | main | Eurasia | 39.93000 | 16.150000 | GBGL20116-15 | KM459400 | Italy |
| RVcoll12Q795 | main | Eurasia | 39.93000 | 16.170000 | WMB2847-13 | NA | Italy |
| RVcoll07D899 | main | Eurasia | 39.94200 | 16.147000 | EULEP033-14 | KM459331 | Italy |
| RVcoll14I327 | main | Eurasia | 39.99000 | 18.010000 | WMB4780-14 | NA | Italy |
| GWORZ057-10 | main | Eurasia | 39.99190 | 15.793100 | GWORZ057-10 | HM913968-SUPPRESSED | Italy |
| GWORR414-10 | main | Eurasia | 40.08310 | 15.727700 | GWORR414-10 | HM904300-SUPPRESSED | Italy |
| RVcoll07E062 | main | Eurasia | 40.14310 | 15.869200 | GBMIN32566-13 | JN084702 | Italy |
| RVcoll09X521 | main | Eurasia | 40.24800 | -1.582000 | EULEP120-14 | KM459374 | Spain |
| RVcoll08L092 | main | Eurasia | 40.25700 | -1.603000 | EULEP071-14 | KM459346 | Spain |
| RVcoll08L276 | main | Eurasia | 40.32100 | -0.332000 | EZSPC759-10 | HM901614 | Spain |
| RVcoll08H434 | main | Eurasia | 40.37300 | -3.366000 | EZSPN384-09 | GU676654 | Spain |
| RVcoll07F062 | main | Eurasia | 40.53900 | -0.146000 | EULEP035-14 | KM459333 | Spain |
| RVcoll08H410 | main | Eurasia | 40.54800 | -3.685000 | GBGL20069-15 | KM459337 | Spain |
| RVcoll09T560 | Capri | Capri | 40.55000 | 14.220000 | EULEP095-14 | KM459357 | Italy |
| RVcoll09T559 | Capri | Capri | 40.55000 | 14.230000 | GBMIN32564-13 | JN084706 | Italy |
| RVcoll10C516 | main | Eurasia | 40.57800 | 14.330000 | WMB3838-14 | NA | Italy |
| RVcoll09V396 | main | Eurasia | 40.65000 | 0.761000 | EULEP103-14 | KM459360 | Spain |
| RVcoll15C233 | main | Eurasia | 40.66700 | 16.614000 | BIBSA1044-15 | NA | Italy |
| RVcoll14A082 | main | Eurasia | 40.67000 | 14.473000 | WMB6539-18 | NA | Italy |
| RVcoll08H592 | main | Eurasia | 40.67100 | -2.673000 | EZSPN426-09 | GU676611 | Spain |
| RVcollLD2321 | Ischia | Ischia | 40.72900 | 13.884000 | GBGL20148-15 | KM459434 | Italy |
| RVcollLD2322 | Ischia | Ischia | 40.72900 | 13.884000 | GBGL20149-15 | KM459435 | Italy |
| RVcoll11J575 | Ischia | Ischia | 40.73000 | 13.900000 | GBGL20131-15 | KM459417 | Italy |
| RVcoll11J576 | Ischia | Ischia | 40.73000 | 13.900000 | GBGL20132-15 | KM459418 | Italy |
| RVcoll11J577 | Ischia | Ischia | 40.73000 | 13.900000 | GBGL20133-15 | KM459419 | Italy |
| RVcoll19C177 | main | Eurasia | 40.77700 | 17.414000 | BIBSA2030-19 | NA | Italy |
| RVcoll10B514 | main | Eurasia | 40.79200 | 0.310000 | GBGL20096-15 | KM459379 | Spain |

Table S2: Archived and newly generated *Polyommatus icarus* sequences used for CO1 mtDNA analysis *(continued)*

| Sample ID | Island | Geographic region | Latitude (dec. deg.) | Longitude (dec. deg.) | BOLD Process ID | GenBank Accession | Country |
|---|---|---|---|---|---|---|---|
| RVcoll10A841 | main | Eurasia | 40.82400 | 0.373000 | GBGL20094-15 | KM459377 | Spain |
| RVcoll10A866 | main | Eurasia | 40.82500 | 0.367000 | WMB3476-14 | NA | Spain |
| RVcoll15C144 | main | Eurasia | 40.92100 | 15.632000 | BIBSA1017-15 | NA | Italy |
| RVcoll12L143 | main | Eurasia | 40.98000 | -3.940000 | EZSPM752-12 | KM517851 | Spain |
| RVcoll12L149 | main | Eurasia | 41.07000 | -3.860000 | EULEP234-14 | KP870761 | Spain |
| RVcoll08M674 | main | Eurasia | 41.28000 | 0.866000 | EZSPC669-09 | GU669617 | Spain |
| RVcoll12Q454 | main | Eurasia | 41.28100 | -3.358000 | WMB3998-14 | NA | Spain |
| RVcoll12Q586 | main | Eurasia | 41.30000 | 13.617000 | GBGL20145-15 | KM459431 | Italy |
| RVcoll08H255 | main | Eurasia | 41.31300 | 0.130000 | EULEP048-14 | KM459335 | Spain |
| RVcoll12L152 | main | Eurasia | 41.33000 | -3.270000 | EULEP235-14 | KP870575 | Spain |
| RVcoll14F591 | main | Eurasia | 41.37100 | 23.633000 | EULEP1243-15 | NA | Greece |
| RVcoll11E685 | Corsica | Corsica | 41.37700 | 9.179000 | GBGL20104-15 | KM459388 | France |
| RVcoll11E686 | Corsica | Corsica | 41.37700 | 9.179000 | GBGL20105-15 | KM459389 | France |
| RVcoll09T515 | Corsica | Corsica | 41.38700 | 9.165000 | GBMIN32543-13 | JN084701 | France |
| RVcoll14D350 | main | Eurasia | 41.44400 | -4.513000 | WMB4468-14 | NA | Spain |
| RVcoll15M827 | main | Eurasia | 41.45700 | 14.382000 | BIBSA1335-15 | NA | Italy |
| RVcoll07C463 | main | Eurasia | 41.50600 | 2.098000 | EZSPC284-09 | JN114415 | Spain |
| RVcoll08H232 | main | Eurasia | 41.50700 | 2.099000 | EZSPC662-09 | GU669625 | Spain |
| RVcoll08J116 | main | Eurasia | 41.51200 | -8.077000 | WMB6529-18 | NA | Portugal |
| RVcoll12O004 | Corsica | Corsica | 41.53400 | 8.867000 | GBGL20140-15 | KM459426 | France |
| RVcoll10A994 | main | Eurasia | 41.55400 | 23.614000 | WMB6532-18 | NA | Bulgaria |
| RVcoll07D884 | main | Eurasia | 41.59830 | 13.099400 | GBMIN32541-13 | JN084705 | Italy |
| RVcoll06G507 | main | Eurasia | 41.63700 | -0.346000 | EZSPN142-09 | GU676797 | Spain |
| RVcoll08J127 | main | Eurasia | 41.68200 | -7.698000 | EZSPN578-09 | GU676569 | Portugal |
| RVcoll08J150 | main | Eurasia | 41.70000 | -7.650000 | EULEP051-14 | KM459341 | Portugal |
| RVcoll13S657 | main | Eurasia | 41.72000 | 15.760000 | WMB4142-14 | NA | Italy |
| RVcoll11E763 | Corsica | Corsica | 41.76500 | 9.147000 | GBGL20106-15 | KM459390 | France |
| RVcoll09V960 | main | Eurasia | 41.76610 | 23.421600 | GBMIN32545-13 | JN084697 | Bulgaria |
| RVcoll07D812 | main | Eurasia | 41.76690 | 12.315300 | GBMIN32567-13 | JN084700 | Italy |
| RVcoll15A610 | main | Eurasia | 41.79500 | 12.220000 | OXB1077-15 | NA | Italy |
| RVcoll06G433 | main | Eurasia | 41.80900 | 2.296000 | EZROM751-08 | JN114417 | Spain |
| RVcoll07D864 | main | Eurasia | 41.82300 | 13.276000 | EULEP032-14 | KM459330 | Italy |
| RVcoll12L141 | main | Eurasia | 41.84000 | -3.340000 | EZSPM751-12 | KM517859 | Spain |
| RVcoll08L473 | main | Eurasia | 41.86600 | 0.598000 | EZSPC666-09 | GU669622 | Spain |
| RVcoll13S266 | main | Eurasia | 41.88900 | 1.053000 | WMB4044-14 | NA | Spain |
| RVcoll08J199 | main | Eurasia | 41.89000 | -7.730000 | EULEP052-14 | KM459342 | Portugal |
| RVcoll15M991 | main | Eurasia | 41.95500 | 14.966000 | BIBSA1054-15 | NA | Italy |
| RVcoll12L159 | main | Eurasia | 41.98000 | -2.070000 | EZSPM754-12 | KM517862 | Spain |
| RVcoll11E830 | Corsica | Corsica | 41.98800 | 9.190000 | GBGL20107-15 | KM459391 | France |
| RVcoll12O000 | Corsica | Corsica | 42.01700 | 8.733000 | GBGL20138-15 | KM459424 | France |
| RVcoll12O002 | Corsica | Corsica | 42.03300 | 9.033000 | GBGL20139-15 | KM459425 | France |
| RVcoll12L160 | main | Eurasia | 42.07000 | -2.630000 | EULEP238-14 | KP870886 | Spain |
| RVcoll11E853 | Corsica | Corsica | 42.07600 | 9.194000 | GBGL20108-15 | KM459392 | France |
| RVcoll11E854 | Corsica | Corsica | 42.07600 | 9.194000 | GBGL20109-15 | KM459393 | France |
| RVcoll11E865 | Corsica | Corsica | 42.09300 | 9.323000 | GBGL20110-15 | KM459394 | France |
| RVcoll13S568 | San Dominp | San Domino | 42.11100 | 15.486000 | WMB3040-14 | NA | Italy |

Table S2: Archived and newly generated *Polyommatus icarus* sequences used for CO1 mtDNA analysis *(continued)*

| **Sample ID** | **Island** | **Geographic region** | **Latitude (dec. deg.)** | **Longitude (dec. deg.)** | **BOLD Process ID** | **GenBank Accession** | **Country** |
|---|---|---|---:|---:|---|---|---|
| RVcoll15M985 | main | Eurasia | 42.13300 | 14.657000 | BIBSA1389-15 | NA | Italy |
| RVcoll08R488 | main | Eurasia | 42.13600 | -8.576000 | EZSPM271-09 | GU675792 | Spain |
| RVcoll11E873 | Corsica | Corsica | 42.16300 | 9.267000 | GBGL20111-15 | KM459395 | France |
| RVcoll12L153 | main | Eurasia | 42.17100 | -2.288000 | EZSPM753-12 | KM517856 | Spain |
| RVcoll12L154 | main | Eurasia | 42.18000 | -2.290000 | EULEP236-14 | KP870585 | Spain |
| RVcoll12L155 | main | Eurasia | 42.19000 | -2.510000 | EULEP237-14 | KP871101 | Spain |
| RVcoll11J554 | main | Eurasia | 42.19500 | 3.095000 | WMB3698-14 | NA | Spain |
| RVcoll16L194 | main | Eurasia | 42.21000 | 11.715000 | EULEP5722-17 | NA | Italy |
| RVcoll13S518 | main | Eurasia | 42.21300 | 14.061000 | WMB5172-14 | NA | Italy |
| RVcollLD1552 | Corsica | Corsica | 42.28600 | 8.887000 | GBGL20147-15 | KM459433 | France |
| RVcollLD0479 | Corsica | Corsica | 42.28800 | 9.198000 | GBGL20151-15 | KM459438 | France |
| RVcoll11E884 | Corsica | Corsica | 42.29700 | 9.182000 | GBGL20112-15 | KM459396 | France |
| RVcoll11E885 | Corsica | Corsica | 42.29700 | 9.182000 | GBGL20113-15 | KM459397 | France |
| RVcollLD1550 | Corsica | Corsica | 42.33000 | 9.480000 | GBGL20146-15 | KM459432 | France |
| RVcoll08P076 | main | Eurasia | 42.35000 | 1.717000 | EZSPC283-09 | JN114416 | Spain |
| RVcoll11J235 | main | Eurasia | 42.36300 | 2.998000 | WMB3688-14 | NA | Spain |
| RVcoll11J238 | main | Eurasia | 42.36300 | 2.998000 | WMB3689-14 | NA | Spain |
| RVcoll09V894 | main | Eurasia | 42.40000 | -2.904000 | WMB3407-14 | NA | Spain |
| RVcoll11I382 | Argentario | Argentario | 42.42800 | 11.159000 | GBGL20122-15 | KM459406 | Italy |
| RVcoll11I383 | Argentario | Argentario | 42.42800 | 11.159000 | GBGL20123-15 | KM459407 | Italy |
| RVcoll11I384 | Argentario | Argentario | 42.42800 | 11.159000 | GBGL20124-15 | KM459408 | Italy |
| RVcoll11J578 | Argentario | Argentario | 42.42800 | 11.159000 | GBGL20134-15 | KM459420 | Italy |
| RVcoll11J579 | Argentario | Argentario | 42.42800 | 11.159000 | GBGL20135-15 | KM459421 | Italy |
| RVcoll12Q578 | main | Eurasia | 42.43300 | 13.583000 | GBGL20144-15 | KM459430 | Italy |
| RVcoll07C623 | main | Eurasia | 42.44800 | 1.781000 | EZSPC285-09 | JN114414 | Spain |
| RVcoll14A121 | main | Eurasia | 42.45800 | 11.421000 | BIBSA969-15 | NA | Italy |
| RVcoll08P209 | main | Eurasia | 42.48900 | 1.856000 | GBGL20079-15 | KM459351 | France |
| RVcollLD0384 | Corsica | Corsica | 42.52000 | 9.167000 | WMB6070-18 | NA | France |
| RVcollLD0385 | Corsica | Corsica | 42.54000 | 9.180000 | GBMIN16302-13 | JX678166 | France |
| OXBTGS1270 | main | Eurasia | 42.57780 | 1.666050 | OXB1552-16 | NA | Andorra |
| OXBTGS1271 | main | Eurasia | 42.57780 | 1.666050 | OXB1553-16 | NA | Andorra |
| RVcoll11I332 | Pianosa | Pianosa | 42.58000 | 10.080000 | GBGL20117-15 | KM459401 | Italy |
| RVcoll11I333 | Pianosa | Pianosa | 42.58000 | 10.080000 | GBGL20118-15 | KM459402 | Italy |
| RVcoll11I334 | Pianosa | Pianosa | 42.58000 | 10.080000 | GBGL20119-15 | KM459403 | Italy |
| RVcoll11I335 | Pianosa | Pianosa | 42.58000 | 10.080000 | GBGL20120-15 | KM459404 | Italy |
| RVcoll11I336 | Pianosa | Pianosa | 42.58000 | 10.080000 | GBGL20121-15 | KM459405 | Italy |
| RVcoll14D922 | main | Eurasia | 42.58000 | 11.130000 | WMB4480-14 | NA | Italy |
| RVcoll08J899 | main | Eurasia | 42.72800 | -6.648000 | EZSPM325-09 | GU675911 | Spain |
| RVcoll09T572 | Elba | Elba | 42.75200 | 10.205000 | EULEP097-14 | KM459358 | Italy |
| RVcoll10C439 | Elba | Elba | 42.75200 | 10.205000 | GBGL20098-15 | KM459381 | Italy |
| RVcoll10C440 | Elba | Elba | 42.75200 | 10.205000 | GBGL20099-15 | KM459382 | Italy |
| RVcoll08R458 | main | Eurasia | 42.76700 | -8.694000 | EZSPM246-09 | GU675791 | Spain |
| RVcoll11J568 | Elba | Elba | 42.78500 | 10.391000 | GBGL20129-15 | KM459415 | Italy |
| RVcoll11J569 | Elba | Elba | 42.78500 | 10.391000 | GBGL20130-15 | KM459416 | Italy |
| RVcoll09V741 | main | Eurasia | 42.82100 | -0.331000 | WMB3399-14 | NA | Spain |
| RVcoll14D348 | main | Eurasia | 42.83500 | -4.647000 | WMB4467-14 | NA | Spain |

Table S2: Archived and newly generated *Polyommatus icarus* sequences used for CO1 mtDNA analysis *(continued)*

| Sample ID | Island | Geographic region | Latitude (dec. deg.) | Longitude (dec. deg.) | BOLD Process ID | GenBank Accession | Country |
|---|---|---|---|---|---|---|---|
| RVcoll14W014 | main | Eurasia | 42.86000 | 10.970000 | BIBSA978-15 | NA | Italy |
| RVcoll12P242 | main | Eurasia | 42.92500 | 2.958000 | WMB5151-14 | NA | France |
| RVcoll130711PX33 | main | Eurasia | 42.94100 | -6.588000 | EZSPM874-12 | KM517857 | Spain |
| RVcoll14E286 | Corsica | Corsica | 42.95500 | 9.446000 | BIBSA1294-15 | NA | France |
| RVcoll09X800 | main | Eurasia | 42.96200 | 10.532000 | GBGL20092-15 | KM459375 | Italy |
| RVcoll12O603 | Levant | Levant | 43.02000 | 6.434000 | GBGL20142-15 | KM459428 | France |
| RVcoll14A389 | main | Eurasia | 43.02300 | 13.627000 | WMB4293-14 | NA | Italy |
| RVcoll14A397 | main | Eurasia | 43.02300 | 13.627000 | WMB4300-14 | NA | Italy |
| RVcoll08P416 | main | Eurasia | 43.02900 | -5.066000 | GBMIN32565-13 | JN084704 | Spain |
| RVcoll10C177 | main | Eurasia | 43.03900 | 2.926000 | WMB119-11 | KM517843 | France |
| RVcoll10C178 | main | Eurasia | 43.03900 | 2.926000 | WMB120-11 | KM517840 | France |
| RVcoll10C180 | main | Eurasia | 43.03900 | 2.926000 | GBGL20097-15 | KM459380 | France |
| RVcoll130711PX19 | main | Eurasia | 43.04400 | -6.603000 | EZSPM873-12 | KM517858 | Spain |
| RVcoll13S619 | Capraia | Capraia | 43.05000 | 9.820000 | GBGL20059-15 | KM459322 | Italy |
| RVcoll13S620 | Capraia | Capraia | 43.05000 | 9.820000 | GBGL20058-15 | KM459321 | Italy |
| RVcoll13S621 | Capraia | Capraia | 43.05000 | 9.820000 | GBGL20152-15 | KM459439 | Italy |
| RVcoll11Y003 | main | Eurasia | 43.06000 | 10.610000 | GBGL20137-15 | KM459423 | Italy |
| RVcoll09X551 | main | Eurasia | 43.08500 | -5.361000 | WMB3438-14 | NA | Spain |
| RVcoll08P653 | main | Eurasia | 43.09500 | -5.858000 | EZSPM141-09 | GU675755 | Spain |
| RVcoll12P295 | main | Eurasia | 43.13300 | 3.061000 | WMB3904-14 | NA | France |
| RVcoll10C501 | main | Eurasia | 43.13600 | 11.560000 | GBGL20101-15 | KM459384 | Italy |
| RVcoll15A519 | main | Eurasia | 43.50800 | 12.318000 | OXB877-15 | NA | Italy |
| RVcoll12O601 | main | Eurasia | 43.51600 | 3.668000 | GBGL20141-15 | KM459427 | France |
| RVcoll09V243 | main | Eurasia | 43.55400 | 5.730000 | EULEP102-14 | KM459359 | France |
| RVcoll10B623 | main | Eurasia | 43.55400 | 5.730000 | WMB106-11 | KM517834 | France |
| RVcoll10A444 | main | Eurasia | 43.59600 | 5.167000 | GBGL20093-15 | KM459376 | France |
| RVcoll12P711 | main | Eurasia | 43.72100 | 4.821000 | WMB3950-14 | NA | France |
| RVcoll15A904 | main | Eurasia | 43.76800 | 12.981000 | OXB954-15 | NA | Italy |
| RVcoll15A548 | main | Eurasia | 43.83100 | 11.836000 | OXB904-15 | NA | Italy |
| RVcoll16A047 | main | Eurasia | 43.86600 | 10.333000 | OXB1160-15 | NA | Italy |
| RVcoll10C505 | main | Eurasia | 43.89000 | 11.130000 | GBGL20102-15 | KM459385 | Italy |
| RVcoll09X205 | main | Eurasia | 43.89600 | 5.920000 | GBGL20087-15 | KM459366 | France |
| RVcoll09X210 | main | Eurasia | 43.89600 | 5.920000 | EULEP113-14 | KM459367 | France |
| RVcoll12Q394 | main | Eurasia | 43.97200 | 7.389000 | WMB3993-14 | NA | France |
| RVcoll10C500 | main | Eurasia | 44.00000 | 7.860000 | GBGL20100-15 | KM459383 | Italy |
| RVcoll10C507 | main | Eurasia | 44.00200 | 7.946000 | GBGL20103-15 | KM459386 | Italy |
| RVcoll12P526 | main | Eurasia | 44.01300 | 3.849000 | WMB5153-14 | NA | France |
| RVcoll09X222 | main | Eurasia | 44.07110 | 5.356100 | GBMIN32573-13 | JN084688 | France |
| RVcoll14N977 | main | Eurasia | 44.10500 | 12.160000 | OXB588-15 | NA | Italy |
| RVcoll12O605 | main | Eurasia | 44.10900 | 7.308000 | GBGL20143-15 | KM459429 | France |
| RVcoll10A606 | main | Eurasia | 44.12100 | 6.225000 | WMB3461-14 | NA | France |
| RVcoll09X260 | main | Eurasia | 44.14220 | 5.136100 | GBMIN32568-13 | JN084698 | France |
| RVcoll15A532 | main | Eurasia | 44.17200 | 9.776000 | OXB890-15 | NA | Italy |
| RVcoll14E067 | main | Eurasia | 44.20100 | 7.240000 | BIBSA268-15 | NA | Italy |
| RVcoll14I531 | main | Eurasia | 44.22600 | 9.550000 | WMB4984-14 | NA | Italy |
| RVcoll14I533 | main | Eurasia | 44.22600 | 9.550000 | WMB4986-14 | NA | Italy |

Table S2: Archived and newly generated *Polyommatus icarus* sequences used for CO1 mtDNA analysis *(continued)*

| **Sample ID** | **Island** | **Geographic region** | **Latitude (dec. deg.)** | **Longitude (dec. deg.)** | **BOLD Process ID** | **GenBank Accession** | **Country** |
|---|---|---|---:|---:|---|---|---|
| RVcoll11Y001 | main | Eurasia | 44.39900 | 8.512000 | GBGL20136-15 | KM459422 | Italy |
| RVcoll14E116 | main | Eurasia | 44.41600 | 6.995000 | BIBSA314-15 | NA | Italy |
| OXBTGS1308 | main | Eurasia | 44.44470 | 5.210590 | OXB1590-16 | NA | France |
| RVcoll14I590 | main | Eurasia | 44.44800 | 1.417000 | WMB5293-14 | NA | France |
| RVcoll08M425 | main | Eurasia | 44.46600 | 28.480000 | EZRMN209-08 | HQ005054 | Romania |
| RVcoll16A007 | main | Eurasia | 44.52000 | -0.600000 | BIBSA916-15 | NA | France |
| RVcoll14D555 | main | Eurasia | 44.52000 | 8.700000 | BIBSA120-15 | NA | Italy |
| RVcoll16A000 | main | Eurasia | 44.59000 | -0.580000 | BIBSA909-15 | NA | France |
| RVcoll11J219 | main | Eurasia | 44.64200 | 6.076000 | GBGL20128-15 | KM459414 | France |
| RVcoll10B458 | main | Eurasia | 44.68800 | 15.381000 | GBGL20095-15 | KM459378 | Croatia |
| OXBTGS1307 | main | Eurasia | 44.79950 | 5.259680 | OXB1589-16 | NA | France |
| RVcoll14L193 | main | Eurasia | 44.80900 | 11.100000 | WMB5062-14 | NA | Italy |
| RVcoll14L220 | main | Eurasia | 44.83600 | 12.250000 | WMB5089-14 | NA | Italy |
| RVcoll07D053 | main | Eurasia | 44.85200 | 28.872000 | EZRMN210-08 | HQ005055 | Romania |
| LEATJ1211-16 | main | Eurasia | 44.85700 | 13.947000 | LEATJ1211-16 | NA | Croatia |
| RVcoll07C964 | main | Eurasia | 44.87100 | 22.414000 | EZROM504-08 | HQ005047 | Romania |
| RVcoll14L206 | main | Eurasia | 44.92100 | 11.574000 | WMB5075-14 | NA | Italy |
| RVcoll15A942 | main | Eurasia | 44.93100 | 10.366000 | OXB992-15 | NA | Italy |
| RVcoll08M230 | main | Eurasia | 44.97100 | 25.687000 | EZRMN206-08 | HQ005056 | Romania |
| RVcoll15A952 | main | Eurasia | 44.97400 | 10.414000 | OXB1002-15 | NA | Italy |
| RVcoll14I008 | main | Eurasia | 45.03400 | 8.900000 | BIBSA340-15 | NA | Italy |
| RVcoll06M943 | main | Eurasia | 45.09400 | 26.533000 | EZROM500-08 | HQ005051 | Romania |
| RVcoll14G762 | main | Eurasia | 45.09600 | 28.389000 | EULEP1957-15 | NA | Romania |
| RVcoll15M327 | main | Eurasia | 45.12700 | 10.828000 | BIBSA1209-15 | NA | Italy |
| RVcoll07D104 | main | Eurasia | 45.21700 | 26.555000 | EZROM502-08 | HQ005048 | Romania |
| RVcoll08M630 | main | Eurasia | 45.29900 | 22.894000 | EZRMN208-08 | HQ005052 | Romania |
| RVcoll14N078 | main | Eurasia | 45.31000 | 11.696000 | OXB364-15 | NA | Italy |
| RVcoll15A655 | main | Eurasia | 45.32800 | 9.509000 | OXB1122-15 | NA | Italy |
| RVcoll15F830 | main | Eurasia | 45.65800 | -1.120000 | BIBSA1250-15 | NA | France |
| RVcoll07E207 | main | Eurasia | 45.66670 | 7.229700 | GBMIN32542-13 | JN084703 | Italy |
| RVcoll14I094 | main | Eurasia | 45.71000 | 7.472000 | BIBSA426-15 | NA | Italy |
| RVcoll11I974 | main | Eurasia | 45.86900 | 6.681000 | WMB3671-14 | NA | France |
| RVcoll14N971 | main | Eurasia | 45.88100 | 10.885000 | OXB582-15 | NA | Italy |
| LEPAA120-16 | main | Eurasia | 45.90680 | 8.920580 | NA | MK186704 | Switzerland |
| RVcoll15L950 | main | Eurasia | 45.94600 | 13.591000 | BIBSA1137-15 | NA | Italy |
| RVcoll15M107 | main | Eurasia | 46.09200 | 6.403000 | OXB1253-15 | NA | France |
| LEPAA348-16 | main | Eurasia | 46.09600 | 7.115380 | NA | MK186701 | Switzerland |
| RVcoll15M134 | main | Eurasia | 46.11600 | 5.628000 | OXB1274-15 | NA | France |
| RVcoll15L971 | main | Eurasia | 46.13000 | 13.499000 | BIBSA1139-15 | NA | Italy |
| RVcoll14N053 | main | Eurasia | 46.15300 | 8.333000 | OXB339-15 | NA | Italy |
| RVcoll08M374 | main | Eurasia | 46.18100 | 27.251000 | EZRMN207-08 | HQ005053 | Romania |
| RVcoll15M205 | main | Eurasia | 46.27800 | 11.419000 | BIBSA1175-15 | NA | Italy |
| RVcoll15G529 | main | Eurasia | 46.29767 | 8.063680 | EULEP4529-16 | NA | Switzerland |
| GWOSZ099-11 | main | Eurasia | 46.30130 | 11.445700 | GWOSZ099-11 | KX040372 | Italy |
| RVcoll15F875 | main | Eurasia | 46.43300 | -1.027000 | BIBSA1259-15 | NA | France |
| RVcoll15G985 | main | Eurasia | 46.45880 | 8.681800 | EULEP4530-16 | NA | Switzerland |

Table S2: Archived and newly generated *Polyommatus icarus* sequences used for CO1 mtDNA analysis *(continued)*

| **Sample ID** | **Island** | **Geographic region** | **Latitude (dec. deg.)** | **Longitude (dec. deg.)** | **BOLD Process ID** | **GenBank Accession** | **Country** |
|---|---|---|---:|---:|---|---|---|
| RVcoll15H602 | main | Eurasia | 46.49690 | 9.908600 | EULEP4531-16 | NA | Switzerland |
| LEASS697-17 | main | Eurasia | 46.56810 | 14.350800 | LEASS697-17 | NA | Austria |
| RVcoll13U411 | main | Eurasia | 46.58900 | 12.853000 | OXB746-15 | NA | Italy |
| PHLAC364-10 | main | Eurasia | 46.59700 | 11.439000 | PHLAC364-10 | JN820118 | Italy |
| LEATD021-13 | main | Eurasia | 46.60600 | 10.553000 | LEATD021-13 | NA | Italy |
| LEPAA517-16 | main | Eurasia | 46.62050 | 9.331900 | NA | MK186703 | Switzerland |
| RVcoll06V653 | main | Eurasia | 46.69800 | 23.549000 | EZROM501-08 | HQ005049 | Romania |
| LEASS917-17 | main | Eurasia | 46.78270 | 13.039200 | LEASS917-17 | NA | Austria |
| LEASS536-17 | main | Eurasia | 46.78330 | 15.533300 | LEASS536-17 | NA | Austria |
| RVcoll07D439 | main | Eurasia | 46.79900 | 23.959000 | EZROM503-08 | HQ005050 | Romania |
| RVcoll15I355 | main | Eurasia | 46.84066 | 13.438470 | EULEP4532-16 | NA | Austria |
| ABOLD435-16 | main | Eurasia | 47.06700 | 15.650000 | ABOLD435-16 | NA | Austria |
| LEPAA061-16 | main | Eurasia | 47.08550 | 7.109410 | NA | MK186705 | Switzerland |
| LEPAA130-16 | main | Eurasia | 47.08550 | 7.109410 | NA | MK186702 | Switzerland |
| PHLAH454-12 | main | Eurasia | 47.15200 | 10.166000 | PHLAH454-12 | KM573399 | Austria |
| LEPAA372-16 | main | Eurasia | 47.16910 | 8.691700 | NA | MK186706 | Switzerland |
| RVcoll15I899 | main | Eurasia | 47.25298 | 9.545890 | EULEP4534-16 | NA | Liechtenstein |
| RVcoll15M757 | main | Eurasia | 47.26100 | 4.570000 | OXB1392-15 | NA | France |
| RVcoll19C151 | main | Eurasia | 47.30300 | 1.349000 | BIBSA1989-19 | NA | France |
| LEATG234-14 | main | Eurasia | 47.30800 | 11.200000 | LEATG234-14 | NA | Austria |
| TLMFLep13865 | main | Eurasia | 47.31100 | 11.721000 | LEATG078-14 | NA | Austria |
| RVcoll16L110 | BelleIleenMer | BelleIleenMer | 47.31300 | -3.205000 | BIBSA1958-19 | NA | France |
| RVcoll16L111 | BelleIleenMer | BelleIleenMer | 47.31300 | -3.205000 | BIBSA1959-19 | NA | France |
| RVcoll15M788 | main | Eurasia | 47.32000 | 4.040000 | OXB1413-15 | NA | France |
| RVcoll16L138 | BelleIleenMer | BelleIleenMer | 47.36800 | -3.203000 | BIBSA1960-19 | NA | France |
| RVcoll16L139 | BelleIleenMer | BelleIleenMer | 47.36800 | -3.203000 | BIBSA1961-19 | NA | France |
| RVcoll15I714 | main | Eurasia | 47.45970 | 13.618100 | EULEP4533-16 | NA | Austria |
| RVcoll16L174 | main | Eurasia | 47.54100 | -3.135000 | BIBSA1970-19 | NA | France |
| GBLAA056-14 | main | Eurasia | 47.55610 | 7.679440 | GBLAA056-14 | MH419055 | Germany |
| RVcoll16L146 | main | Eurasia | 47.56800 | -3.133000 | BIBSA1969-19 | NA | France |
| RVcoll07C366 | main | Eurasia | 47.63300 | 25.367000 | EZROM593-08 | HQ005046 | Romania |
| RVcoll15M718 | main | Eurasia | 47.65300 | 3.758000 | OXB1363-15 | NA | France |
| RVcoll15M727 | main | Eurasia | 47.65300 | 3.758000 | OXB1367-15 | NA | France |
| RVcoll14U783 | main | Eurasia | 47.75000 | 1.980000 | OXB820-15 | NA | France |
| GWORO805-09 | main | Eurasia | 47.81600 | 11.480000 | GWORO805-09 | GU688449 | Germany |
| RVcoll15M642 | main | Eurasia | 47.88500 | 3.228000 | OXB1216-15 | NA | France |
| LEASS533-17 | main | Eurasia | 47.94000 | 16.711700 | LEASS533-17 | NA | Austria |
| ABOLD070-16 | main | Eurasia | 48.03300 | 16.250000 | ABOLD070-16 | NA | Austria |
| FBLMT902-09 | main | Eurasia | 48.16460 | 11.481500 | FBLMT902-09 | GU655005 | Germany |
| RVcoll14U785 | main | Eurasia | 48.28000 | 0.020000 | OXB822-15 | NA | France |
| ODOPE242-11 | main | Eurasia | 48.32220 | 10.929100 | ODOPE242-11 | KX044865 | Germany |
| RVcoll16I980 | main | Eurasia | 48.56280 | 20.403300 | EULEP5032-16 | NA | Slovakia |
| RVcoll15M678 | main | Eurasia | 48.68800 | 1.919000 | OXB1334-15 | NA | France |
| ODOPE364-11 | main | Eurasia | 48.91750 | 11.916400 | ODOPE364-11 | KX040462 | Germany |
| BCZSMLep25454 | main | Eurasia | 49.04450 | 12.497300 | NA | NA | Germany |
| FBLMT894-09 | main | Eurasia | 49.04450 | 12.497300 | FBLMT894-09 | HM391783 | Germany |

Table S2: Archived and newly generated *Polyommatus icarus* sequences used for CO1 mtDNA analysis *(continued)*

| **Sample ID** | **Island** | **Geographic region** | **Latitude (dec. deg.)** | **Longitude (dec. deg.)** | **BOLD Process ID** | **GenBank Accession** | **Country** |
|---|---|---|---:|---:|---|---|---|
| BCZSMLep75749 | main | Eurasia | 49.18910 | 7.204800 | NA | NA | Germany |
| GWOSU026-11 | main | Eurasia | 49.90320 | 9.823230 | GWOSU026-11 | KX046037 | Germany |
| RVcoll14V039 | main | Eurasia | 50.18300 | 36.400000 | EULEP2351-15 | NA | Ukraine |
| OXBTGS945 | main | Eurasia | 50.21000 | -3.709000 | OXB570-15 | NA | United Kingdom |
| RVcoll14B625 | main | Eurasia | 50.21000 | 36.430000 | WMB6540-18 | NA | Ukraine |
| RVcoll15M616 | main | Eurasia | 50.28400 | 2.958000 | OXB1200-15 | NA | France |
| BCZSMLep37416 | main | Eurasia | 50.36620 | 11.857200 | NA | NA | Germany |
| FBLMW315-10 | main | Eurasia | 50.36620 | 11.857200 | FBLMW315-10 | HQ563555 | Germany |
| OXBTGS343 | main | Eurasia | 50.37400 | -5.131000 | OXB458-15 | NA | United Kingdom |
| OXBTGS344 | main | Eurasia | 50.37400 | -5.131000 | OXB459-15 | NA | United Kingdom |
| RVcoll07C552 | main | Eurasia | 50.44000 | 8.920000 | GBMIN32547-13 | JN084693 | Germany |
| OXBTGS944 | main | Eurasia | 50.51000 | -4.110000 | OXB569-15 | NA | United Kingdom |
| OXBTGS943 | main | Eurasia | 50.57000 | -3.900000 | OXB568-15 | NA | United Kingdom |
| OXBTGS731 | main | Eurasia | 50.59800 | -1.971000 | OXB510-15 | NA | United Kingdom |
| OXBTGS732 | main | Eurasia | 50.59800 | -1.971000 | OXB511-15 | NA | United Kingdom |
| OXBTGS674 | main | Eurasia | 50.66700 | -2.097000 | OXB503-15 | NA | United Kingdom |
| OXBTGS675 | main | Eurasia | 50.66700 | -2.097000 | OXB504-15 | NA | United Kingdom |
| RVcoll15M583 | main | Eurasia | 50.70200 | 2.234000 | OXB1317-15 | NA | France |
| OXBTGS951 | main | Eurasia | 50.81000 | -1.101000 | OXB268-15 | NA | United Kingdom |
| RVcoll14V231 | main | Eurasia | 50.96400 | 2.953000 | EULEP2433-15 | NA | Belgium |
| OXBTGS650 | main | Eurasia | 51.26100 | -2.143000 | OXB496-15 | NA | United Kingdom |
| OXBTGS651 | main | Eurasia | 51.26100 | -2.143000 | OXB497-15 | NA | United Kingdom |
| OXBTGS456 | main | Eurasia | 51.28100 | -0.710000 | OXB477-15 | NA | United Kingdom |
| OXBTGS457 | main | Eurasia | 51.28100 | -0.710000 | OXB478-15 | NA | United Kingdom |
| OXBTGS468 | main | Eurasia | 51.28100 | -0.710000 | OXB480-15 | NA | United Kingdom |
| OXBTGS793 | main | Eurasia | 51.48200 | -3.621000 | OXB521-15 | NA | United Kingdom |
| OXBTGS794 | main | Eurasia | 51.48200 | -3.621000 | OXB522-15 | NA | United Kingdom |
| OXBTGS769 | main | Eurasia | 51.53600 | -3.757000 | OXB516-15 | NA | United Kingdom |
| OXBTGS770 | main | Eurasia | 51.53600 | -3.757000 | OXB517-15 | NA | United Kingdom |
| OXBTGS541 | main | Eurasia | 51.68000 | -4.270000 | OXB488-15 | NA | United Kingdom |
| OXBTGS542 | main | Eurasia | 51.68000 | -4.270000 | OXB489-15 | NA | United Kingdom |
| OXBTGS518 | main | Eurasia | 51.73000 | -0.803000 | OXB486-15 | NA | United Kingdom |
| OXBTGS519 | main | Eurasia | 51.73000 | -0.803000 | OXB487-15 | NA | United Kingdom |
| OXBTGS1332 | Britain | Britain | 51.73890 | -0.798920 | OXB1614-16 | NA | United Kingdom |
| RVcoll16I288 | main | Eurasia | 51.95240 | 21.981900 | EULEP4960-16 | NA | Poland |
| RVcoll14I860 | main | Eurasia | 52.14700 | 21.200000 | EULEP2023-15 | NA | Poland |
| RVcoll12Z115 | main | Eurasia | 52.25300 | -8.503000 | WMB4026-14 | NA | Ireland |
| BCZSMLep82863 | main | Eurasia | 52.30110 | 13.261400 | NA | NA | Germany |
| RVcoll16I578 | main | Eurasia | 53.15890 | 17.823700 | EULEP4981-16 | NA | Poland |
| RVcoll16I161 | main | Eurasia | 53.20600 | 11.345900 | EULEP4945-16 | NA | Germany |
| GBLAA1061-15 | main | Eurasia | 54.05500 | 9.594000 | GBLAA1061-15 | MH419694 | Germany |
| RVcoll15Q070 | main | Eurasia | 54.41000 | 38.510000 | EULEP4536-16 | NA | Russian Federation |
| RVcoll12Z169 | main | Eurasia | 54.45000 | -8.448000 | WMB4034-14 | NA | Ireland |
| OXBTGS933 | main | Eurasia | 54.50000 | -2.730000 | OXB558-15 | NA | United Kingdom |
| OXBTGS934 | main | Eurasia | 54.50000 | -2.730000 | OXB559-15 | NA | United Kingdom |
| OXBTGS875 | main | Eurasia | 54.64100 | -1.184000 | OXB529-15 | NA | United Kingdom |

Table S2: Archived and newly generated *Polyommatus icarus* sequences used for CO1 mtDNA analysis *(continued)*

| Sample ID | Island | Geographic region | Latitude (dec. deg.) | Longitude (dec. deg.) | BOLD Process ID | GenBank Accession | Country |
|---|---|---|---|---|---|---|---|
| OXBTGS876 | main | Eurasia | 54.64100 | -1.184000 | OXB530-15 | NA | United Kingdom |
| OXBTGS877 | main | Eurasia | 54.64100 | -1.184000 | OXB531-15 | NA | United Kingdom |
| OXBTGS861 | main | Eurasia | 54.69000 | -1.491000 | OXB527-15 | NA | United Kingdom |
| OXBTGS862 | main | Eurasia | 54.69000 | -1.491000 | OXB528-15 | NA | United Kingdom |
| EULEP362-14 | main | Eurasia | 54.90000 | 24.230000 | EULEP362-14 | MM23847 | Lithuania |
| RVcoll12R452 | main | Eurasia | 55.18700 | -4.918000 | WMB4019-14 | NA | United Kingdom |
| RVcoll15Q093 | main | Eurasia | 55.55370 | 38.875100 | EULEP4540-16 | NA | Russian Federation |
| RVcoll15Q094 | main | Eurasia | 55.55370 | 38.875100 | EULEP4541-16 | NA | Russian Federation |
| OXBTGS159 | main | Eurasia | 56.32000 | -5.590000 | OXB435-15 | NA | United Kingdom |
| OXBTGS160 | main | Eurasia | 56.32000 | -5.590000 | OXB436-15 | NA | United Kingdom |
| OXBTGS161 | main | Eurasia | 56.32000 | -5.590000 | OXB437-15 | NA | United Kingdom |
| OXBTGS017 | main | Eurasia | 56.39200 | -5.506000 | OXB387-15 | NA | United Kingdom |
| OXBTGS018 | main | Eurasia | 56.39200 | -5.506000 | OXB388-15 | NA | United Kingdom |
| OXBTGS019 | main | Eurasia | 56.39200 | -5.506000 | OXB389-15 | NA | United Kingdom |
| OXBTGS020 | main | Eurasia | 56.39200 | -5.506000 | OXB390-15 | NA | United Kingdom |
| OXBTGS925 | main | Eurasia | 56.42000 | -5.750000 | OXB550-15 | NA | United Kingdom |
| OXBTGS926 | main | Eurasia | 56.42000 | -5.750000 | OXB551-15 | NA | United Kingdom |
| OXBTGS927 | main | Eurasia | 56.42000 | -5.750000 | OXB552-15 | NA | United Kingdom |
| OXBTGS928 | main | Eurasia | 56.42000 | -5.750000 | OXB553-15 | NA | United Kingdom |
| OXBTGS929 | main | Eurasia | 56.42000 | -5.750000 | OXB554-15 | NA | United Kingdom |
| OXBTGS918 | main | Eurasia | 56.58000 | -5.740000 | OXB543-15 | NA | United Kingdom |
| OXBTGS919 | main | Eurasia | 56.71000 | -5.280000 | OXB544-15 | NA | United Kingdom |
| RVcoll12R455 | main | Eurasia | 56.99300 | -5.824000 | WMB4020-14 | NA | United Kingdom |
| RVcoll08L303 | main | Eurasia | 57.13000 | 10.010000 | GBMIN32546-13 | JN084695 | Denmark |
| OXBTGS035 | outer Hebrides | Eurasia | 57.21700 | -7.423000 | OXB396-15 | NA | United Kingdom |
| OXBTGS061 | outer Hebrides | Eurasia | 57.30600 | -7.397000 | OXB397-15 | NA | United Kingdom |
| OXBTGS062 | outer Hebrides | Eurasia | 57.30600 | -7.397000 | OXB398-15 | NA | United Kingdom |
| OXBTGS100 | outer Hebrides | Eurasia | 57.65700 | -7.371000 | OXB418-15 | NA | United Kingdom |
| OXBTGS101 | outer Hebrides | Eurasia | 57.72700 | -7.195000 | OXB419-15 | NA | United Kingdom |
| OXBTGS102 | outer Hebrides | Eurasia | 57.72700 | -7.195000 | OXB420-15 | NA | United Kingdom |
| OXBTGS103 | outer Hebrides | Eurasia | 57.72700 | -7.195000 | OXB421-15 | NA | United Kingdom |
| OXBTGS104 | outer Hebrides | Eurasia | 57.72700 | -7.195000 | OXB422-15 | NA | United Kingdom |
| OXBTGS105 | outer Hebrides | Eurasia | 57.72700 | -7.195000 | OXB423-15 | NA | United Kingdom |
| EULEP335-14 | main | Eurasia | 59.35700 | 24.626300 | EULEP335-14 | MM23820 | Estonia |
| LON963-12 | main | Eurasia | 59.39000 | 10.518000 | LON963-12 | KX049307 | Norway |
| LON231-08 | main | Eurasia | 59.89330 | 10.737200 | LON231-08 | KX049168 | Norway |
| RVcoll07C710 | main | Eurasia | 60.44000 | 26.190000 | GBMIN32570-13 | JN084694 | Finland |
| LEFIE954-10 | main | Eurasia | 62.54200 | 29.526000 | LEFIE954-10 | HM874671-SUPPRESSED | Finland |
| RVcoll16G983 | main | Eurasia | 63.88345 | 15.682640 | EULEP4756-16 | NA | Sweden |
| LON634-09 | main | Eurasia | 64.48080 | 13.661900 | LON634-09 | KX047754 | Norway |
| LEFIJ522-10 | main | Eurasia | 64.79700 | 25.317000 | LEFIJ522-10 | JF853641-SUPPRESSED | Finland |
| LEFIJ521-10 | main | Eurasia | 69.31200 | 25.732000 | LEFIJ521-10 | KM572032 | Finland |
| RVcoll16C054 | main | Eurasia | 44.90700 | 4.723000 | BIBSA2352-20 | NA | France |
| RVcoll20A044 | main | Eurasia | 40.59400 | 14.379000 | NA | NA | Italy |
| MABUT045-10 | main | Eurasia | 35.46670 | 72.583300 | MABUT045-10 | HQ990363 | Pakistan |
| MABUT046-10 | main | Eurasia | 33.90000 | 73.383300 | MABUT046-10 | HQ990364 | Pakistan |

Table S2: Archived and newly generated *Polyommatus icarus* sequences used for CO1 mtDNA analysis *(continued)*

| **Sample ID** | **Island** | **Geographic region** | **Latitude (dec. deg.)** | **Longitude (dec. deg.)** | **BOLD Process ID** | **GenBank Accession** | **Country** |
|---|---|---|---:|---:|---|---|---|
| MABUT126-10 | main | Eurasia | 33.90000 | 73.383300 | MABUT126-10 | HQ990435 | Pakistan |
| LOWA002-06 | main | Eurasia | 49.63300 | 83.567000 | LOWA002-06 | FJ663998 | Kazakhstan |
| LOWA001-06 | main | Eurasia | 49.63300 | 83.567000 | LOWA001-06 | FJ663999 | Kazakhstan |
| OXB1552-16 | main | Eurasia | 42.57780 | 1.666050 | OXB1552-16 | NA | Andorra |
| OXB1553-16 | main | Eurasia | 42.57780 | 1.666050 | OXB1553-16 | NA | Andorra |
| OXB1589-16 | main | Eurasia | 44.79950 | 5.259680 | OXB1589-16 | NA | France |
| OXB1590-16 | main | Eurasia | 44.44470 | 5.210590 | OXB1590-16 | NA | France |
| OXB1614-16 | main | Eurasia | 51.73890 | -0.798920 | OXB1614-16 | NA | United Kingdom |
| CNCBF1012-14 | main | America | 45.68900 | -74.089000 | CNCBF1012-14 | NA | Canada |
| CNCBF652-14 | main | America | 45.68900 | -74.089000 | CNCBF652-14 | NA | Canada |
| CNCBF653-14 | main | America | 45.68900 | -74.089000 | CNCBF653-14 | NA | Canada |
| EZBNA882-07 | main | America | 45.68900 | -74.089000 | EZBNA882-07 | NA | Canada |
| EZBNA884-07 | main | America | 45.68900 | -74.089000 | EZBNA884-07 | NA | Canada |
| EZBNA885-07 | main | America | 45.68900 | -74.089000 | EZBNA885-07 | NA | Canada |
| EZBNA886-07 | main | America | 45.68900 | -74.089000 | EZBNA886-07 | NA | Canada |
| EZBNA887-07 | main | America | 45.68900 | -74.089000 | EZBNA887-07 | NA | Canada |
| EZHBA247-07 | main | Eurasia | 45.43700 | 5.856000 | EZHBA247-07 | NA | France |
| EZHBA353-07 | main | Eurasia | 48.00000 | 7.700000 | EZHBA353-07 | NA | Germany |
| EZHBA539-07 | main | Eurasia | 54.98000 | 82.890000 | EZHBA539-07 | NA | Russia |
| EZHBA540-07 | main | Eurasia | 54.98000 | 82.890000 | EZHBA540-07 | NA | Russia |
| EZHBA710-07 | main | Eurasia | 48.00000 | 7.700000 | EZHBA710-07 | NA | Germany |
| EZHBA711-07 | main | Eurasia | 48.00000 | 7.700000 | EZHBA711-07 | NA | Germany |
| IRANB381-08 | main | Eurasia | 36.12000 | 51.200000 | IRANB381-08 | NA | Iran |
| IRANB382-08 | main | Eurasia | 36.12000 | 51.200000 | IRANB382-08 | NA | Iran |
| IRANB390-08 | main | Eurasia | 36.15000 | 51.300000 | IRANB390-08 | NA | Iran |
| IRANB403-08 | main | Eurasia | 38.58300 | 44.367000 | IRANB403-08 | NA | Iran |
| IRANB414-08 | main | Eurasia | 34.59300 | 47.086000 | IRANB414-08 | NA | Iran |
| IRANB415-08 | main | Eurasia | 34.59300 | 47.086000 | IRANB415-08 | NA | Iran |
| IRANB416-08 | main | Eurasia | 34.59300 | 47.086000 | IRANB416-08 | NA | Iran |
| LOWAB113-07 | main | Eurasia | 40.08300 | 44.917000 | LOWAB113-07 | NA | Armenia |
| LOWAB167-09 | main | Eurasia | 40.84030 | 41.159700 | LOWAB167-09 | NA | Turkey |
| LOWAB287-09 | main | Eurasia | 40.52640 | 41.973100 | LOWAB287-09 | NA | Turkey |
| NLLEA1447-14 | main | Eurasia | 52.17000 | 4.500000 | NLLEA1447-14 | NA | Netherlands |
| NLLEA1480-14 | main | Eurasia | 52.33000 | 4.550000 | NLLEA1480-14 | NA | Netherlands |
| NLLEA1500-14 | main | Eurasia | 52.07700 | 5.565000 | NLLEA1500-14 | NA | Netherlands |
| NLLEA1503-14 | main | Eurasia | 52.52700 | 4.959000 | NLLEA1503-14 | NA | Netherlands |
| BERf30 | outer Hebrides | Britain | 57.71385 | -7.213534 | NA | MT151364 | United Kingdom |
| BERf35 | outer Hebrides | Britain | 57.71385 | -7.213534 | NA | MT151355 | United Kingdom |
| BERf37 | outer Hebrides | Britain | 57.71385 | -7.213534 | NA | MT151363 | United Kingdom |
| BERf43 | outer Hebrides | Britain | 57.71385 | -7.213534 | NA | MW394241 | United Kingdom |
| BERm27 | outer Hebrides | Britain | 57.71385 | -7.213534 | NA | MW394242 | United Kingdom |
| BERm28 | outer Hebrides | Britain | 57.71385 | -7.213534 | NA | MW394243 | United Kingdom |
| BERm29 | outer Hebrides | Britain | 57.71385 | -7.213534 | NA | MW394244 | United Kingdom |
| BERm36 | outer Hebrides | Britain | 57.71385 | -7.213534 | NA | MT151362 | United Kingdom |
| BERm40 | outer Hebrides | Britain | 57.71385 | -7.213534 | NA | MT151356 | United Kingdom |
| BERm42 | outer Hebrides | Britain | 57.71385 | -7.213534 | NA | MT151361 | United Kingdom |

Table S2: Archived and newly generated *Polyommatus icarus* sequences used for CO1 mtDNA analysis *(continued)*

| **Sample ID** | **Island** | **Geographic region** | **Latitude (dec. deg.)** | **Longitude (dec. deg.)** | **BOLD Process ID** | **GenBank Accession** | **Country** |
|---|---|---|---:|---:|---|---|---|
| BERm45 | outer Hebrides | Britain | 57.71385 | -7.213534 | NA | MT151365 | United Kingdom |
| BMDf149 | Britain | Britain | 51.78256 | -1.125439 | NA | MW394245 | United Kingdom |
| BMDf154 | Britain | Britain | 51.78256 | -1.125439 | NA | MW394246 | United Kingdom |
| BMDf270 | Britain | Britain | 51.78256 | -1.125439 | NA | MW394247 | United Kingdom |
| BMDf274 | Britain | Britain | 51.78256 | -1.125439 | NA | MW394248 | United Kingdom |
| BMDf274 | Britain | Britain | 51.78256 | -1.125439 | NA | MW394249 | United Kingdom |
| BMDf276 | Britain | Britain | 51.78256 | -1.125439 | NA | MW394250 | United Kingdom |
| BMDf279 | Britain | Britain | 51.78256 | -1.125439 | NA | MW394251 | United Kingdom |
| BMDm147 | Britain | Britain | 51.78256 | -1.125439 | NA | MW394252 | United Kingdom |
| BMDm148 | Britain | Britain | 51.78256 | -1.125439 | NA | MW394253 | United Kingdom |
| BMDm150 | Britain | Britain | 51.78256 | -1.125439 | NA | MW394254 | United Kingdom |
| BMDm151 | Britain | Britain | 51.78256 | -1.125439 | NA | MW394255 | United Kingdom |
| BMDm152 | Britain | Britain | 51.78256 | -1.125439 | NA | MW394256 | United Kingdom |
| BMDm153 | Britain | Britain | 51.78256 | -1.125439 | NA | MW394257 | United Kingdom |
| BWDf3 | Britain | Britain | 50.57975 | -3.898247 | NA | MW394258 | United Kingdom |
| BWDf5 | Britain | Britain | 50.57975 | -3.898247 | NA | MW394259 | United Kingdom |
| BWDm1 | Britain | Britain | 50.57975 | -3.898247 | NA | MW394260 | United Kingdom |
| BWDm2 | Britain | Britain | 50.57975 | -3.898247 | NA | MW394261 | United Kingdom |
| BWDm4 | Britain | Britain | 50.57975 | -3.898247 | NA | MW394262 | United Kingdom |
| BWDm6 | Britain | Britain | 50.57975 | -3.898247 | NA | MW394263 | United Kingdom |
| BWDm7 | Britain | Britain | 50.57975 | -3.898247 | NA | MW394264 | United Kingdom |
| DGCf87 | Britain | Britain | 57.87810 | -4.016982 | NA | MT151348 | United Kingdom |
| DGCm84 | Britain | Britain | 57.87810 | -4.016982 | NA | MT151339 | United Kingdom |
| DGCm86 | Britain | Britain | 57.87810 | -4.016982 | NA | MT151338 | United Kingdom |
| DGCm94 | Britain | Britain | 57.87810 | -4.016982 | NA | MT151340 | United Kingdom |
| DGCm99 | Britain | Britain | 57.87810 | -4.016982 | NA | MW394273 | United Kingdom |
| DGCm100 | Britain | Britain | 57.87810 | -4.016982 | NA | MT151349 | United Kingdom |
| ETBf203 | Britain | Britain | 50.76919 | 0.243747 | NA | MW394274 | United Kingdom |
| ETBm188 | Britain | Britain | 50.76919 | 0.243747 | NA | MW394275 | United Kingdom |
| ETBm189 | Britain | Britain | 50.76919 | 0.243747 | NA | MW394276 | United Kingdom |
| ETBm190 | Britain | Britain | 50.76919 | 0.243747 | NA | MW394277 | United Kingdom |
| ETBm191 | Britain | Britain | 50.76919 | 0.243747 | NA | MW394278 | United Kingdom |
| ETBm199 | Britain | Britain | 50.76919 | 0.243747 | NA | MW394279 | United Kingdom |
| ETBm200 | Britain | Britain | 50.76919 | 0.243747 | NA | MW394280 | United Kingdom |
| ETBm201 | Britain | Britain | 50.76919 | 0.243747 | NA | MW394281 | United Kingdom |
| ETBm202 | Britain | Britain | 50.76919 | 0.243747 | NA | MW394282 | United Kingdom |
| ETBm205 | Britain | Britain | 50.76919 | 0.243747 | NA | MW394283 | United Kingdom |
| FRNfM02 | Britain | Britain | 44.15734 | 1.982669 | NA | MW394284 | France |
| FRNfM04 | main | Eurasia | 44.15734 | 1.982669 | NA | MW394285 | France |
| FRNmM03 | main | Eurasia | 44.15734 | 1.982669 | NA | MW394286 | France |
| FRNmM05 | main | Eurasia | 44.15734 | 1.982669 | NA | MW394287 | France |
| FRNmM06 | main | Eurasia | 44.15734 | 1.982669 | NA | MW394288 | France |
| MDCf135 | Britain | Britain | 53.29568 | -3.843251 | NA | MW394289 | United Kingdom |
| MDCf136 | Britain | Britain | 53.29568 | -3.843251 | NA | MW394290 | United Kingdom |
| MDCf138 | Britain | Britain | 53.29568 | -3.843251 | NA | MW394291 | United Kingdom |
| MDCf139 | Britain | Britain | 53.29568 | -3.843251 | NA | MW394292 | United Kingdom |

Table S2: Archived and newly generated *Polyommatus icarus* sequences used for CO1 mtDNA analysis *(continued)*

| **Sample ID** | **Island** | **Geographic region** | **Latitude (dec. deg.)** | **Longitude (dec. deg.)** | **BOLD Process ID** | **GenBank Accession** | **Country** |
|---|---|---|---:|---:|---|---|---|
| MDCf141 | Britain | Britain | 53.29568 | -3.843251 | NA | MW394295 | United Kingdom |
| MDCf142 | Britain | Britain | 53.29568 | -3.843251 | NA | MW394293 | United Kingdom |
| MDCf144 | Britain | Britain | 53.29568 | -3.843251 | NA | MW394294 | United Kingdom |
| MDCm145 | Britain | Britain | 53.29568 | -3.843251 | NA | MW394296 | United Kingdom |
| MLGf3 | Britain | Britain | 56.99196 | -5.834874 | NA | MW394297 | United Kingdom |
| MLGf5 | Britain | Britain | 56.99196 | -5.834874 | NA | MT151347 | United Kingdom |
| MLGf8 | Britain | Britain | 56.99196 | -5.834874 | NA | MW394298 | United Kingdom |
| MLGf10 | Britain | Britain | 56.99196 | -5.834874 | NA | MT151352 | United Kingdom |
| MLGf11 | Britain | Britain | 56.99196 | -5.834874 | NA | MT151341 | United Kingdom |
| MLGm1 | Britain | Britain | 56.99196 | -5.834874 | NA | MW394299 | United Kingdom |
| MLGm2 | Britain | Britain | 56.99196 | -5.834874 | NA | MT151353 | United Kingdom |
| MLGm4 | Britain | Britain | 56.99196 | -5.834874 | NA | MW394300 | United Kingdom |
| MLGm6 | Britain | Britain | 56.99196 | -5.834874 | NA | MW394301 | United Kingdom |
| MLGm9 | Britain | Britain | 56.99196 | -5.834874 | NA | MW394302 | United Kingdom |
| MLGm14 | Britain | Britain | 56.99196 | -5.834874 | NA | MT151350 | United Kingdom |
| MLGm17 | Britain | Britain | 56.99196 | -5.834874 | NA | MT151351 | United Kingdom |
| MMSf211 | Britain | Britain | 52.16802 | 1.255096 | NA | MW394303 | United Kingdom |
| MMSf215 | Britain | Britain | 52.16802 | 1.255096 | NA | MW394304 | United Kingdom |
| MMSf217 | Britain | Britain | 52.16802 | 1.255096 | NA | MW394305 | United Kingdom |
| MMSf221 | Britain | Britain | 52.16802 | 1.255096 | NA | MW394306 | United Kingdom |
| MMSf222 | Britain | Britain | 52.16802 | 1.255096 | NA | MW394307 | United Kingdom |
| MMSm207 | Britain | Britain | 52.16802 | 1.255096 | NA | MW394308 | United Kingdom |
| MMSm208 | Britain | Britain | 52.16802 | 1.255096 | NA | MW394309 | United Kingdom |
| MMSm209 | Britain | Britain | 52.16802 | 1.255096 | NA | MW394310 | United Kingdom |
| MMSm210 | Britain | Britain | 52.16802 | 1.255096 | NA | MW394311 | United Kingdom |
| MMSm212 | Britain | Britain | 52.16802 | 1.255096 | NA | MW394312 | United Kingdom |
| MMSm213 | Britain | Britain | 52.16802 | 1.255096 | NA | MW394313 | United Kingdom |
| MMSm214 | Britain | Britain | 52.16802 | 1.255096 | NA | MW394314 | United Kingdom |
| MMSm220 | Britain | Britain | 52.16802 | 1.255096 | NA | MW394315 | United Kingdom |
| OBNf111 | Britain | Britain | 56.43357 | -5.482266 | NA | MW394316 | United Kingdom |
| OBNf121 | Britain | Britain | 56.43357 | -5.482266 | NA | MT151342 | United Kingdom |
| OBNm110 | Britain | Britain | 56.43357 | -5.482266 | NA | MT151354 | United Kingdom |
| OBNm112 | Britain | Britain | 56.43357 | -5.482266 | NA | MT151345 | United Kingdom |
| OBNm113 | Britain | Britain | 56.43357 | -5.482266 | NA | MT151344 | United Kingdom |
| OBNm115 | Britain | Britain | 56.43357 | -5.482266 | NA | MW394317 | United Kingdom |
| OBNm116 | Britain | Britain | 56.43357 | -5.482266 | NA | MW394318 | United Kingdom |
| OBNm117 | Britain | Britain | 56.43357 | -5.482266 | NA | MT151343 | United Kingdom |
| OBNm118 | Britain | Britain | 56.43357 | -5.482266 | NA | MW394319 | United Kingdom |
| OBNm119 | Britain | Britain | 56.43357 | -5.482266 | NA | MW394320 | United Kingdom |
| OBNm120 | Britain | Britain | 56.43357 | -5.482266 | NA | MW394321 | United Kingdom |
| PCPf175 | Britain | Britain | 51.67422 | -4.308992 | NA | MW394322 | United Kingdom |
| PCPm156 | Britain | Britain | 51.67422 | -4.308992 | NA | MW394323 | United Kingdom |
| PCPm157 | Britain | Britain | 51.67422 | -4.308992 | NA | MW394324 | United Kingdom |
| PCPm158 | Britain | Britain | 51.67422 | -4.308992 | NA | MW394325 | United Kingdom |
| PCPm159 | Britain | Britain | 51.67422 | -4.308992 | NA | MW394326 | United Kingdom |
| PCPm160 | Britain | Britain | 51.67422 | -4.308992 | NA | MW394327 | United Kingdom |

Table S2: Archived and newly generated *Polyommatus icarus* sequences used for CO1 mtDNA analysis *(continued)*

| **Sample ID** | **Island** | **Geographic region** | **Latitude (dec. deg.)** | **Longitude (dec. deg.)** | **BOLD Process ID** | **GenBank Accession** | **Country** |
|---|---|---|---:|---:|---|---|---|
| PCPm172 | Britain | Britain | 51.67422 | -4.308992 | NA | MW394328 | United Kingdom |
| PCPm176 | Britain | Britain | 51.67422 | -4.308992 | NA | MW394329 | United Kingdom |
| PCPm177 | Britain | Britain | 51.67422 | -4.308992 | NA | MW394330 | United Kingdom |
| RHDf242 | Britain | Britain | 54.71391 | -1.477433 | NA | MW394331 | United Kingdom |
| RHDf245 | Britain | Britain | 54.71391 | -1.477433 | NA | MW394332 | United Kingdom |
| RHDf253 | Britain | Britain | 54.71391 | -1.477433 | NA | MW394333 | United Kingdom |
| RHDm239 | Britain | Britain | 54.71391 | -1.477433 | NA | MW394334 | United Kingdom |
| RHDm240 | Britain | Britain | 54.71391 | -1.477433 | NA | MW394335 | United Kingdom |
| RHDm241 | Britain | Britain | 54.71391 | -1.477433 | NA | MW394336 | United Kingdom |
| RHDm247 | Britain | Britain | 54.71391 | -1.477433 | NA | MW394337 | United Kingdom |
| RHDm254 | Britain | Britain | 54.71391 | -1.477433 | NA | MW394338 | United Kingdom |
| RHDm255 | Britain | Britain | 54.71391 | -1.477433 | NA | MW394339 | United Kingdom |
| RHDm256 | Britain | Britain | 54.71391 | -1.477433 | NA | MW394340 | United Kingdom |
| RHDm257 | Britain | Britain | 54.71391 | -1.477433 | NA | MT151346 | United Kingdom |
| CFWm223 | Britain | Britain | 53.25443 | -0.281520 | NA | MW394265 | United Kingdom |
| CFWm225 | Britain | Britain | 53.25443 | -0.281520 | NA | MW394266 | United Kingdom |
| CFWm226 | Britain | Britain | 53.25443 | -0.281520 | NA | MW394267 | United Kingdom |
| CFWm229 | Britain | Britain | 53.25443 | -0.281520 | NA | MW394268 | United Kingdom |
| CFWm230 | Britain | Britain | 53.25443 | -0.281520 | NA | MW394269 | United Kingdom |
| CFWm231 | Britain | Britain | 53.25443 | -0.281520 | NA | MW394270 | United Kingdom |
| CFWm233 | Britain | Britain | 53.25443 | -0.281520 | NA | MW394271 | United Kingdom |
| CFWm234 | Britain | Britain | 53.25443 | -0.281520 | NA | MW394272 | United Kingdom |
| RVSfR3 | Britain | Britain | 56.02397 | -2.596053 | NA | MW394341 | United Kingdom |
| RVSfR4 | Britain | Britain | 56.02397 | -2.596053 | NA | MW394342 | United Kingdom |
| RVSmR1 | Britain | Britain | 56.02397 | -2.596053 | NA | MW394343 | United Kingdom |
| RVSmR2 | Britain | Britain | 56.02397 | -2.596053 | NA | MW394344 | United Kingdom |
| RVSmR5 | Britain | Britain | 56.02397 | -2.596053 | NA | MW394345 | United Kingdom |
| RVSmR1 | Britain | Britain | 56.02397 | -2.596053 | NA | MW394346 | United Kingdom |
| RVSmR6 | Britain | Britain | 56.02397 | -2.596053 | NA | MT151366 | United Kingdom |
| TULf72 | outer Hebrides | Britain | 58.18553 | -7.025334 | NA | MT151360 | United Kingdom |
| TULf73 | outer Hebrides | Britain | 58.18553 | -7.025334 | NA | MT151368 | United Kingdom |
| TULf77 | outer Hebrides | Britain | 58.18553 | -7.025334 | NA | MW394347 | United Kingdom |
| TULm53 | outer Hebrides | Britain | 58.18553 | -7.025334 | NA | MT151367 | United Kingdom |
| TULm55 | outer Hebrides | Britain | 58.18553 | -7.025334 | NA | MT151359 | United Kingdom |
| TULm60 | outer Hebrides | Britain | 58.18553 | -7.025334 | NA | MW394348 | United Kingdom |
| TULm62 | outer Hebrides | Britain | 58.18553 | -7.025334 | NA | MT151358 | United Kingdom |
| TULm63 | outer Hebrides | Britain | 58.18553 | -7.025334 | NA | MW394349 | United Kingdom |
| TULm71 | outer Hebrides | Britain | 58.18553 | -7.025334 | NA | MT151357 | United Kingdom |

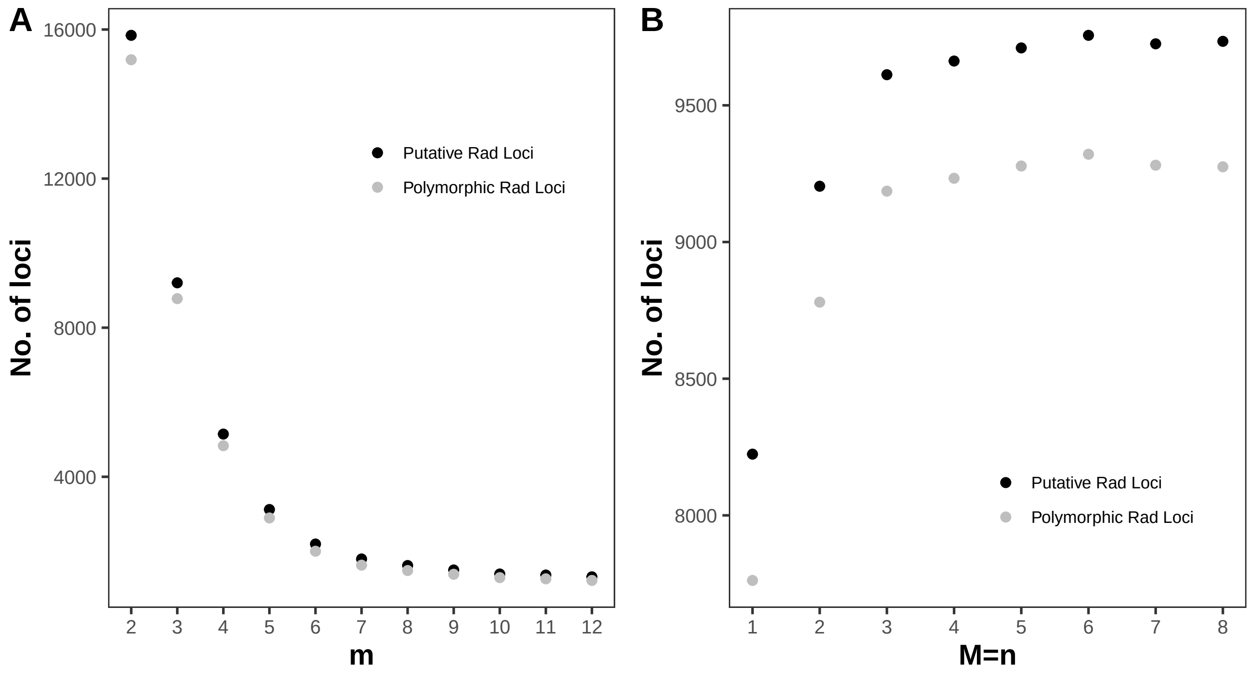


**Figure S2** Tests of different combinations of Stacks parameters **m (A)** and **M=n (B)** on assembly of total and polymorphic RAD loci.

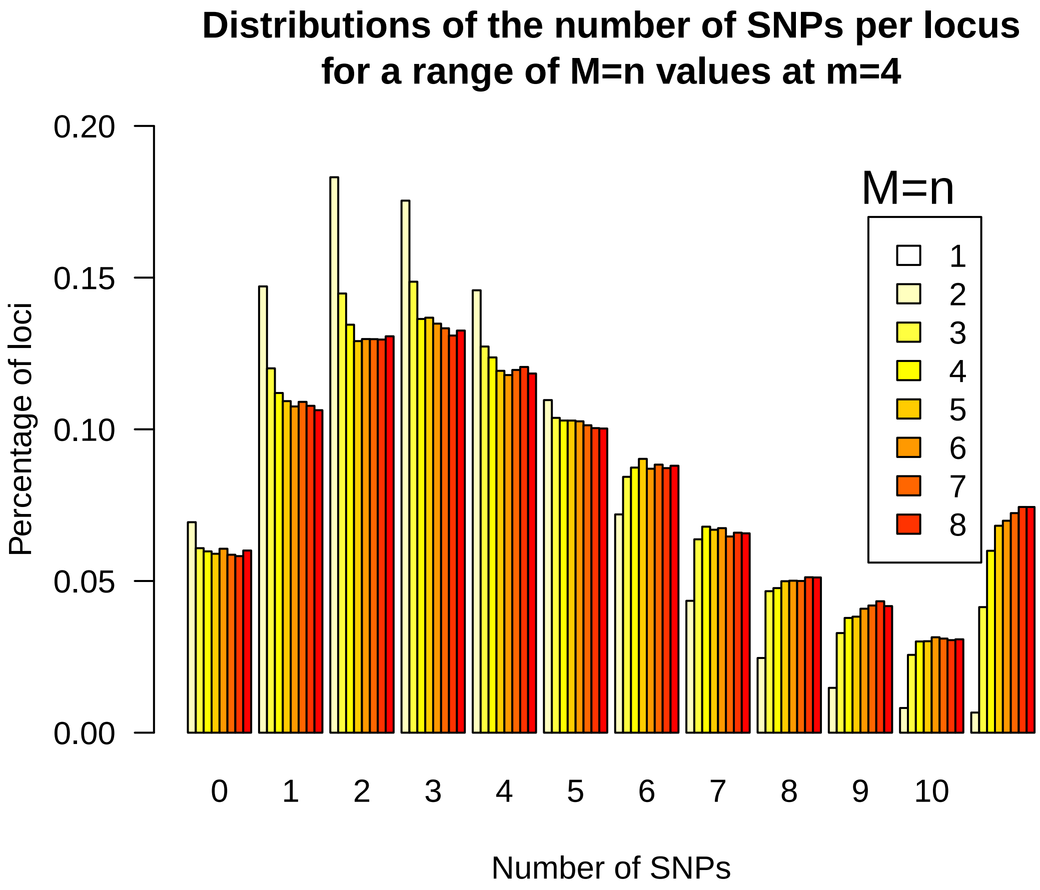


**Figure S3** Tests of different combination of Stacks parameters **M**=**n** for **m**=4 on the disribution of SNPs across loci.

Table S3: Summary of RAD Loci assembly using Stacks

| **Sample** | **SRA Accession** | **Total Reads** | **POP** | **Average Coverage of Stacks** | **Standard Deviation** | **No. of Reads Used** |
|---|---|---|---|---|---|---|
| BERf030 | SRR11238036 | 3129338 | BER | 50.31 | 175.75 | 1395803(44.6%) |
| BERf035 | SRR11238036 | 3317737 | BER | 57.68 | 194.96 | 1379735(41.6%) |
| BERf037 | SRR11238036 | 3695537 | BER | 43.18 | 158.72 | 1435221(38.8%) |
| BERf043 | SRR11238036 | 3071772 | BER | 60.87 | 200.79 | 1295991(42.2%) |
| BERm027 | SRR11238036 | 3195023 | BER | 55.42 | 188.97 | 1307905(40.9%) |
| BERm028 | SRR11238036 | 6952171 | BER | 39.15 | 145.69 | 1656005(23.8%) |
| BERm029 | SRR11238036 | 8470130 | BER | 37.75 | 137.99 | 1769026(20.9%) |
| BERm036 | SRR11238036 | 3855171 | BER | 49.6 | 178.65 | 1456264(37.8%) |
| BERm038 | SRR11238036 | 2859523 | BER | 54.79 | 182.72 | 1306837(45.7%) |
| BERm039 | SRR11238036 | 2149488 | BER | 71.11 | 200.67 | 1027671(47.8%) |
| BERm040 | SRR11238036 | 3457223 | BER | 52.38 | 183.1 | 1387460(40.1%) |
| BERm041 | SRR11238036 | 1436794 | BER | 81.07 | 184.1 | 772559(53.8%) |
| BERm042 | SRR11238036 | 3264460 | BER | 51.94 | 178.12 | 1265899(38.8%) |
| BERm044 | SRR11238036 | 3122980 | BER | 66.41 | 205.14 | 1273322(40.8%) |
| BERm045 | SRR11238036 | 3168433 | BER | 49.44 | 171.94 | 1325130(41.8%) |
| BERm046 | SRR11238036 | 3620747 | BER | 47.17 | 173.38 | 1374133(38.0%) |
| BMDf149 | SRR11238035 | 3713282 | BMD | 37.51 | 124.13 | 1742297(46.9%) |
| BMDf154 | SRR11238035 | 3984391 | BMD | 39.01 | 129.17 | 1874337(47.0%) |
| BMDf270 | SRR11238035 | 2961330 | BMD | 34.51 | 104.28 | 1473334(49.8%) |
| BMDf272 | SRR11238035 | 3200952 | BMD | 32.74 | 104.73 | 1597267(49.9%) |
| BMDf274 | SRR11238035 | 2556011 | BMD | 29.1 | 87.06 | 1181892(46.2%) |
| BMDf276 | SRR11238035 | 2740520 | BMD | 29.3 | 83.73 | 1362646(49.7%) |
| BMDf278 | SRR11238035 | 1893829 | BMD | 35.29 | 103.17 | 857798(45.3%) |
| BMDf279 | SRR11238035 | 3436779 | BMD | 33.29 | 107.73 | 1680352(48.9%) |
| BMDm147 | SRR11238035 | 3537215 | BMD | 38.25 | 126.23 | 1771228(50.1%) |
| BMDm148 | SRR11238035 | 3441037 | BMD | 37.82 | 121.45 | 1695779(49.3%) |
| BMDm150 | SRR11238035 | 3947571 | BMD | 34.31 | 114.36 | 1986204(50.3%) |
| BMDm151 | SRR11238035 | 3500869 | BMD | 36.55 | 116.57 | 1792859(51.2%) |
| BMDm152 | SRR11238035 | 3707329 | BMD | 37.31 | 121.18 | 1733345(46.8%) |
| BMDm153 | SRR11238035 | 3620544 | BMD | 37.27 | 115.17 | 1669465(46.1%) |
| BMDm271 | SRR11238035 | 1723401 | BMD | 40.73 | 96.16 | 847625(49.2%) |
| BMDm275 | SRR11238035 | 1390969 | BMD | 32.82 | 88.13 | 689129(49.5%) |
| BWDf3 | SRR11238035 | 4621287 | BWD | 37.19 | 115.73 | 2277809(49.3%) |
| BWDf5 | SRR11238035 | 4042362 | BWD | 38.4 | 123.28 | 1979966(49.0%) |
| BWDm1 | SRR11238035 | 4163695 | BWD | 35.64 | 106.9 | 2041968(49.0%) |
| BWDm2 | SRR11238035 | 4653162 | BWD | 36.64 | 106.19 | 2354239(50.6%) |
| BWDm4 | SRR11238035 | 4663845 | BWD | 37.78 | 110.85 | 2271000(48.7%) |
| BWDm6 | SRR11238035 | 4508000 | BWD | 37.58 | 117.53 | 2150872(47.7%) |
| BWDm7 | SRR11238035 | 4181443 | BWD | 36.49 | 113.84 | 2160048(51.7%) |
| CFWm223 | SRR11238035 | 2672555 | CFW | 34.91 | 107.73 | 1379267(51.6%) |

Table S3: Summary of RAD Loci assembly using Stacks *(continued)*

| **Sample** | **SRA Accession** | **Total Reads** | **POP** | **Average Coverage of Stacks** | **Standard Deviation** | **No. of Reads Used** |
|---|---|---|---|---|---|---|
| CFWm224 | SRR11238035 | 931489 | CFW | 33.02 | 80.62 | 473217(50.8%) |
| CFWm225 | SRR11238035 | 184682 | CFW | n/a | n/a | n/a |
| CFWm226 | SRR11238035 | 2763326 | CFW | 36.66 | 114.4 | 1369747(49.6%) |
| CFWm227 | SRR11238035 | 585059 | CFW | 26.38 | 57.71 | 271176(46.4%) |
| CFWm228 | SRR11238035 | 2217151 | CFW | 33.67 | 106.71 | 1064328(48.0%) |
| CFWm229 | SRR11238035 | 2446267 | CFW | 32.52 | 90.83 | 1237435(50.6%) |
| CFWm230 | SRR11238035 | 2313054 | CFW | 30.33 | 93.79 | 1141870(49.4%) |
| CFWm231 | SRR11238035 | 2292118 | CFW | 33.33 | 107.58 | 1174822(51.3%) |
| CFWm232 | SRR11238035 | 256147 | CFW | n/a | n/a | n/a |
| CFWm233 | SRR11238035 | 2173984 | CFW | 30.64 | 100.68 | 1097179(50.5%) |
| CFWm234 | SRR11238035 | 2793732 | CFW | 32.94 | 110.1 | 1414162(50.6%) |
| DGCf087 | SRR11238036 | 3347527 | DGC | 54.93 | 189.48 | 1471911(44.0%) |
| DGCf106 | SRR11238036 | 1842158 | DGC | 73.95 | 189.68 | 935008(50.8%) |
| DGCm084 | SRR11238036 | 2989975 | DGC | 54.63 | 180.93 | 1274767(42.6%) |
| DGCm085 | SRR11238036 | 3159714 | DGC | 52.7 | 182.13 | 1350353(42.7%) |
| DGCm086 | SRR11238036 | 3187641 | DGC | 46.93 | 166.77 | 1359170(42.6%) |
| DGCm094 | SRR11238036 | 2713601 | DGC | 46.67 | 161.8 | 1217329(44.9%) |
| DGCm095 | SRR11238036 | 1943761 | DGC | 70.31 | 187.58 | 1031539(53.1%) |
| DGCm096 | SRR11238036 | 2103517 | DGC | 58.69 | 171.15 | 982742(46.7%) |
| DGCm097 | SRR11238036 | 2862619 | DGC | 55.08 | 182.06 | 1272021(44.4%) |
| DGCm098 | SRR11238036 | 2721654 | DGC | 65.46 | 197.52 | 1196617(44.0%) |
| DGCm099 | SRR11238036 | 2839000 | DGC | 52.21 | 175.53 | 1220008(43.0%) |
| DGCm100 | SRR11238036 | 7576466 | DGC | 37.61 | 142.97 | 1675952(22.1%) |
| DGCm101 | SRR11238036 | 2847149 | DGC | 63.24 | 198.45 | 1292574(45.4%) |
| DGCm102 | SRR11238036 | 3267814 | DGC | 51.3 | 179.45 | 1398105(42.8%) |
| ETBm188 | SRR11238035 | 3010657 | ETB | 33.87 | 105.18 | 1534229(51.0%) |
| ETBf203 | SRR11238035 | 2180678 | ETB | 36.2 | 113.04 | 1014987(46.5%) |
| ETBm189 | SRR11238035 | 3911583 | ETB | 37.79 | 122.08 | 2002976(51.2%) |
| ETBm190 | SRR11238035 | 3447634 | ETB | 34.86 | 114.97 | 1767783(51.3%) |
| ETBm191 | SRR11238035 | 2339047 | ETB | 32.52 | 102.82 | 1252671(53.6%) |
| ETBm192 | SRR11238035 | 1294040 | ETB | 27.19 | 77.33 | 678519(52.4%) |
| ETBm193 | SRR11238035 | 1289852 | ETB | 30.53 | 87.21 | 679367(52.7%) |
| ETBm194 | SRR11238035 | 1472576 | ETB | 31.97 | 92.99 | 795190(54.0%) |
| ETBm199 | SRR11238035 | 2165577 | ETB | 33.65 | 105.4 | 1106314(51.1%) |
| ETBm200 | SRR11238035 | 2661036 | ETB | 35.11 | 112.4 | 1370634(51.5%) |
| ETBm201 | SRR11238035 | 2335170 | ETB | 31.7 | 99.09 | 1166820(50.0%) |
| ETBm202 | SRR11238035 | 3364649 | ETB | 35.45 | 113.06 | 1723046(51.2%) |
| ETBm204 | SRR11238035 | 2017944 | ETB | 38.1 | 107.19 | 998453(49.5%) |
| ETBm205 | SRR11238035 | 2054444 | ETB | 34.09 | 105.32 | 1047208(51.0%) |
| FRNfM02 | SRR11238035 | 4128301 | FRN | 33.69 | 105.83 | 2134077(51.7%) |

Table S3: Summary of RAD Loci assembly using Stacks *(continued)*

| Sample | SRA Accession | Total Reads | POP | Average Coverage of Stacks | Standard Deviation | No. of Reads Used |
|---|---|---|---|---|---|---|
| FRNfM04 | SRR11238035 | 3691899 | FRN | 31.37 | 96.31 | 1755740(47.6%) |
| FRNmM03 | SRR11238035 | 2576724 | FRN | 31.86 | 97.21 | 1293688(50.2%) |
| FRNmM05 | SRR11238035 | 3167685 | FRN | 36.48 | 111.38 | 1674427(52.9%) |
| FRNmM06 | SRR11238035 | 3345278 | FRN | 33.2 | 100.24 | 1727404(51.6%) |
| FRNmM07 | SRR11238035 | 2992795 | FRN | 32.87 | 98.82 | 1537909(51.4%) |
| MDCf134 | SRR11238036 | 1531260 | MDC | 73.68 | 173.21 | 746189(48.7%) |
| MDCf135 | SRR11238036 | 3193961 | MDC | 55 | 187.99 | 1402656(43.9%) |
| MDCf136 | SRR11238036 | 3182956 | MDC | 55.02 | 189.9 | 1311574(41.2%) |
| MDCf138 | SRR11238036 | 3392174 | MDC | 55.54 | 190.24 | 1392772(41.1%) |
| MDCf139 | SRR11238036 | 2894914 | MDC | 60.18 | 190.11 | 1285166(44.4%) |
| MDCf142 | SRR11238036 | 2816815 | MDC | 55.25 | 184.98 | 1217701(43.2%) |
| MDCf144 | SRR11238036 | 3491288 | MDC | 53.85 | 188.32 | 1428296(40.9%) |
| MDCf146 | SRR11238036 | 2823133 | MDC | 64.6 | 197.79 | 1155219(40.9%) |
| MDCm140 | SRR11238036 | 1750868 | MDC | 78.76 | 193.98 | 876127(50.0%) |
| MDCm141 | SRR11238036 | 3675988 | MDC | 50.44 | 182.7 | 1434632(39.0%) |
| MDCm143 | SRR11238036 | 3264423 | MDC | 49.5 | 174.36 | 1355402(41.5%) |
| MDCm145 | SRR11238036 | 3276452 | MDC | 53.34 | 184.17 | 1305359(39.8%) |
| MLGf003 | SRR11238036 | 3331468 | MLG | 44.93 | 163.27 | 1330734(39.9%) |
| MLGf005 | SRR11238036 | 3269479 | MLG | 53.71 | 182.14 | 1322214(40.4%) |
| MLGf008 | SRR11238036 | 2944072 | MLG | 53.13 | 181.42 | 1373732(46.7%) |
| MLGf010 | SRR11238036 | 3277359 | MLG | 57.57 | 190.14 | 1343152(41.0%) |
| MLGf011 | SRR11238036 | 3511746 | MLG | 53.3 | 183.08 | 1387321(39.5%) |
| MLGm001 | SRR11238036 | 3364196 | MLG | 47.52 | 169.04 | 1451538(43.1%) |
| MLGm002 | SRR11238036 | 2901880 | MLG | 49.48 | 167.43 | 1273448(43.9%) |
| MLGm004 | SRR11238036 | 2581961 | MLG | 48.66 | 164.76 | 1110997(43.0%) |
| MLGm006 | SRR11238036 | 4754478 | MLG | 41.36 | 157.36 | 1444616(30.4%) |
| MLGm007 | SRR11238036 | 3078368 | MLG | 54.91 | 183.7 | 1372169(44.6%) |
| MLGm009 | SRR11238036 | 3157171 | MLG | 50.16 | 173.76 | 1344150(42.6%) |
| MLGm012 | SRR11238036 | 3134956 | MLG | 57.55 | 192.04 | 1296623(41.4%) |
| MLGm013 | SRR11238036 | 3484557 | MLG | 52.91 | 183.06 | 1363605(39.1%) |
| MLGm014 | SRR11238036 | 2858870 | MLG | 55.4 | 184.75 | 1194456(41.8%) |
| MLGm016 | SRR11238036 | 2214552 | MLG | 71.4 | 200.21 | 1068911(48.3%) |
| MLGm017 | SRR11238036 | 3695528 | MLG | 55.43 | 192.61 | 1524083(41.2%) |
| MMSf206 | SRR11238035 | 3707087 | MMS | 33.52 | 104.64 | 1885075(50.9%) |
| MMSf211 | SRR11238035 | 2202122 | MMS | 29.2 | 89.98 | 1081684(49.1%) |
| MMSf215 | SRR11238035 | 3840292 | MMS | 36.92 | 113.55 | 1933486(50.3%) |
| MMSf217 | SRR11238035 | 2707625 | MMS | 30.72 | 92.05 | 1344134(49.6%) |
| MMSf221 | SRR11238035 | 4208537 | MMS | 35.72 | 104.23 | 2149741(51.1%) |
| MMSf222 | SRR11238035 | 3952532 | MMS | 32.63 | 96.17 | 1968865(49.8%) |
| MMSm207 | SRR11238035 | 3517181 | MMS | 34.73 | 109.29 | 1836941(52.2%) |

Table S3: Summary of RAD Loci assembly using Stacks *(continued)*

| **Sample** | **SRA Accession** | **Total Reads** | **POP** | **Average Coverage of Stacks** | **Standard Deviation** | **No. of Reads Used** |
|---|---|---|---|---|---|---|
| MMSm208 | SRR11238035 | 6585205 | MMS | 39.8 | 105.73 | 3123181(47.4%) |
| MMSm209 | SRR11238035 | 3346988 | MMS | 33.87 | 102.05 | 1753505(52.4%) |
| MMSm210 | SRR11238035 | 3233531 | MMS | 34.83 | 103.98 | 1678764(51.9%) |
| MMSm212 | SRR11238035 | 3408275 | MMS | 34.06 | 109.98 | 1801319(52.9%) |
| MMSm213 | SRR11238035 | 2318554 | MMS | 31.13 | 94.72 | 1241729(53.6%) |
| MMSm214 | SRR11238035 | 3178148 | MMS | 30.59 | 87.25 | 1651822(52.0%) |
| MMSm220 | SRR11238035 | 4093188 | MMS | 35.24 | 108.45 | 2039613(49.8%) |
| OBNf111 | SRR11238036 | 2630822 | OBN | 64.41 | 196.32 | 1209675(46.0%) |
| OBNf121 | SRR11238036 | 2690338 | OBN | 60.94 | 185.86 | 1177813(43.8%) |
| OBNm110 | SRR11238036 | 2681431 | OBN | 47.36 | 159.57 | 1284679(47.9%) |
| OBNm112 | SRR11238036 | 2751978 | OBN | 63.41 | 197.95 | 1272679(46.2%) |
| OBNm113 | SRR11238036 | 3194428 | OBN | 64.07 | 207.61 | 1391156(43.5%) |
| OBNm114 | SRR11238036 | 227420 | OBN | n/a | n/a | n/a |
| OBNm115 | SRR11238036 | 2830379 | OBN | 61.76 | 197.96 | 1348164(47.6%) |
| OBNm116 | SRR11238036 | 3026011 | OBN | 44.43 | 160.46 | 1329775(43.9%) |
| OBNm117 | SRR11238036 | 2249328 | OBN | 60 | 180.73 | 1145652(50.9%) |
| OBNm118 | SRR11238036 | 1753843 | OBN | 32.88 | 112.75 | 858594(49.0%) |
| OBNm119 | SRR11238036 | 2786199 | OBN | 51.81 | 171.04 | 1253529(45.0%) |
| OBNm120 | SRR11238036 | 3188535 | OBN | 63.35 | 203.53 | 1375248(43.1%) |
| OBNm122 | SRR11238036 | 1963010 | OBN | 63.4 | 179.44 | 969701(49.4%) |
| OBNm123 | SRR11238036 | 1680297 | OBN | 63.39 | 171.31 | 833328(49.6%) |
| OBNm124 | SRR11238036 | 754559 | OBN | 69.41 | 137.52 | 393118(52.1%) |
| PCPm161 | SRR11238035 | 1798002 | PCP | 34.68 | 94.93 | 897699(49.9%) |
| PCPf175 | SRR11238035 | 3534243 | PCP | 32.62 | 98.96 | 1772238(50.1%) |
| PCPm156 | SRR11238035 | 3363460 | PCP | 32.67 | 106.51 | 1705822(50.7%) |
| PCPm157 | SRR11238035 | 2928578 | PCP | 31.7 | 101.07 | 1523220(52.0%) |
| PCPm158 | SRR11238035 | 3130749 | PCP | 30.38 | 96.42 | 1614384(51.6%) |
| PCPm159 | SRR11238035 | 3436320 | PCP | 34.03 | 114.25 | 1757627(51.1%) |
| PCPm160 | SRR11238035 | 2575990 | PCP | 33.15 | 111.19 | 1289258(50.0%) |
| PCPm162 | SRR11238035 | 1826962 | PCP | 31.34 | 93.84 | 824280(45.1%) |
| PCPm172 | SRR11238035 | 2959900 | PCP | 37.35 | 110.05 | 1551248(52.4%) |
| PCPm173 | SRR11238035 | 1373989 | PCP | 28.97 | 75.26 | 696769(50.7%) |
| PCPm174 | SRR11238035 | 781039 | PCP | 35.39 | 76.64 | 390583(50.0%) |
| PCPm176 | SRR11238035 | 1701200 | PCP | 29.14 | 87.69 | 877375(51.6%) |
| PCPm177 | SRR11238035 | 2015460 | PCP | 28.74 | 80.42 | 1035047(51.4%) |
| RHDf242 | SRR11238035 | 3187266 | RHD | 35.64 | 111.25 | 1396716(43.8%) |
| RHDf245 | SRR11238035 | 3282691 | RHD | 32.61 | 103.53 | 1403744(42.8%) |
| RHDf253 | SRR11238035 | 1198067 | RHD | 54.19 | 130.59 | 528822(44.1%) |
| RHDm239 | SRR11238035 | 3816448 | RHD | 31.9 | 100.38 | 1870860(49.0%) |
| RHDm240 | SRR11238035 | 3883290 | RHD | 34.65 | 115.21 | 1893018(48.7%) |

Table S3: Summary of RAD Loci assembly using Stacks *(continued)*

| **Sample** | **SRA Accession** | **Total Reads** | **POP** | **Average Coverage of Stacks** | **Standard Deviation** | **No. of Reads Used** |
|---|---|---:|---|---|---|---|
| RHDm241 | SRR11238035 | 1863382 | RHD | 31.93 | 92.75 | 944340(50.7%) |
| RHDm246 | SRR11238035 | 1379829 | RHD | 29.96 | 82.25 | 664141(48.1%) |
| RHDm247 | SRR11238035 | 4254163 | RHD | 33.1 | 102.82 | 2017608(47.4%) |
| RHDm248 | SRR11238035 | 3382304 | RHD | 34.67 | 110.64 | 1778628(52.6%) |
| RHDm254 | SRR11238035 | 2413011 | RHD | 33.44 | 99.08 | 1220013(50.6%) |
| RHDm255 | SRR11238035 | 2782584 | RHD | 33.81 | 105.43 | 1372736(49.3%) |
| RHDm256 | SRR11238035 | 2032467 | RHD | 33.49 | 99.93 | 997506(49.1%) |
| RHDm257 | SRR11238035 | 4126879 | RHD | 34.04 | 111.19 | 2019138(48.9%) |
| RVSfR3 | SRR11238036 | 3181026 | RVS | 63.88 | 200.8 | 1364721(42.9%) |
| RVSfR4 | SRR11238036 | 2034084 | RVS | 70.88 | 191.46 | 977547(48.1%) |
| RVSmR1 | SRR11238036 | 3068970 | RVS | 53.43 | 178.86 | 1340289(43.7%) |
| RVSmR2 | SRR11238036 | 2084868 | RVS | 65.18 | 186.73 | 1036699(49.7%) |
| RVSmR5 | SRR11238036 | 2532622 | RVS | 65.86 | 200.21 | 1181758(46.7%) |
| RVSmR6 | SRR11238036 | 2743057 | RVS | 61.31 | 190.52 | 1279604(46.6%) |
| TULf054 | SRR11238036 | 3159299 | TUL | 63.25 | 203.62 | 1340220(42.4%) |
| TULf061 | SRR11238036 | 2598097 | TUL | 63.25 | 197.59 | 1136439(43.7%) |
| TULf064 | SRR11238036 | 3009782 | TUL | 77.02 | 199.59 | 380263(12.6%) |
| TULf072 | SRR11238036 | 3142924 | TUL | 55.04 | 187.55 | 1387313(44.1%) |
| TULf073 | SRR11238036 | 3443965 | TUL | 51.62 | 178.8 | 1416133(41.1%) |
| TULf075 | SRR11238036 | 3457076 | TUL | 54.03 | 188.55 | 1427764(41.3%) |
| TULf077 | SRR11238036 | 3473548 | TUL | 51.94 | 183.85 | 1380019(39.7%) |
| TULf078 | SRR11238036 | 3054081 | TUL | 53.43 | 183.06 | 1280522(41.9%) |
| TULm053 | SRR11238036 | 1575217 | TUL | 95.78 | 208.01 | 819583(52.0%) |
| TULm055 | SRR11238036 | 1455895 | TUL | 113.89 | 237.36 | 722208(49.6%) |
| TULm060 | SRR11238036 | 3382266 | TUL | 53.34 | 184.86 | 1299380(38.4%) |
| TULm062 | SRR11238036 | 3109015 | TUL | 60.93 | 195.83 | 1265235(40.7%) |
| TULm063 | SRR11238036 | 3525244 | TUL | 47.52 | 173.53 | 1442273(40.9%) |
| TULm069 | SRR11238036 | 2774500 | TUL | 55.47 | 182.19 | 1162454(41.9%) |
| TULm070 | SRR11238036 | 2561524 | TUL | 69.26 | 203.09 | 1125614(43.9%) |
| TULm071 | SRR11238036 | 4247179 | TUL | 42.8 | 163.37 | 1444012(34.0%) |

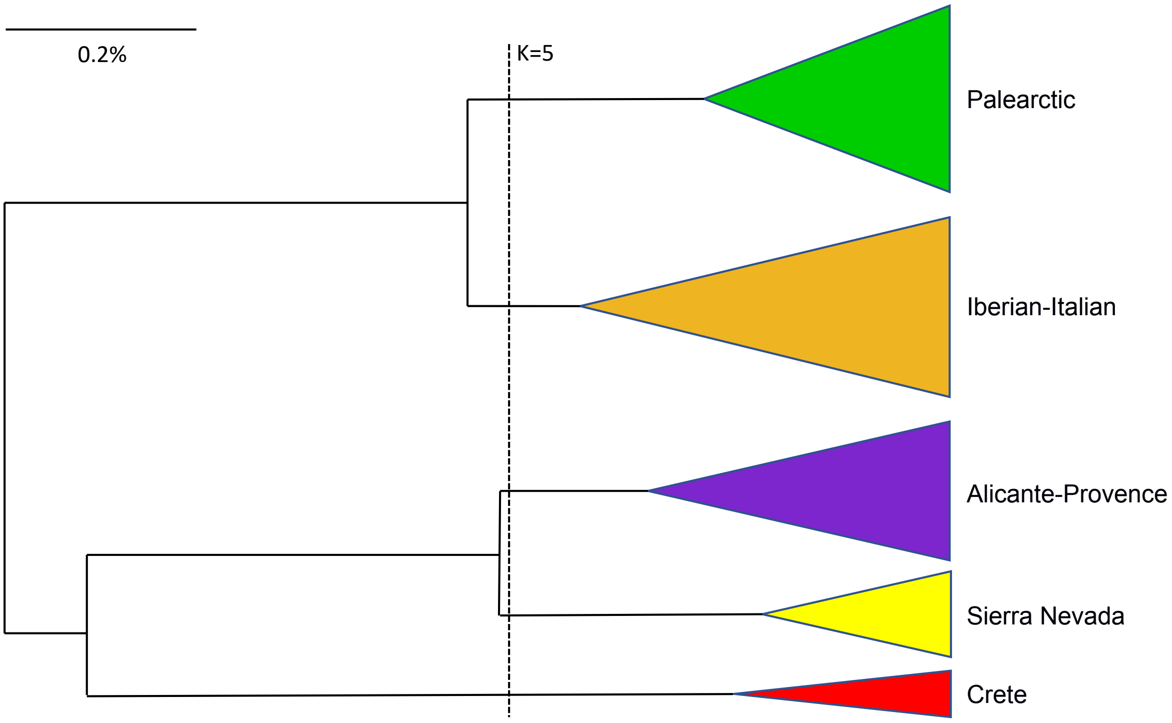


**Figure S4** UPGMA clustering of all 587 *Polyommatus icarus* mitochondrial *CO1* sequences (Table S2). The horizontal red bar cuts at k=5, and these 5 clusters corresponds exactly to the *CO1* lineage classification based in Dincă *et al.* (2011). The size of the collapsed tips is proportional to the log of the sample size in those lineages.

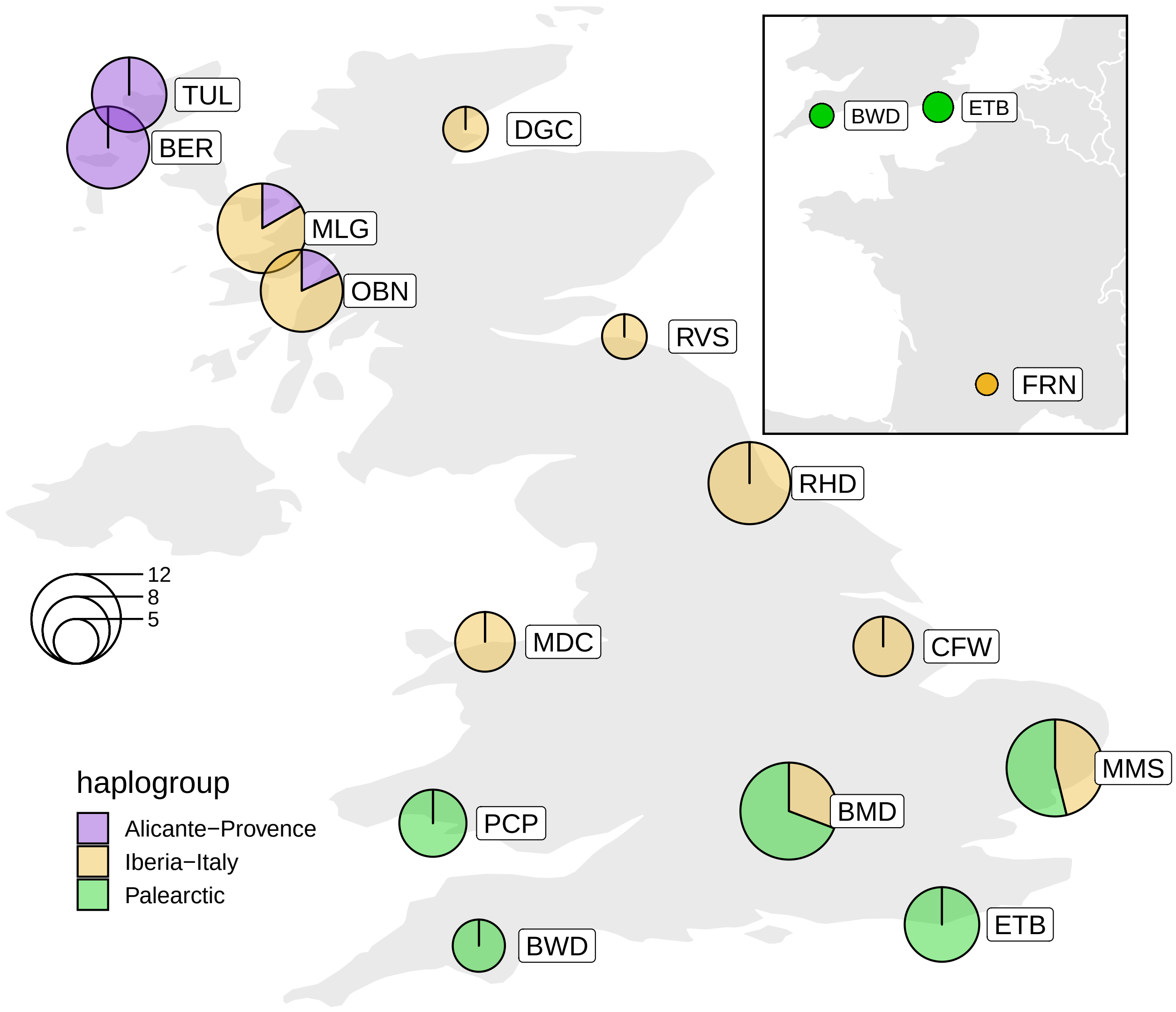


**Figure S5** *CO1* Haplotype composition in the British Isles. Lineages or haplogroups are classified are based on Dincă *et al.* (2011). Circles are proportional to sample size.

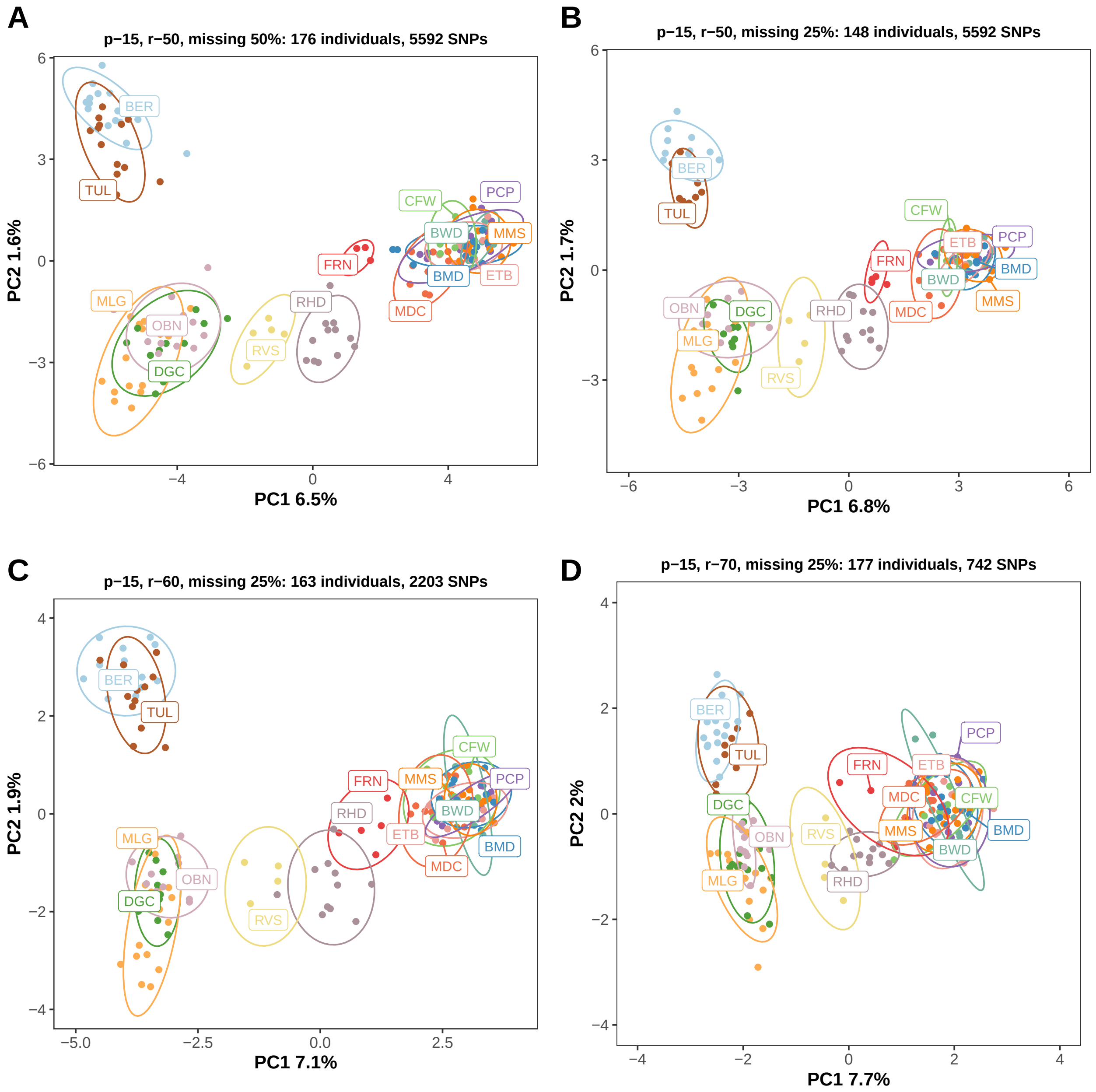


**Figure S6** Population structure based on principal component analysis (PCA) of ddRADseq SNP datasets with varying levels of missing data and number of markers. **(A)** This SNP marker dataset is the same as Figure 3 but only individuals with > 50% missing data were removed. **(B)** This is a dataset with the same exclusion criteria for missing data (>25%) as in Figure 3 but with only unlinked SNPs (1st SNP on each RAD locus). (C) and (D) are PCAs based on a datasets with more stringent criteria for including loci (present in at least 60% or 70% of individuals across all localilites), and hence lesser number of loci. Ninety-% Confidence Interval (CI) ellipses for PC1 and PC2 for each locality are also shown.

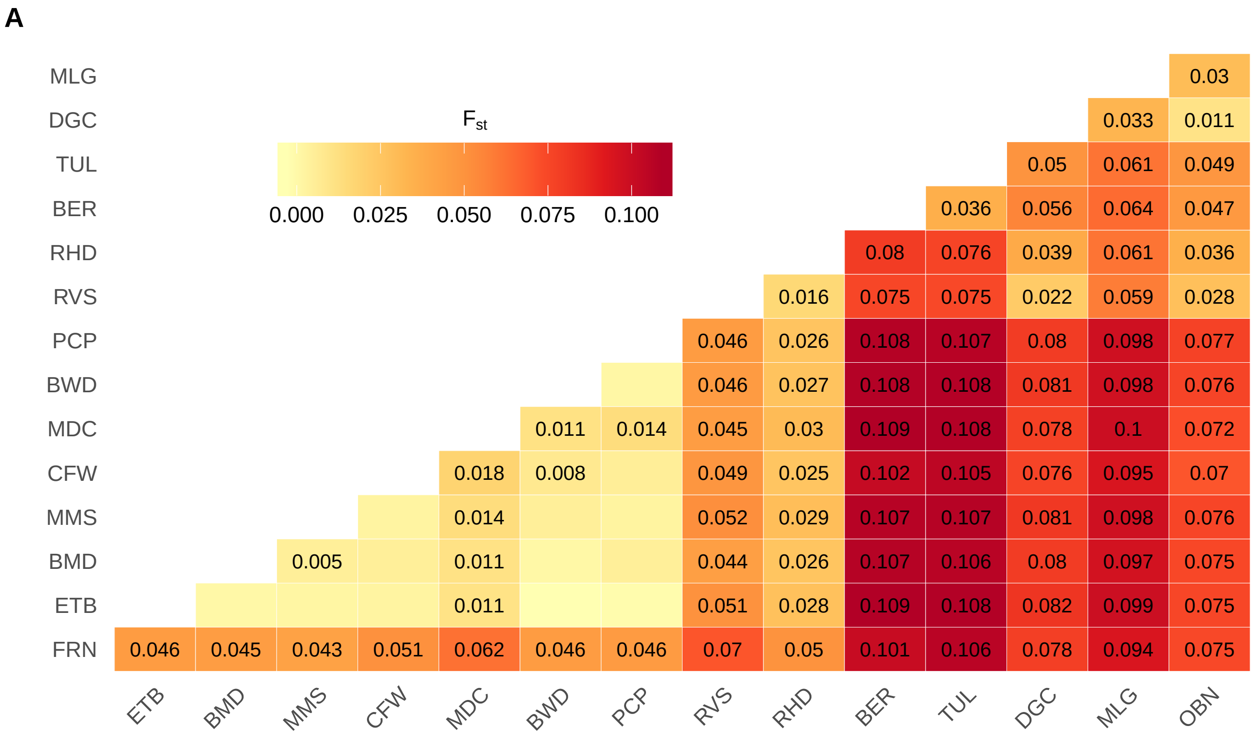


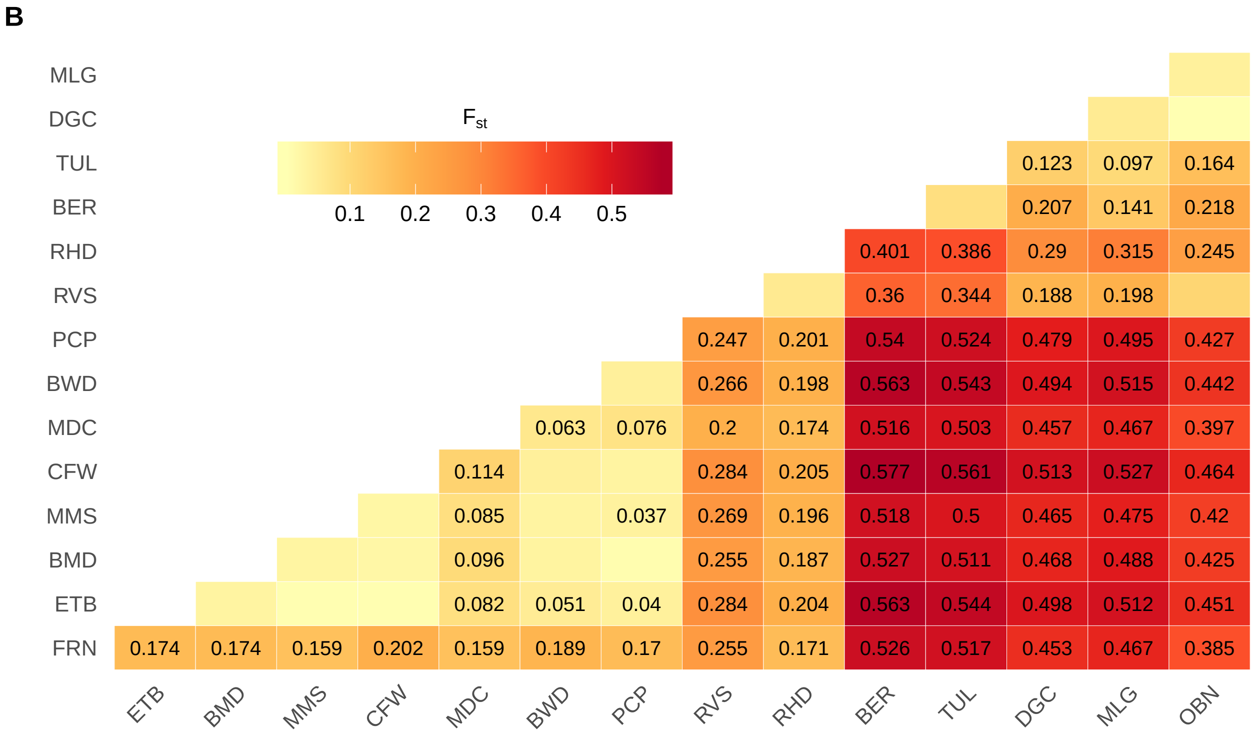


**Figure S7** Weir and Cockerham estimates of population pariwise $F_{st}$ across all 15 populations using **(A)** 5387 putatively neutral SNPS and **(B)** 104 outlier SNPs. Pariwise estimates that were statistically different from zero (based on 10000 permuations) are displayed. Populations are grouped according to geographical distance.

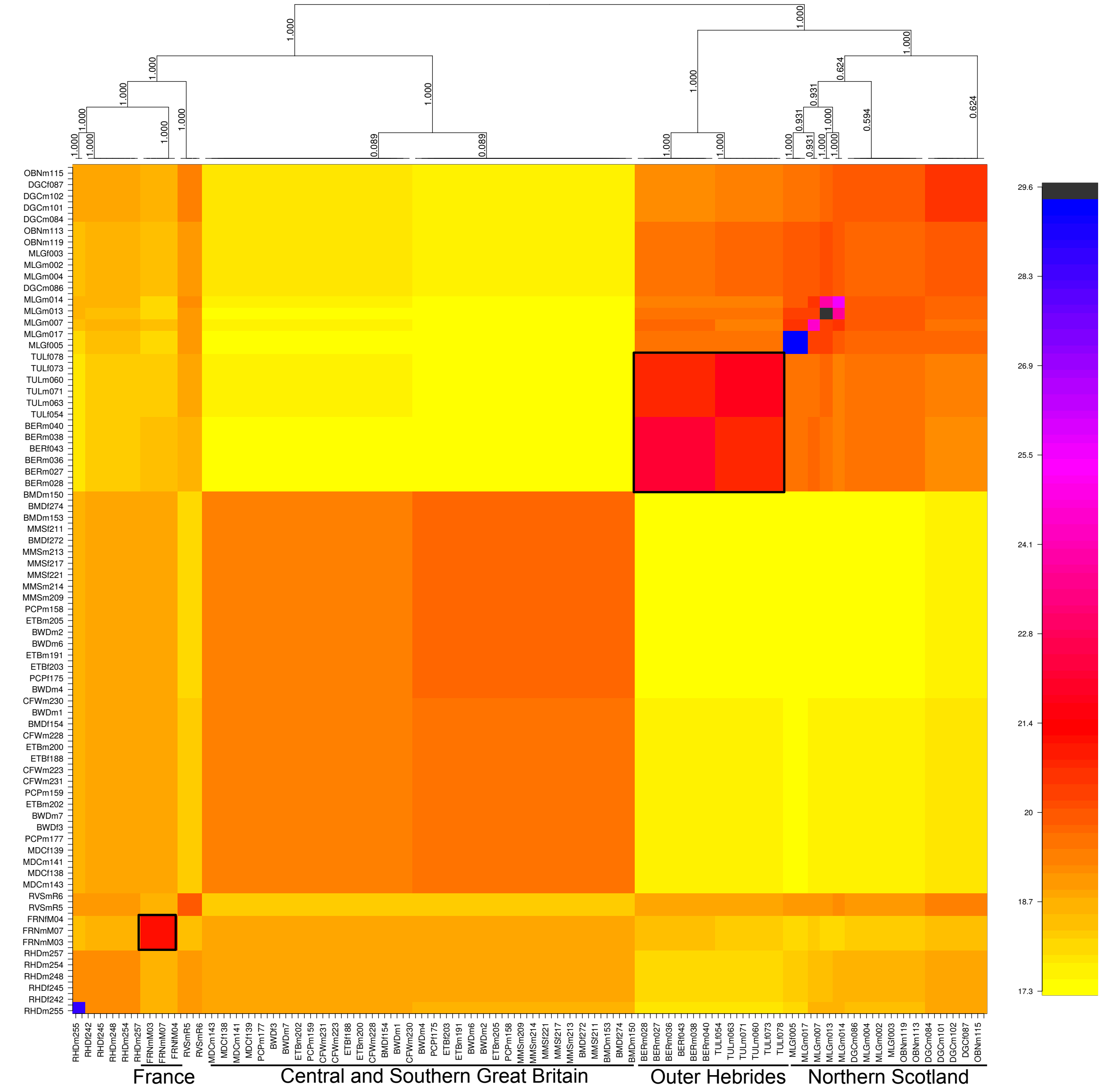


**Figure S8** Clustered fineRADstructure population-averaged co-ancestry matrix based on 148 individuals and 5387 putatively neutral SNPs. Individuals from Northern Scottish, Southern English and Welsh, and French locations cluster together. The Outer Hebrides (large black outline) show further structuring within the Northern Scottish cluster. Individuals from southern Scotland (RVS) and northern England (RHD) show varying levels of coancestry with northern and southern populations, reperesenting hybrid populations in case of RVS and RHD. FRN (small black outline) also shares coancestry with northern and southern populations but represents a distinct cluster of individuals. Some individuals from MLG share high ancestry and may be close relatives. Note: note all individuals are labelled.

Table S4: Summary of Reads mapping toWolbachia

| **Sample** | **Total Reads** | **Population** | **% Total Reads Classified by Centrifuge** | **% Total Reads Classified as Microbial (Archeae and Bacteria)** | **% Total Reads Classified as Bacterial** | **% of Classified Reads mapping to Wolbachia wNo** | **% of Classified Reads mapping to other Wolbachia taxa** |
|---|---|---|---|---|---|---|---|
| BERf030 | 3129338 | BER | 5.11 | 5.11 | 5.10 | 43.1900000 | 0.5481900 |
| BERf035 | 3317737 | BER | 5.73 | 5.73 | 5.73 | 49.1400000 | 0.4403900 |
| BERf037 | 3695537 | BER | 4.61 | 4.61 | 4.59 | 26.1900000 | 0.3305567 |
| BERf043 | 3071772 | BER | 5.19 | 5.19 | 5.18 | 42.6200000 | 0.2948000 |
| BERm027 | 3195023 | BER | 3.78 | 3.78 | 3.77 | 12.8000000 | 0.2317670 |
| BERm028 | 6952171 | BER | 3.13 | 3.13 | 3.12 | 0.0106300 | 0.0000000 |
| BERm029 | 8470130 | BER | 2.67 | 2.67 | 2.66 | 0.0055870 | 0.0004656 |
| BERm036 | 3855171 | BER | 3.43 | 3.43 | 3.40 | 11.6400000 | 0.2016500 |
| BERm038 | 2859523 | BER | 3.40 | 3.40 | 3.38 | 0.0733600 | 0.0010790 |
| BERm039 | 2149488 | BER | 5.26 | 5.26 | 5.25 | 14.4900000 | 0.2747400 |
| BERm040 | 3457223 | BER | 3.32 | 3.32 | 3.31 | 12.3900000 | 0.2036490 |
| BERm041 | 1436794 | BER | 5.01 | 5.01 | 4.99 | 0.1270000 | 0.0000000 |
| BERm042 | 3264460 | BER | 3.56 | 3.56 | 3.55 | 20.0500000 | 0.3241100 |
| BERm044 | 3122980 | BER | 3.96 | 3.96 | 3.95 | 20.4600000 | 0.2758500 |
| BERm045 | 3168433 | BER | 3.61 | 3.61 | 3.60 | 12.7500000 | 0.2543100 |
| BERm046 | 3620747 | BER | 2.84 | 2.84 | 2.83 | 0.0159400 | 0.0000000 |
| BMDf149 | 3713282 | BMD | 3.43 | 3.43 | 3.42 | 0.0187000 | 0.0000000 |
| BMDf154 | 3984391 | BMD | 3.27 | 3.27 | 3.26 | 0.0638600 | 0.0000000 |
| BMDf270 | 2961330 | BMD | 5.52 | 5.52 | 5.49 | 0.0233700 | 0.0000000 |
| BMDf272 | 3200952 | BMD | 5.51 | 5.51 | 5.48 | 0.0435900 | 0.0000000 |
| BMDf274 | 2556011 | BMD | 7.99 | 7.99 | 7.92 | 0.0615200 | 0.0000000 |
| BMDf276 | 2740520 | BMD | 7.65 | 7.65 | 7.59 | 0.0482800 | 0.0000000 |
| BMDf278 | 1893829 | BMD | 7.78 | 7.78 | 7.76 | 0.0559900 | 0.0000000 |
| BMDf279 | 3436779 | BMD | 5.17 | 5.17 | 5.14 | 0.0237200 | 0.0000000 |
| BMDm147 | 3537215 | BMD | 3.64 | 3.64 | 3.63 | 0.0097380 | 0.0000000 |
| BMDm148 | 3441037 | BMD | 4.06 | 4.06 | 4.05 | 0.0074120 | 0.0000000 |
| BMDm150 | 3947571 | BMD | 4.98 | 4.98 | 4.94 | 0.0026990 | 0.0000000 |
| BMDm151 | 3500869 | BMD | 4.15 | 4.15 | 4.13 | NA | 0.0007204 |
| BMDm152 | 3707329 | BMD | 4.69 | 4.69 | 4.68 | 0.0054030 | 0.0000000 |
| BMDm153 | 3620544 | BMD | 4.41 | 4.41 | 4.39 | 0.0077000 | 0.0000000 |
| BMDm271 | 1723401 | BMD | 8.17 | 8.17 | 8.15 | 0.0022490 | 0.0000000 |
| BMDm275 | 1390969 | BMD | 8.70 | 8.70 | 8.68 | 0.0069260 | 0.0000000 |
| BWDf3 | 4621287 | BWD | 3.75 | 3.75 | 3.71 | 0.0406700 | 0.0000000 |
| BWDf5 | 4042362 | BWD | 4.75 | 4.75 | 4.74 | 0.0108500 | 0.0000000 |
| BWDm1 | 4163695 | BWD | 3.84 | 3.84 | 3.82 | 0.0065330 | 0.0000000 |
| BWDm2 | 4653162 | BWD | 4.55 | 4.55 | 4.52 | 0.0033600 | 0.0000000 |

Table S4: Summary of Reads mapping toWolbachia *(continued)*

| **Sample** | **Total Reads** | **Population** | **% Total Reads Classified by Centrifuge** | **% Total Reads Classified as Microbial (Archeae and Bacteria)** | **% Total Reads Classified as Bacterial** | **% of Classified Reads mapping to Wolbachia wNo** | **% of Classified Reads mapping to other Wolbachia taxa** |
|---|---|---|---|---|---|---|---|
| BWDm4 | 4663845 | BWD | 9.33 | 9.33 | 9.30 | 0.0055160 | 0.0000000 |
| BWDm6 | 4508000 | BWD | 3.88 | 3.88 | 3.87 | 0.0067830 | 0.0000000 |
| BWDm7 | 4181443 | BWD | 7.75 | 7.75 | 7.71 | 0.0003616 | 0.0003616 |
| DGCf087 | 3347527 | DGC | 4.00 | 4.00 | 3.99 | 6.5330000 | 0.1235270 |
| DGCf106 | 1842158 | DGC | 4.74 | 4.74 | 4.73 | 4.9620000 | 0.0853450 |
| DGCm084 | 2989975 | DGC | 3.50 | 3.50 | 3.50 | 7.9210000 | 0.1539700 |
| DGCm085 | 3159714 | DGC | 3.79 | 3.79 | 3.78 | 11.7200000 | 0.1708470 |
| DGCm086 | 3187641 | DGC | 3.76 | 3.76 | 3.75 | 8.4960000 | 0.1615860 |
| DGCm094 | 2713601 | DGC | 4.09 | 4.09 | 4.08 | 8.3930000 | 0.1404810 |
| DGCm095 | 1943761 | DGC | 5.29 | 5.29 | 5.28 | 0.0183700 | 0.0000000 |
| DGCm096 | 2103517 | DGC | 4.45 | 4.45 | 4.43 | 0.0753400 | 0.0022500 |
| DGCm097 | 2862619 | DGC | 3.43 | 3.43 | 3.42 | 10.3400000 | 0.1570170 |
| DGCm098 | 2721654 | DGC | 3.69 | 3.69 | 3.68 | 0.0571900 | 0.0000000 |
| DGCm099 | 2839000 | DGC | 3.34 | 3.34 | 3.33 | 0.0369800 | 0.0000000 |
| DGCm100 | 7576466 | DGC | 2.98 | 2.98 | 2.98 | 7.2910000 | 0.1005850 |
| DGCm101 | 2847149 | DGC | 3.40 | 3.40 | 3.39 | 6.5730000 | 0.1118720 |
| DGCm102 | 3267814 | DGC | 4.07 | 4.07 | 4.06 | 6.8670000 | 0.1086340 |
| ETBf188 | 3010657 | ETB | 4.98 | 4.98 | 4.94 | 0.0056900 | 0.0000000 |
| ETBf203 | 2180678 | ETB | 5.88 | 5.88 | 5.87 | 0.0308700 | 0.0000000 |
| ETBm189 | 3911583 | ETB | 3.48 | 3.48 | 3.46 | NA | 0.0000000 |
| ETBm190 | 3447634 | ETB | 4.85 | 4.85 | 4.80 | 0.0031670 | 0.0000000 |
| ETBm191 | 2339047 | ETB | 5.90 | 5.90 | 5.86 | 0.0193800 | 0.0000000 |
| ETBm192 | 1294040 | ETB | 7.82 | 7.82 | 7.79 | 0.0112100 | 0.0010190 |
| ETBm193 | 1289852 | ETB | 7.41 | 7.41 | 7.39 | NA | 0.0000000 |
| ETBm194 | 1472576 | ETB | 7.02 | 7.02 | 7.00 | 0.0030650 | 0.0010220 |
| ETBm199 | 2165577 | ETB | 5.72 | 5.72 | 5.70 | 0.0042370 | 0.0000000 |
| ETBm200 | 2661036 | ETB | 5.07 | 5.07 | 5.04 | 0.0061210 | 0.0000000 |
| ETBm201 | 2335170 | ETB | 6.35 | 6.35 | 6.30 | 0.0121200 | 0.0000000 |
| ETBm202 | 3364649 | ETB | 4.83 | 4.83 | 4.80 | 0.0083660 | 0.0000000 |
| ETBm204 | 2017944 | ETB | 6.55 | 6.55 | 6.53 | 0.0054950 | 0.0000000 |
| ETBm205 | 2054444 | ETB | 6.23 | 6.23 | 6.21 | 0.0008167 | 0.0000000 |
| FRNfM02 | 4128301 | FRN | 4.92 | 4.92 | 4.86 | 0.0226500 | 0.0000000 |
| FRNfM04 | 3691899 | FRN | 4.88 | 4.88 | 4.84 | 0.0713100 | 0.0017680 |
| FRNmM03 | 2576724 | FRN | 6.69 | 6.69 | 6.65 | 0.0085340 | 0.0000000 |
| FRNmM05 | 3167685 | FRN | 4.72 | 4.72 | 4.69 | 0.0048750 | 0.0000000 |
| FRNmM06 | 3345278 | FRN | 5.45 | 5.45 | 5.41 | 0.0017350 | 0.0000000 |

Table S4: Summary of Reads mapping toWolbachia *(continued)*

| **Sample** | **Total Reads** | **Population** | **% Total Reads Classified by Centrifuge** | **% Total Reads Classified as Microbial (Archeae and Bacteria)** | **% Total Reads Classified as Bacterial** | **% of Classified Reads mapping to Wolbachia wNo** | **% of Classified Reads mapping to other Wolbachia taxa** |
|---|---:|---|---:|---:|---:|---:|---:|
| FRNmM07 | 2992795 | FRN | 5.82 | 5.82 | 5.79 | 0.0030250 | 0.0000000 |
| MDCf134 | 1531260 | MDC | 4.18 | 4.18 | 4.16 | 0.1970000 | 0.0016020 |
| MDCf135 | 3193961 | MDC | 2.51 | 2.51 | 2.50 | 0.1118000 | 0.0000000 |
| MDCf136 | 3182956 | MDC | 2.34 | 2.34 | 2.33 | 0.2309000 | 0.0014430 |
| MDCf138 | 3392174 | MDC | 2.52 | 2.52 | 2.52 | 0.1846000 | 0.0012230 |
| MDCf139 | 2894914 | MDC | 3.08 | 3.08 | 3.08 | 0.1566000 | 0.0000000 |
| MDCf142 | 2816815 | MDC | 2.85 | 2.85 | 2.84 | 0.1359000 | 0.0027180 |
| MDCf144 | 3491288 | MDC | 2.41 | 2.41 | 2.40 | 0.1269000 | 0.0024170 |
| MDCf146 | 2823133 | MDC | 2.81 | 2.81 | 2.80 | 0.2156000 | 0.0065710 |
| MDCm140 | 1750868 | MDC | 4.76 | 4.76 | 4.75 | 0.0051320 | 0.0000000 |
| MDCm141 | 3675988 | MDC | 2.54 | 2.54 | 2.53 | 0.0652600 | 0.0022900 |
| MDCm143 | 3264423 | MDC | 2.91 | 2.91 | 2.90 | 0.1208000 | 0.0011080 |
| MDCm145 | 3276452 | MDC | 2.77 | 2.77 | 2.76 | NA | 0.0000000 |
| MLGf003 | 3331468 | MLG | 3.36 | 3.36 | 3.35 | 0.2643000 | 0.0009576 |
| MLGf005 | 3269479 | MLG | 3.63 | 3.63 | 3.60 | 5.9680000 | 0.0919720 |
| MLGf008 | 2944072 | MLG | 3.97 | 3.97 | 3.96 | 4.6690000 | 0.1006500 |
| MLGf010 | 3277359 | MLG | 3.35 | 3.35 | 3.34 | 6.8430000 | 0.3241526 |
| MLGf011 | 3511746 | MLG | 3.15 | 3.15 | 3.15 | 6.7270000 | 0.0875231 |
| MLGm001 | 3364196 | MLG | 3.46 | 3.46 | 3.45 | 0.0938200 | 0.0000000 |
| MLGm002 | 2901880 | MLG | 4.23 | 4.23 | 4.22 | 10.8800000 | 0.2047000 |
| MLGm004 | 2581961 | MLG | 3.06 | 3.06 | 3.05 | 0.0723600 | 0.0000000 |
| MLGm006 | 4754478 | MLG | 4.31 | 4.31 | 4.30 | 4.3210000 | 0.0948030 |
| MLGm007 | 3078368 | MLG | 3.46 | 3.46 | 3.45 | 7.9880000 | 0.1522960 |
| MLGm009 | 3157171 | MLG | 3.35 | 3.35 | 3.33 | 0.0067730 | 0.0000000 |
| MLGm012 | 3134956 | MLG | 3.30 | 3.30 | 3.29 | 5.9460000 | 0.0911140 |
| MLGm013 | 3484557 | MLG | 2.74 | 2.74 | 2.73 | 0.1393000 | 0.0021940 |
| MLGm014 | 2858870 | MLG | 3.66 | 3.66 | 3.66 | 4.4670000 | 0.0841600 |
| MLGm016 | 2214552 | MLG | 3.83 | 3.83 | 3.82 | 4.4960000 | 0.0690170 |
| MLGm017 | 3695528 | MLG | 3.31 | 3.31 | 3.29 | 6.1700000 | 0.1152330 |
| MMSf206 | 3707087 | MMS | 4.55 | 4.55 | 4.51 | 0.0484500 | 0.0000000 |
| MMSf211 | 2202122 | MMS | 7.40 | 7.40 | 7.36 | 0.0252100 | 0.0000000 |
| MMSf215 | 3840292 | MMS | 3.75 | 3.75 | 3.73 | 0.0136600 | 0.0000000 |
| MMSf217 | 2707625 | MMS | 6.44 | 6.44 | 6.40 | 0.0817800 | 0.0000000 |
| MMSf221 | 4208537 | MMS | 4.26 | 4.26 | 4.23 | 0.0153000 | 0.0000000 |
| MMSf222 | 3952532 | MMS | 4.84 | 4.84 | 4.79 | 0.1184000 | 0.0000000 |
| MMSm207 | 3517181 | MMS | 4.52 | 4.52 | 4.49 | 0.0097640 | 0.0000000 |

Table S4: Summary of Reads mapping toWolbachia *(continued)*

| **Sample** | **Total Reads** | **Population** | **% Total Reads Classified by Centrifuge** | **% Total Reads Classified as Microbial (Archeae and Bacteria)** | **% Total Reads Classified as Bacterial** | **% of Classified Reads mapping to Wolbachia wNo** | **% of Classified Reads mapping to other Wolbachia taxa** |
|---|---|---|---|---|---|---|---|
| MMSm208 | 6585205 | MMS | 4.21 | 4.21 | 4.18 | 0.0117000 | 0.0003776 |
| MMSm209 | 3346988 | MMS | 5.09 | 5.09 | 5.05 | 0.0073450 | 0.0000000 |
| MMSm210 | 3233531 | MMS | 5.83 | 5.83 | 5.81 | 0.0044030 | 0.0000000 |
| MMSm212 | 3408275 | MMS | 4.63 | 4.63 | 4.58 | 0.0073910 | 0.0006719 |
| MMSm213 | 2318554 | MMS | 6.87 | 6.87 | 6.83 | 0.0019820 | 0.0000000 |
| MMSm214 | 3178148 | MMS | 5.55 | 5.55 | 5.50 | 0.0053980 | 0.0000000 |
| MMSm220 | 4093188 | MMS | 4.51 | 4.51 | 4.48 | 0.0079450 | 0.0000000 |
| OBNf111 | 2630822 | OBN | 3.09 | 3.09 | 3.08 | 0.0737100 | 0.0000000 |
| OBNf121 | 2690338 | OBN | 3.92 | 3.92 | 3.91 | 7.1730000 | 0.1592340 |
| OBNm110 | 2681431 | OBN | 4.03 | 4.03 | 4.01 | 6.8400000 | 0.1763280 |
| OBNm112 | 2751978 | OBN | 3.52 | 3.52 | 3.51 | 7.8850000 | 0.1613200 |
| OBNm113 | 3194428 | OBN | 3.31 | 3.31 | 3.30 | 5.1820000 | 0.0582510 |
| OBNm114 | 227420 | OBN | 9.96 | 9.96 | 9.95 | 2.0440000 | 0.0882720 |
| OBNm115 | 2830379 | OBN | 3.21 | 3.21 | 3.20 | 0.0188400 | 0.0000000 |
| OBNm116 | 3026011 | OBN | 3.84 | 3.84 | 3.82 | 3.3610000 | 0.0687910 |
| OBNm117 | 2249328 | OBN | 4.09 | 4.09 | 4.07 | 5.2970000 | 0.1056250 |
| OBNm118 | 1753843 | OBN | 8.01 | 8.01 | 7.93 | 0.0175800 | 0.0000000 |
| OBNm119 | 2786199 | OBN | 3.96 | 3.96 | 3.95 | 0.0226700 | 0.0009444 |
| OBNm120 | 3188535 | OBN | 3.09 | 3.09 | 3.08 | 0.2973000 | 0.0000000 |
| OBNm122 | 1963010 | OBN | 4.32 | 4.32 | 4.30 | 5.5760000 | 0.0974170 |
| OBNm123 | 1680297 | OBN | 6.19 | 6.19 | 6.16 | 4.1080000 | 0.0782600 |
| OBNm124 | 754559 | OBN | 10.30 | 10.30 | 10.30 | 0.0121200 | 0.0000000 |
| PCPf161 | 1798002 | PCP | 7.89 | 7.89 | 7.86 | 0.0037070 | 0.0000000 |
| PCPf175 | 3534243 | PCP | 4.66 | 4.66 | 4.62 | 0.0262300 | 0.0000000 |
| PCPm156 | 3363460 | PCP | 5.61 | 5.61 | 5.56 | 0.0057320 | 0.0005732 |
| PCPm157 | 2928578 | PCP | 6.23 | 6.23 | 6.18 | 0.0035160 | 0.0005861 |
| PCPm158 | 3130749 | PCP | 6.42 | 6.42 | 6.36 | 0.0069110 | 0.0005316 |
| PCPm159 | 3436320 | PCP | 4.92 | 4.92 | 4.88 | 0.0037330 | 0.0000000 |
| PCPm160 | 2575990 | PCP | 6.46 | 6.46 | 6.42 | 0.0037580 | 0.0000000 |
| PCPm162 | 1826962 | PCP | 9.87 | 9.87 | 9.84 | 0.0023580 | 0.0000000 |
| PCPm172 | 2959900 | PCP | 5.01 | 5.01 | 4.98 | 0.0020970 | 0.0000000 |
| PCPm173 | 1373989 | PCP | 10.20 | 10.20 | 10.20 | 0.0029820 | 0.0000000 |
| PCPm174 | 781039 | PCP | 9.21 | 9.21 | 9.20 | 0.0014330 | 0.0000000 |
| PCPm176 | 1701200 | PCP | 6.86 | 6.86 | 6.83 | 0.0008968 | 0.0008968 |
| PCPm177 | 2015460 | PCP | 9.13 | 9.13 | 9.10 | 0.0023210 | 0.0000000 |
| RHDf242 | 3187266 | RHD | 4.44 | 4.44 | 4.42 | 0.0647600 | 0.0000000 |

Table S4: Summary of Reads mapping toWolbachia *(continued)*

| **Sample** | **Total Reads** | **Population** | **% Total Reads Classified by Centrifuge** | **% Total Reads Classified as Microbial (Archeae and Bacteria)** | **% Total Reads Classified as Bacterial** | **% of Classified Reads mapping to Wolbachia wNo** | **% of Classified Reads mapping to other Wolbachia taxa** |
|---|---|---|---|---|---|---|---|
| RHDf245 | 3282691 | RHD | 6.46 | 6.46 | 6.43 | 0.1333000 | 0.0010330 |
| RHDf253 | 1198067 | RHD | 8.33 | 8.33 | 8.32 | 0.0166300 | 0.0000000 |
| RHDm239 | 3816448 | RHD | 5.78 | 5.78 | 5.73 | 0.0043790 | 0.0000000 |
| RHDm240 | 3883290 | RHD | 4.90 | 4.90 | 4.87 | 0.0011090 | 0.0000000 |
| RHDm241 | 1863382 | RHD | 7.74 | 7.74 | 7.71 | 0.0021670 | 0.0000000 |
| RHDm246 | 1379829 | RHD | 8.43 | 8.43 | 8.41 | 0.0071050 | 0.0000000 |
| RHDm247 | 4254163 | RHD | 5.74 | 5.74 | 5.69 | 0.0039870 | 0.0000000 |
| RHDm248 | 3382304 | RHD | 4.92 | 4.92 | 4.89 | 0.0063290 | 0.0000000 |
| RHDm254 | 2413011 | RHD | 6.91 | 6.91 | 6.88 | 0.0025260 | 0.0000000 |
| RHDm255 | 2782584 | RHD | 6.73 | 6.73 | 6.71 | 0.0044750 | 0.0000000 |
| RHDm256 | 2032467 | RHD | 7.49 | 7.49 | 7.47 | 0.0068410 | 0.0000000 |
| RHDm257 | 4126879 | RHD | 5.50 | 5.50 | 5.45 | 5.0120000 | 0.2209100 |
| CFWm223 | 2672555 | CFW | 5.79 | 5.79 | 5.76 | 0.0033720 | 0.0000000 |
| CFWm224 | 931489 | CFW | 9.74 | 9.74 | 9.72 | 0.0011380 | 0.0000000 |
| CFWm225 | 184682 | CFW | 11.10 | 11.10 | 11.10 | NA | 0.0000000 |
| CFWm226 | 2763326 | CFW | 4.99 | 4.99 | 4.97 | 0.0030070 | 0.0000000 |
| CFWm227 | 585059 | CFW | 11.60 | 11.60 | 11.60 | 0.0060920 | 0.0000000 |
| CFWm228 | 2217151 | CFW | 7.42 | 7.42 | 7.39 | NA | 0.0000000 |
| CFWm229 | 2446267 | CFW | 7.31 | 7.31 | 7.28 | 0.0011660 | 0.0000000 |
| CFWm230 | 2313054 | CFW | 7.53 | 7.53 | 7.49 | 0.0030410 | 0.0000000 |
| CFWm231 | 2292118 | CFW | 6.36 | 6.36 | 6.33 | 0.0035850 | 0.0000000 |
| CFWm232 | 256147 | CFW | 11.30 | 11.30 | 11.30 | 0.0035690 | 0.0000000 |
| CFWm233 | 2173984 | CFW | 7.49 | 7.49 | 7.46 | 0.0102300 | 0.0000000 |
| CFWm234 | 2793732 | CFW | 6.07 | 6.07 | 6.04 | 0.0018580 | 0.0000000 |
| RVSfR3 | 3181026 | RVS | 2.76 | 2.76 | 2.75 | 0.5478000 | 0.0011650 |
| RVSfR4 | 2034084 | RVS | 3.97 | 3.97 | 3.97 | 0.0731900 | 0.0000000 |
| RVSmR1 | 3068970 | RVS | 3.33 | 3.33 | 3.32 | 0.0946300 | 0.0000000 |
| RVSmR2 | 2084868 | RVS | 3.79 | 3.79 | 3.77 | 0.1693000 | 0.0000000 |
| RVSmR5 | 2532622 | RVS | 2.96 | 2.96 | 2.95 | 0.0829200 | 0.0014060 |
| RVSmR6 | 2743057 | RVS | 2.76 | 2.76 | 2.75 | 0.0194300 | 0.0000000 |
| TULf054 | 3159299 | TUL | 3.24 | 3.24 | 3.21 | 15.6000000 | 0.2325030 |
| TULf061 | 2598097 | TUL | 5.52 | 5.52 | 5.51 | 38.6200000 | 0.2518600 |
| TULf064 | 3009782 | TUL | 3.48 | 3.48 | 3.46 | 53.5600000 | 0.4500400 |
| TULf072 | 3142924 | TUL | 6.09 | 6.09 | 6.08 | 42.4700000 | 0.3689300 |
| TULf073 | 3443965 | TUL | 5.02 | 5.02 | 5.01 | 40.9600000 | 0.3228800 |
| TULf075 | 3457076 | TUL | 5.27 | 5.27 | 5.26 | 46.9700000 | 0.4356400 |

Table S4: Summary of Reads mapping toWolbachia *(continued)*

| **Sample** | **Total Reads** | **Population** | **% Total Reads Classified by Centrifuge** | **% Total Reads Classified as Microbial (Archeae and Bacteria)** | **% Total Reads Classified as Bacterial** | **% of Classified Reads mapping to Wolbachia wNo** | **% of Classified Reads mapping to other Wolbachia taxa** |
|---|---|---|---|---|---|---|---|
| TULf077 | 3473548 | TUL | 4.68 | 4.68 | 4.67 | 42.8700000 | 0.2906059 |
| TULf078 | 3054081 | TUL | 5.51 | 5.51 | 5.46 | 42.6400000 | 0.4620500 |
| TULm053 | 1575217 | TUL | 5.55 | 5.55 | 5.54 | 11.5300000 | 0.1435170 |
| TULm055 | 1455895 | TUL | 5.94 | 5.94 | 5.93 | 5.4380000 | 0.0895540 |
| TULm060 | 3382266 | TUL | 3.64 | 3.64 | 3.63 | 15.2200000 | 0.2568900 |
| TULm062 | 3109015 | TUL | 3.73 | 3.73 | 3.72 | 19.1000000 | 0.2703500 |
| TULm063 | 3525244 | TUL | 2.99 | 2.99 | 2.97 | 0.0587700 | 0.0000000 |
| TULm069 | 2774500 | TUL | 2.63 | 2.63 | 2.59 | 0.0793400 | 0.0055680 |
| TULm070 | 2561524 | TUL | 4.95 | 4.95 | 4.93 | 19.8500000 | 0.2974900 |
| TULm071 | 4247179 | TUL | 3.48 | 3.48 | 3.46 | 6.5080000 | 0.1192360 |

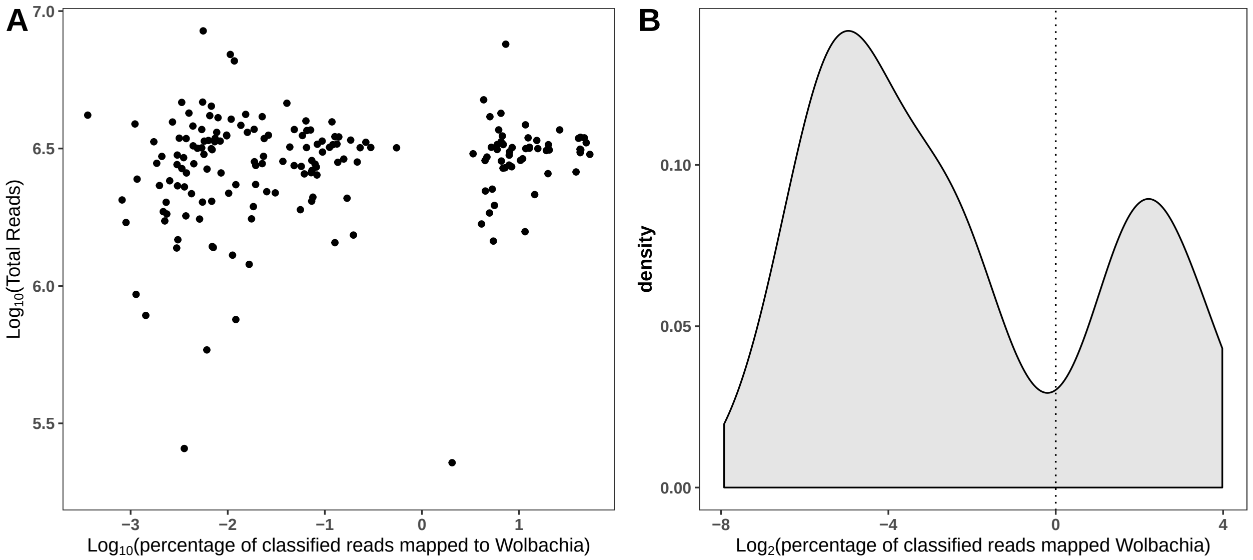


**Figure S9 (A)** Percentage of reads mapped to *Wolbachia* is independent of total read depth. **(B)** There was a natural discontinuity around $\log_2$(% reads mapped to *Wolbachia*) = 0 resulting in a bimodal distribution.

Table S5: Pairwise Fisher's Exact test for proportion infected by Wolbachia

| x | BER | DGC | MLG | OBN | RHD |
|---|---|---|---|---|---|
| **DGC** | 1 | NA | NA | NA | NA |
| **MLG** | 1 | 1 | NA | NA | NA |
| **OBN** | 1 | 1 | 1 | NA | NA |
| **RHD** | **0.02722** | **0.02009** | **0.02722** | 0.23602 | NA |
| **TUL** | 1 | 1 | 1 | 0.80703 | **0.00033** |

Table S6: Genotypes for female-specific sex RAD loci

| Marker* | Genotype Class | Phenotypic Sex | Uninfected | wIca1 | wIca2 |
|---|---|---|---|---|---|
| **9681__65** | **Homozygote** | **female** | 2 | 12 | 0 |
| | | **male** | 81 | 19 | 19 |
| | **Heterozygote** | **female** | 30 | 0 | 6 |
| | | **male** | 0 | 0 | 0 |
| **9781__55** | **Homozygote** | **female** | 0 | 11 | 0 |
| | | **male** | 70 | 17 | 17 |
| | **Heterozygote** | **female** | 26 | 0 | 4 |
| | | **male** | 0 | 0 | 0 |
| **11011__27** | **Homozygote** | **female** | 0 | 11 | 0 |
| | | **male** | 67 | 17 | 16 |
| | **Heterozygote** | **female** | 28 | 0 | 2 |
| | | **male** | 0 | 0 | 0 |
| **22073__63** | **Homozygote** | **female** | 2 | 10 | 0 |
| | | **male** | 68 | 19 | 20 |
| | **Heterozygote** | **female** | 21 | 1 | 2 |
| | | **male** | 0 | 0 | 0 |
| **22073__76** | **Homozygote** | **female** | 2 | 11 | 0 |
| | | **male** | 68 | 18 | 20 |
| | **Heterozygote** | **female** | 21 | 0 | 2 |
| | | **male** | 0 | 0 | 0 |
| **24861__17** | **Homozygote** | **female** | 0 | 9 | 0 |
| | | **male** | 66 | 15 | 19 |
| | **Heterozygote** | **female** | 27 | 0 | 2 |
| | | **male** | 0 | 0 | 0 |
| **24861__38** | **Homozygote** | **female** | 0 | 9 | 0 |
| | | **male** | 66 | 15 | 19 |
| | **Heterozygote** | **female** | 27 | 0 | 2 |
| | | **male** | 0 | 0 | 0 |
| **24861__43** | **Homozygote** | **female** | 0 | 9 | 0 |
| | | **male** | 66 | 15 | 19 |
| | **Heterozygote** | **female** | 26 | 0 | 2 |
| | | **male** | 0 | 0 | 0 |
| **25851__91** | **Homozygote** | **female** | 2 | 12 | 0 |
| | | **male** | 77 | 19 | 18 |
| | **Heterozygote** | **female** | 30 | 0 | 5 |
| | | **male** | 0 | 0 | 0 |
| **396673__60** | **Homozygote** | **female** | 7 | 11 | 2 |
| | | **male** | 70 | 15 | 17 |
| | **Heterozygote** | **female** | 14 | 0 | 2 |
| | | **male** | 0 | 0 | 0 |

* Prefix represents a unique RAD locus and the suffix represents position of a SNP on locus; 1 locus can have multiple SNPS
